# Selection for Decreased BRCA2 Functional Activity in *Homo sapiens* After Divergence from the Chimpanzee-Human Last Common Ancestor

**DOI:** 10.1101/2020.11.16.384677

**Authors:** Jinlong Huang, Yi Zhong, Alvin P. Makohon-Moore, Travis White, Maria Jasin, Mark A. Norell, Ward C. Wheeler, Christine A. Iacobuzio-Donahue

## Abstract

Humans have an increased incidence of epithelial neoplasia compared to non-human primates. We performed a comparative analysis of 21 non-human primate genomes and 54 ancient human genomes to identify variations in known cancer genes that may explain this difference. We identified 299 human-specific fixed non-silent single nucleotide polymorphisms. Bioinformatics analyses for functional consequences identified a number of variants predicted to have altered protein function, one of which was located at the most evolutionarily conserved domain of human BRCA2. This variant, in which a polar threonine residue replaces a hydrophobic methionine residue to codon 2662 within the DSS1 binding domain, decreases the interactions of BRCA2 with other proteins, specifically the binding of BRCA2 and RAD51, as well as the repairing ability of cells for DNA double-strand breaks. We conclude that a 20% reduction in BRCA2 DNA repair ability was positively selected for in the course of human evolution.

**One Sentence Summary:** Reduction of BRCA2 functional activity has been selected for during human evolution since the chimpanzee-human last common ancestor.

## Main Text

Cancer is a genetic disease^1–3^. Evidence gathered over the past two decades indicates that cancer arises from the activation of oncogenes and the inactivation of tumor suppressor genes. On average, somatic alterations of three to five cancer genes are required to generate an infiltrating neoplasm^1,4,5^. All known cancer driver genes can be classified into one or more of 12 pathways representing three core cellular processes: cell fate, cell survival, and genome maintenance. While this paradigm holds for describing cancers in general, malignancies arising in different tissue types show specific patterns with respect to the genes that are targeted within these core processes^1,2^. For example, oncogenic *KRAS* mutations result in a mutant KRAS protein that hyperactivates MAPK/ERK signaling in pancreatic, lung and colorectal adenocarcinomas; whereas inactivating mutations of the tumor suppressor gene *NF1* in melanoma or mesenchymal tumors, a negative regulator of the KRAS protein, has the same downstream consequence^6^. The diversity of tissue types in the human body together with the spectrum of genes targeted within a given signaling pathway have resulted in a consensus list of cancer genes currently recognized^7^.

Cancer is related to multicellularity^8^. Multicellularity requires that individual cells restrain their own growth potential for the benefit of the organism^9^; hence, the development of a neoplasm due to loss of growth control reflects the escape from this context^10^. Indeed, neoplasia has been described in a multitude of vertebrate and invertebrate metazoan species, both extant and extinct^11–16^. The natural prevalence of neoplasia in animal species in the wild is generally unknown as population-level analyses are difficult to perform. In those that have been assessed, viral etiologies, environmental pollutants or transmissible allografting play a role^17^. Available data from captive and domestic animals indicate a modest, yet significant, rate of benign and malignant neoplasm formation although the incidence and spectra of neoplasms are different from that of humans^11,14,18^. Thus, while the propensity to develop neoplasms is a common feature of multicellular animals, cellular complexity and life span alone do not account for the remarkably high cancer rates seen in *Homo sapiens*.

*Pan troglodytes* (chimpanzee) has > 98% genome sequence similarity with *Homo sapiens*^19^ but does not have a high rate of epithelial cancer^14,20^. Since divergence from the chimpanzee-human last common ancestor (CHLCA), *Homo sapiens* has evolved in association with a multitude of environmental, geographic, social and behavioral changes^21^. We hypothesized that a systematic analysis of cancer gene evolution between non-human primates and *Homo sapiens* would reveal mechanisms that increase cancer propensity in our species. Exploration of this hypothesis surpasses mere academic interest given that more than 20 million cases per year are expected by 2025 worldwide^18^.

## Results

### Construction of a variations database of primate whole genomes

We performed a systematic analysis of 21 non-human primate genomes and 54 ancient human genomes (Supplementary Table 1) to investigate variations in whole genomes during evolution from CHLCA (Fig. 1a). These 21 non-human primate genomes are from 12 groups^22,23^ (Supplementary Fig. 1). The ancient human genomes, defined as any human remains aged 2000 years bp (before present) or older, include five humans dated 10,000–100,000 years bp and 49 dated 2,000–10,000 years bp^24^ (Supplementary Table 1). After mapping the sequence data to the human reference genome (GRCh38), we identify ~420 million reliable SNPs/Indels (hereafter referred to as SNPs) in non-human primate genomes and ancient human genomes compared to human reference genome. A large variation in the number of SNPs is observed among species, primarily due to evolutionary relationships and sequencing depth (Supplementary Table 2). Specifically, chimpanzee, our closest extant relative, has ~37 million SNPs compared to human reference genome; *Rhinopithecus* (an old world snub-nosed monkey) has the highest number of SNPs (~110 million) (Supplementary Fig. 2a). Ancient human genomes have a median of ~600K SNPs compared to the human reference genome (Supplementary Fig. 2b). Further annotation of all SNPs for the predicted consequences to their expressed proteins results in a dataset of ~1.6 million non-silent SNPs in one or more non-human primates and ancient human genomes compared to the human reference genome (Fig. 1b and c, Supplementary Table 2). A phylogenetic reconstruction using this dataset retains the accepted relationships among species, indicating a reliable result of SNP calling (Supplementary Fig. 3). This dataset forms the basis for further analyses.

**Fig. 1.**
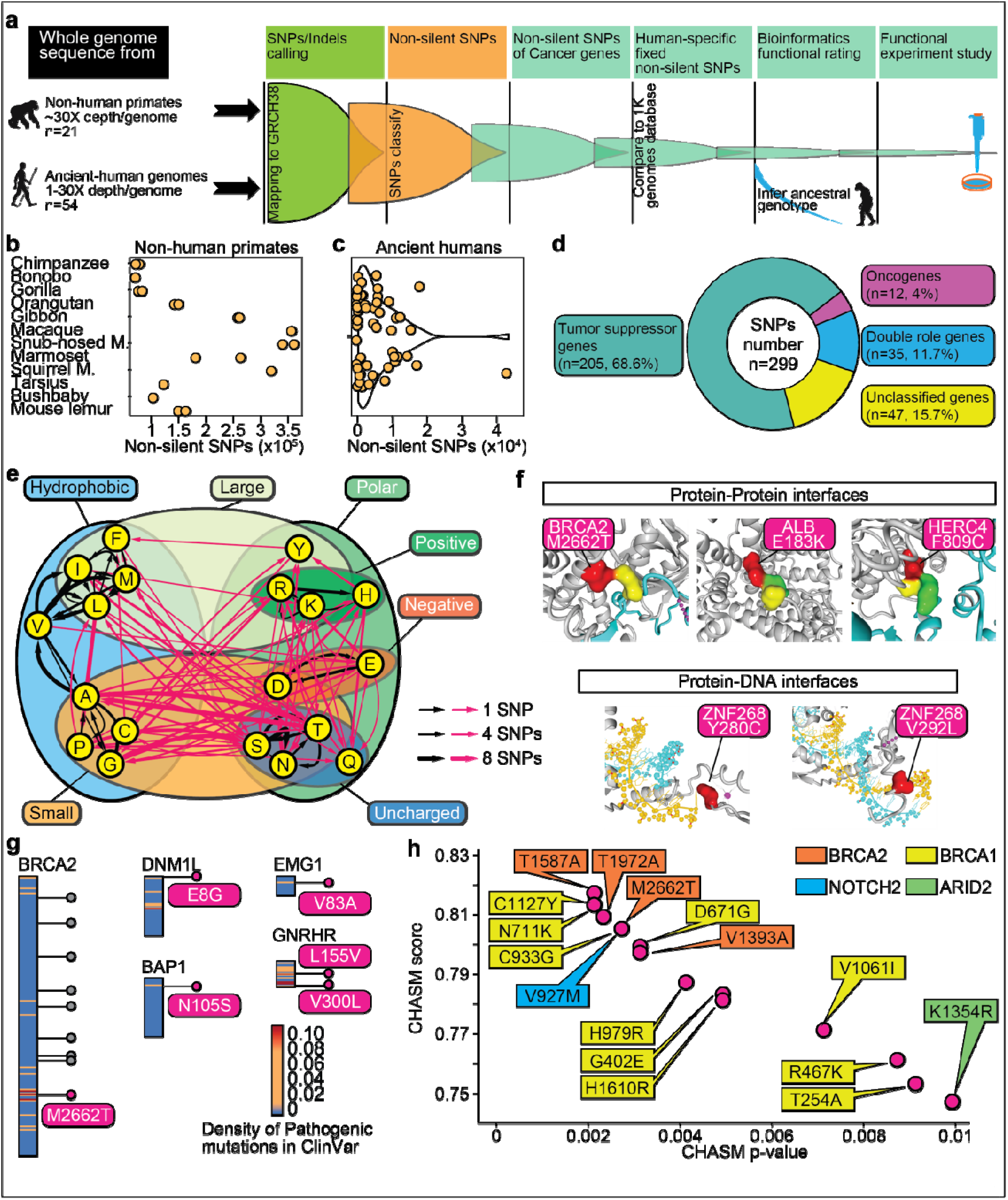
Determine potential functional consequences of human-specific fixed SNPs. **(a)** Schematic for study strategy of human-specific fixed SNPs of cancer genes. **(b)** Number of non-silent SNPs in 21 genomes of 12 non-human primate groups. **(c)** Number of non-silent SNPs in 54 ancient human genomes. **(d)** Distribution of 299 human-specific fixed SNPs in the four categories of cancer genes. **(e)** Changes in the properties of amino acids of 299 human-specific fixed SNPs. A Venn diagram shows the classification of residues based on properties in amino acids. The red arrows denote SNPs with changes of amino acids properties, while the black arrows denote SNPs without changes in amino acids properties. The width of the arrow is proportional to the number of SNPs. **(f)** Five SNPs located at protein-protein or protein-DNA interfaces. **(g)** A comparison between the human-specific fixed SNPs and high reliability “Pathogenic” mutations in the ClinVar database. The dots on the genes denote human-specific fixed SNPs. Each gene is divided into windows of 20 amino acids. The color bar indicates the density of “Pathogenic” mutations for each window. **(h)** CHASM scoring for 299 human-specific fixed SNPs. Sixteen SNPs with p-value smaller than 0.01 are shown.

### Evolutionary features of cancer genes

We next focus on the genetic evolution of 401 cancer genes representing the union of two different driver gene prediction methods (20/20+, TUSON) applied to >700,000 somatic mutations from 34 cancer types^7^ (Supplementary Table 3). These prediction algorithms classify these 401 cancer genes into one of four categories: oncogenes (n=78), tumor suppressor genes (n=211), double role genes (i.e. role is dependent on tumor type, n=20) and unclassified genes (n=92).

We first calculated the sequence similarities of all exons of these 401 cancer genes using the list of non-silent SNPs detected from the 21 non-human primate genomes and 54 ancient human genomes. We find that the exon sequence similarity of oncogenes is higher than that of all other classes of cancer gene whether using data of non-human primates (Supplementary Fig. 4a, Supplementary Table 4) or ancient humans (Supplementary Fig. 4b, Supplementary Table 4). This implies a more conserved functional evolution in oncogenes compared with other cancer gene types. To avoid biases caused by cross species mapping, we also used protein sequences derived from each species assembled genome sequence (Supplementary Fig. 4c, Supplementary Table 5). Similar to that found using SNPs, the protein sequence similarity of oncogenes is higher than that of other cancer gene types indicating that SNPs with functional effects on the protein product in oncogenes were less tolerated than those in other types of cancer genes.

dN/dS (a ratio that measures the relative rates of nonsynonymous and synonymous substitutions) was used as an indicator of selective pressure in protein coding genes^25^. We retrieved the dN and dS of protein-coding genes of the human reference genome comparing to other non-human primate species from the Ensembl database. We determined the dN/dS of the human reference genome for each category of cancer gene and compared these findings to the whole genome background (Supplementary Fig. 4d, Supplementary Table 6). Overall, the average dN/dS of the background genome (0.20-0.35) is higher than that of cancer genes. Among cancer genes specifically the average dN/dS of oncogenes and tumor suppressor genes was 0.08-0.18 and 0.16-0.24, respectively. This relationship holds when the human genome is specifically compared to seven other primate species, again implying stronger negative selection pressure in oncogenes compared to tumor suppressor genes. To further explore this characteristic, we performed the McDonald-Kreitman test^26^ to examine the extent that selection drives the differentiation of cancer genes. Using the SNPs of 88 great apes (nine present-day *Homo sapiens* and 79 non-human great apes) from whole genome sequencing^27^, we calculated the adaptive selection index^28^ for each gene type (Supplementary Table 7). As expected, the level of adaptive selection in the background genome is slightly less than 0 (p-value ≤0.0001, Supplementary Fig. 4e), indicating a minor negative baseline selection pressure. Interestingly, double role genes show extremely dynamic trajectories across species, while oncogenes and tumor suppressor genes were relatively stable. We again identify much stronger negative selection in oncogenes than in tumor suppressor genes.

### Emergent genetic variations during evolution of primates

We next investigated emergent genetic variations during evolution for each branch. Following divergence of humans and chimpanzees from the CHLCA, the human branch has accumulated 388 new variations in these 401 cancer genes (Supplementary Fig. 5a). Furthermore, present-day humans have accumulated six new variations (Supplementary Fig. 5a, b) in six cancer genes (*COL11A1, ZNF292, DCHS1, SCAF11, FANCM, ATRX*) compared to Neanderthal/Denisovan genomes that are the oldest ancient human genomes in this study. The rate of polymorphism of these six sites was determined in the present-day human population by comparison to that found in the 1000 Genomes Project database^29^ (n=2504), indicating that the preponderant genotypes of present-day humans are different from those of non-human primates and Neanderthal/Denisovan. Finally, we calculated the rate at which these mutations accumulated (Supplementary Fig. 5c). Following divergence of apes from old world monkeys ~29 million years ago, 337.5 and 317.3, new non-silent mutations have accumulated in the 401 cancer genes per million years on average. However, the mutation rate slowed during the subsequent evolution of each species so that a mutation rate of 40-115 non-silent mutations per million years for each branch of great apes and lesser apes is inferred. A mutation rate of 57.9 non-silent mutations per million years is inferred for the homo branch, which was similar to all other species.

To further understand the identified SNPs, we filtered the human-specific non-silent 394 SNPs by comparing them to the 1000 Genomes Project database to identify those variants that are fixed in the present-day human population. This analysis reveals 299 human-specific fixed SNPs in 123 cancer genes (Fig. 1d). Tumor suppressor genes were the most common type of cancer gene to have at least one fixed SNP (61.8% of cancer genes) (Supplementary Fig. 5d), and fixed SNPs were most commonly observed in tumor suppressor genes (68.6% of SNPs) (Fig. 1d).

### Determine potential functional consequences of human-specific fixed SNPs

We applied four distinct methods to determine the putative functional effects of these 299 human-specific fixed cancer gene variants (Supplementary Fig. 6). We first examined those variants that involved changes to the properties of amino acid residues. There are 162 SNPs which account for changes in hydrophobicity, charge or molecule size (Fig. 1e, Supplementary Table 8). Next, we applied Mechismo^30^ to determine the predicted changes to the interactome of each protein. We detect three variants (BRCA2^M2662T^, ALB^E183K^, HERC4^F809C^) located at protein-protein interfaces and two variants in the transcriptional repressor ZNF268 (ZNF268^Y280C^ and ZNF268^V292L^) located at protein-DNA interfaces (Fig. 1f). BRCA2 is a central mediator of DNA damage response^31^, and BRCA2^M2662T^ is located within the DSS1 binding domain where it changes the hydrophobicity of the amino acid residue. *ALB* encodes the serum protein albumin^32^, and ALB^E183K^ lies at the interface of albumin dimers and changes the charge of amino acid residue. *HERC4* encodes a ubiquitin ligase^33^, and HERC4^F809C^ lies at the interface between HERC4-HERC4 trimers and changes the size of the residue. ZNF268 is a DNA binding factor and transcriptional repressor^34^, and both ZNF268^Y280C^ and ZNF268^V292L^ lie at the interface between ZNF268 and DNA where ZNF268^Y280C^ alters the hydrophobicity and size of the amino acid residue. We performed comparisons between the 299 human-specific fixed SNPs with known “Pathogenic” mutations in ClinVar^35^. We searched for windows with high densities of “Pathogenic” mutations in ClinVar for each cancer gene. This reveals six human-specific fixed SNPs overlap these windows: BRCA2^M2662T^, DNM1L^E8G^, BAP1^N105S^, EMG1^V83A^, GNRHR^L155V^ and GNRHR^V300L^ (Fig. 1g, Supplementary Table 9). Finally, as an alternative approach to determine potential functional consequences of the 299 human-specific fixed SNPs, we applied CHASM^36^ to determine predicted changes and retained these variations with a p-value less than 0.01. Sixteen SNPs of four genes met this requirement (Fig. 1h). The majority of these variants are in *BRCA2* (n=4) and *BRCA1* (n=10). One SNP each was found in *ARID2* and *NOTCH2*, respectively.

### Evolution pattern of BRCA2 codon 2662

The 299 fixed variants in known cancer genes have accumulated since divergence from the CHLCA. We believe these variations emerged gradually during human evolution, not simultaneously. Although our ability to distinguish the linkage relationship of these variations is currently limited, we can begin to understand cancer risk with respect to human evolution by studying a single mutation that represents a single evolutionary event^37^. We thus focused more closely on the BRCA2^M2662T^ variant because it is the only SNP identified by all four of the above methods as having functional consequences of potential significance in relation to the incidence of cancer in humans. Mutations in *BRCA2* are one of most common causes of inherited cancer risk^38^. Certain variants of the BRCA2 gene increase risk for breast cancer, ovarian cancer and some other cancers^39^. BRCA2 functions in homologous recombinational repair of DNA double-strand breaks (DSB) and does so in part through the actions of a carboxy-proximal region that bind DNA and several other proteins, including DSS1^40^. DSS1 promotes BRCA2-dependent homologous recombination, and BRCA2 mutants with impaired DSS1 binding display reduced activity for homologous recombination^41,42^.

Comparison of the whole protein sequences of BRCA2 of 56 osteichthyan species identified the most conserved domain ranges from codon 2603 to 2688 (Supplementary Fig. 7, Supplementary Table 10). This domain closely interacts with the N-terminal of DSS1. Nine (47.4%) of 19 known “Pathogenic" mutations in ClinVar cluster in this small domain (Supplementary Fig. 8). Furthermore, among the nine BRCA2 SNPs identified in this study BRCA2^M2662T^ is the sole variant present within the BRCA2-DSS1 domain (Fig. 2a, Supplementary Fig. 9).

**Fig. 2.**
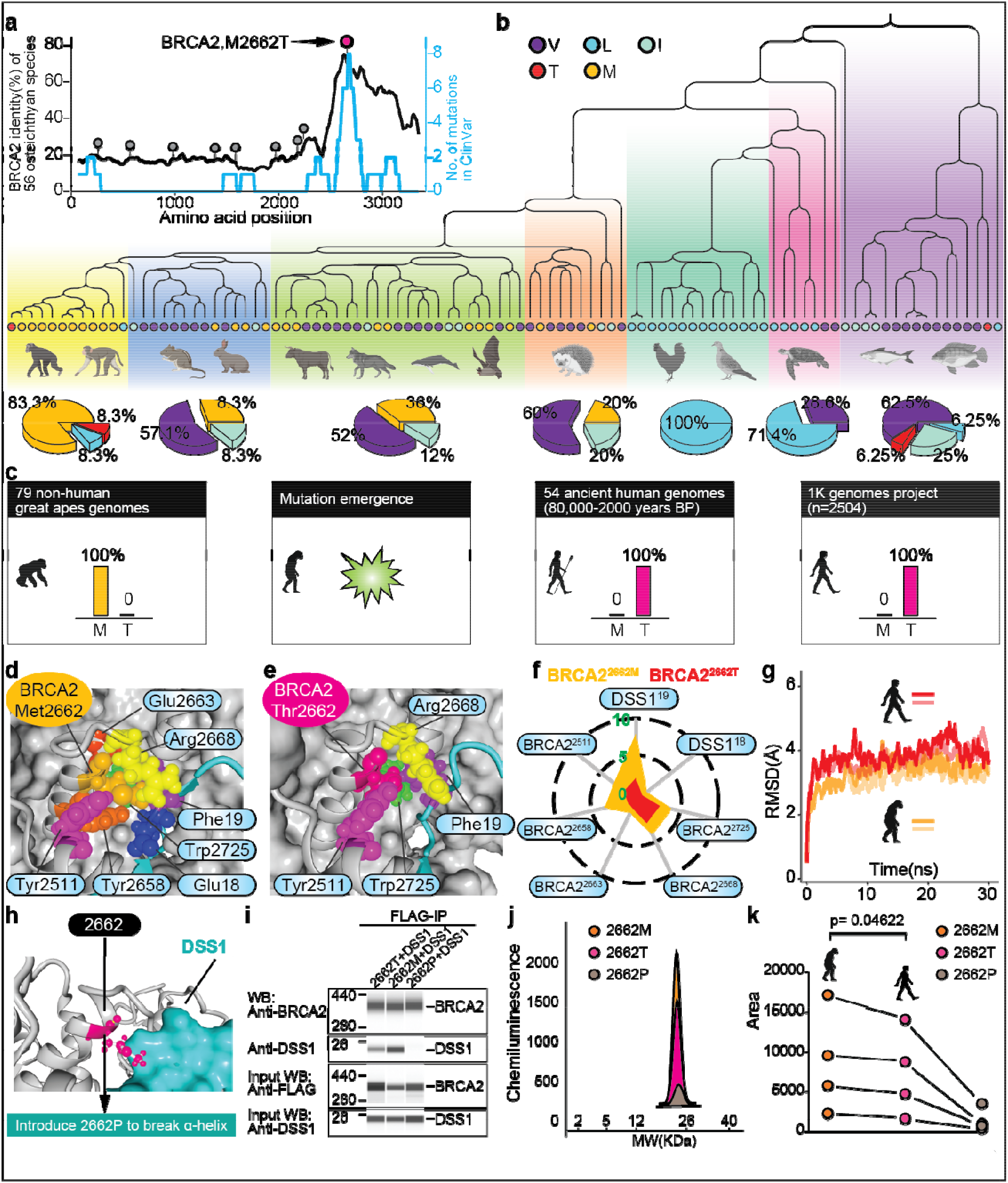
Evolution pattern of BRCA2^2662^, especially BRCA2^M2662T^ decreases the interaction of BRCA2 with DSS1. (**a**) A comparison between BRCA2^M2662T^ and other 8 human-specific fixed SNPs of BRCA2. The black line denotes the identity of BRCA2 based on 56 osteichthyan species. The identity is estimated by using a sliding window (size=130aa). The blue line denotes the density of “Pathogenic” mutations of ClinVar. The density is estimated by using a sliding window (size=130aa). (**b**) Distribution of 5 types of amino acids in BRCA2 codon 2662 or its homologous position for 98 osteichthyan species. **(c)** Graphical sketch of emergence and fixture of BRCA2^M2662T^. **(d-e)** Homologous modeling of BRCA2-DSS1 complex with the two different amino acids (M and T) within the site of BRCA2 codon 2662. Residues interacting with BRCA2 codon 2662 were shown. **(f)** A radar chart showed hydrophobic strength contributed by BRCA2^2662M^ (yellow) and BRCA2^2662T^ (red). Seven residues (five of BRCA2 and two of DSS1) interact with BRCA2 codon 2662. (**g**) Root mean square deviation (RMSD) of molecular simulation for BRCA2^2662M^-DSS1 complex and BRCA2^2662T^-DSS1 complex. **(h)** Codon 2662 is at the a-helix of BRCA2. **(i)** Lane view of co-immunoprecipitation and western blotting analysis of DSS1 and BRCA2’s variants. **(j)** Graph view of quantitative signal intensity for the bound DSS1 in the BRCA2 variants-immunoprecipitated protein complex. **(k)** BRCA2^M2662T^ reduces but does not abolish BRCA2-DSS1 binding. The introduction of BRCA2^2662P^ virtually abolishes BRCA2-DSS1 binding.

We compared the BRCA2 protein sequence around codon 2662 of 98 osteichthyan species. We find five different amino acids in the position of human BRCA2 codon 2662 and its homologous position (Fig. 2b). Almost all species have a hydrophobic amino acid in this position (M, methionine; I, isoleucine; L, leucine; V, valine) with the exception of *Homo sapiens* (T, threonine) and *Astyanax mexicanus* (blind cave fish). M and V are the canonical amino acids at this site in mammals. M is the primary type of amino acid in primates. Examination of the 1000 Genomes Project database, ancient human genomes and the 79 great ape genomes^27^ confirm that T is fixed in the human genomes and M is fixed primitively in the non-human great apes (Fig. 2c).

### BRCA2^M2662T^ decreases the interaction of BRCA2 with DSS1

DSS1 is an evolutionarily conserved protein that can stabilize BRCA2^41^. We therefore performed homologous modeling for the BRCA2-DSS1 complex with the two different amino acids (M and T) within the site of BRCA2 codon 2662. BRCA2^2662M^ can interact with seven surrounding residues (Fig. 2d), whereas BRCA2^2662T^ interacts with only four residues (Fig. 2e). The reduced number of amino acid residues engaged results in a predicted decrease of hydrophobic strength (Fig. 2f); most notably BRCA2^M2662T^ greatly decreases the interaction between BRCA2^2662^ and DSS1^19F^ (Fig. 2f). To further test this, molecular dynamic simulation was performed to investigate the effect of BRCA2^2662M^ and BRCA2^2662T^ on the interaction of BRCA2 with DSS1. Four independent simulations of BRCA2-DSS1 complex were performed, each run for 30ns (Fig. 2g). From these data, we calculated the RMSD (Root Mean Square Deviation) based on molecular trajectory to measure the interaction of BRCA2 with DSS1 (Supplementary Table 11). This reveals that the BRCA2-DSS1 complex with the BRCA2/2662T residue has the highest RMSD. Based on this result we conclude that during human evolution from the CHLCA, fixation of a threonine at this site leads to decreased interaction of BRCA2 with DSS1.

To further examine whether the alteration of amino acid at code 2662 did affect BRCA2 forming protein complex with DSS1 in vivo, we performed co-immunoprecipitation for BRCA2 and DSS1. We introduced BRCA2^2662P^ as a strong control because Proline can break the α-helix of BRCA2 (Fig. 2h). We found that the bound DSS1 was detected in either BRCA2^2662M^ or BRCA2^2662T^ protein complexes (Fig. 2i). The amount of the bound DSS1 detected in BRCA2^2662M^ protein complexes was increased at an average level of 22.4% (p=0.04622) over than that of in BRCA2^2662T^ complexes (Fig. 2j, k, Supplementary Table 12). The formation of the BRCA2/DSS1 complexes was significantly suppressed by replacing the code 2662 with Proline. These results are consistent with the findings that BRCA2^2662M^ has a stronger interaction with DSS1 than that of BRCA2^2662T^ in molecular dynamics simulation.

### BRCA2^M2662T^ decreases the binding of BRCA2 and RAD51

We consider that this decreased interaction of BRCA2 with DSS1 could further affect RAD51 as BRCA2 both binds RAD51 and potentiates recombinational DNA repair by promoting assembly of RAD51 onto single-stranded DNA (ssDNA)^43^. To test this hypothesis, we knocked down *BRCA2* in the 293T cell line by using CRISPR/Cas9 and a gRNA to target the second exon of *BRCA2* (Fig. 3a) thereby creating two independent single cell clones called 293T_a75 and 293T_a88 (Supplementary Fig. 10). By western blot analysis these two clones have lower expression of BRCA2 than wild type 293T cells (Fig. 3b), therefore they were used for subsequent two-hybrid tests to analyze the effect of various mutations of BRCA2 on the binding capacity of BRCA2 to RAD51 (Supplementary Figs. 11 and 12). We introduced 4 hydrophobic amino acids (M, V, L, I), 3 polar uncharged amino acids (T, N, S) and 3 charged amino acids (R, K, E) at the site of BRCA2 codon 2662. In addition, we introduced BRCA2^2723H^, BRCA2^2725A^ (Fig. 3c, in the context of BRCA2^2662T^) and BRCA2^2662P^ as controls. BRCA2^D2723H^ and BRCA2^W2725A^ are known deleterious mutations and are spatially adjacent to BRCA2 codon 2662 (Fig. 3c). Moreover, BRCA2^D2723H^ is a frequent mutation recorded in the Breast Cancer Information Core database^44^. The relative luciferase activity (the ratio of Firefly luciferase activity to *Renilla* luciferase activity) of BRCA2-RAD51 two-hybrid is used to measure BRCA2-RAD51 interaction (Supplementary Figs. 13-15). The relative luciferase activity of ten mutations at the site of BRCA2 codon 2662 were compared in pairs (Fig. 3d). In general, the relative luciferase activity of hydrophobic amino acids is greater than polar uncharged amino acids. Furthermore, the relative luciferase activity of polar uncharged amino acids is greater than charged amino acids. Collectively the M substitution and V substitution correspond to the highest BRCA2-RAD51 interaction, whereas BRCA2^2723H^ and BRCA2^2725A^ virtually abolishes the binding of BRCA2 and RAD51 (Fig. 3e, Supplementary Figs. 13 and 14). As expected, BRCA2^2662P^ can also abolish BRCA2-RAD51 interaction supporting that codon 2662 plays a key role for BRCA2 activity. We also find that the effect of BRCA2^M2662T^ is consistent with BRCA2^D2723H^, the only difference being that BRCA2^D2723H^ abolishes BRCA2-RAD51 interaction whereas BRCA2^M2662T^ reduces but does not eliminate BRCA2 activity (Fig. 3e). On average, BRCA2-RAD51 interaction is decreased by ~20% from the ancestral version (BRCA2^2662M^) to the human version (BRCA2^2662T^).

**Fig. 3.**
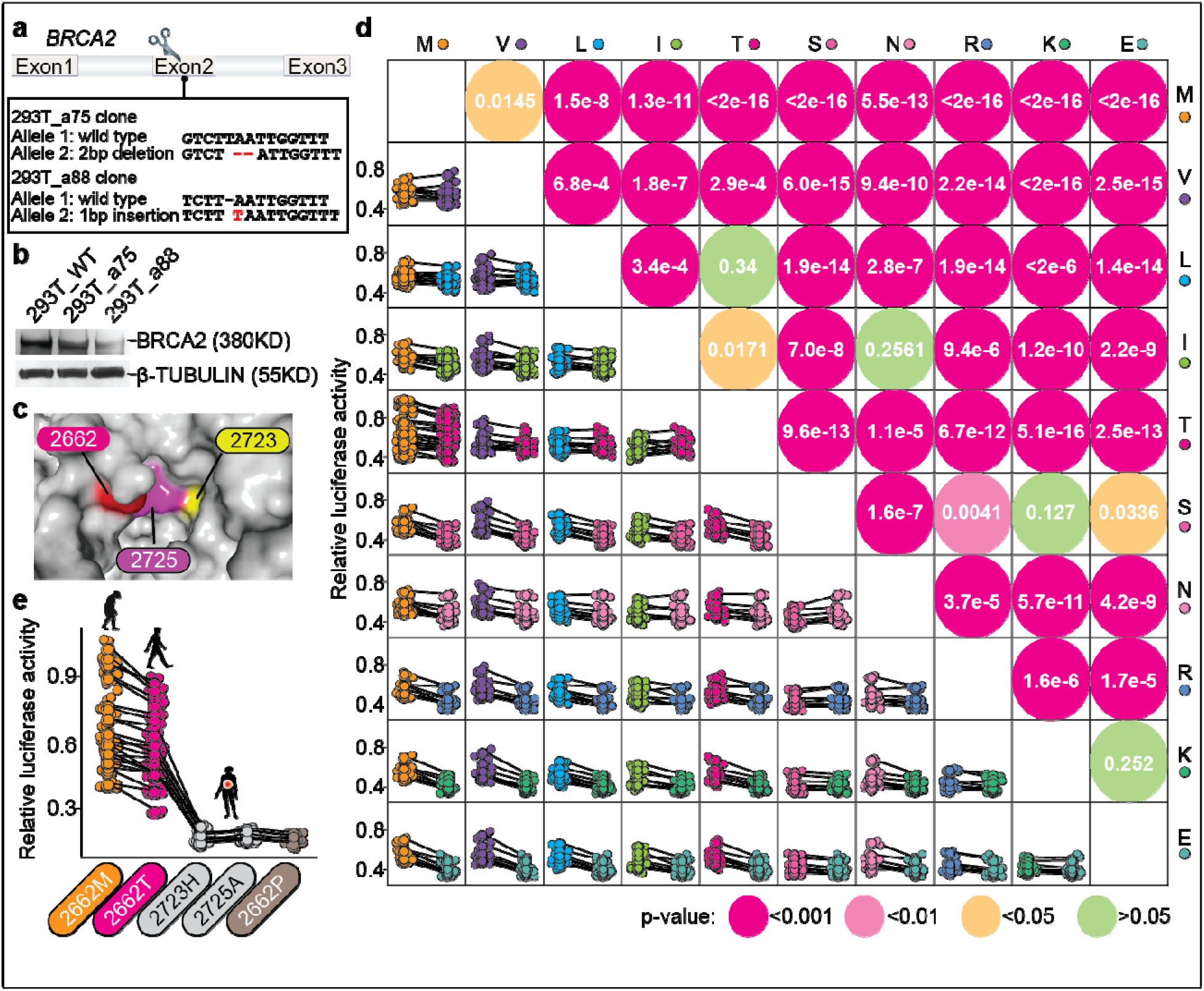
Different amino acids within the site of BRCA2 codon 2662 affect the binding of BRCA2 and RAD51, including BRCA2^M2662T^. (**a**) Edit *BRCA2* in 293T cell line by using CRISPR/Cas9. The sequence of 293T_a75, 293T_a88 clones were shown. **(b)** Western blot analysis showed decreased BRCA2 expression in 293T_a75 clone and 293T_a75 clone a compared with 293T wild-type cells. **(c)** Crystal structure showed BRCA2 codon 2662 is glued to codon 2725 and 2723. **(d)** Result of two-hybrid test for BRCA2 and RAD51. In the lower left, relative fluorescence intensity was shown. Scatter plots showed comparisons of each pair of amino acids. P-value of statistical test were also shown in the upper right. **(e)** BRCA2^M2662^ reduces but does not abolish BRCA2-RAD51 binding. The introduction of BRCA2^2662P^ virtually abolishes BRCA2-RAD51 binding.

We correlate BRCA2 activity with the frequency of each amino acid in 98 diverse animal species (Supplementary Fig. 16). Overall the frequency of M and V in these species is 65.3%. Among 62 mammalian species specifically, the frequency of M and V is as high as 85.2% (Fig. 2b, Supplementary Fig. 16). This raises the hypothesis that many species have a positive preference for highly active BRCA2. Yet humans appear to violate this rule in their recent evolution.

### BRCA2^M2662T^ decreased the DSB repair ability of cells

Since the predominant known role of BRCA2 is repair of DSB, we determined the extent to which BRCA2^M2662T^ affects DSB repair ability. Here we use mini-BRCA2^42^ (Fig. 4a) and V-C8 cells with DR-GFP reporter (Fig. 4b) to assess the effects of each variant. The mini-BRCA2 encompasses a human DNA binding domain (DBD) which can bind to DNA and DSS1 (Fig. 4a). We introduced two mutations in the DBD region which are homologous to BRCA2^2662M^ and BRCA2^2723H^ (in the context of BRCA2^2662T^). V-C8 cells were cotransfected with expression vectors for the I-SceI endonuclease and the mini-BRCA2 protein. Flow cytometry was used to quantify GFP-positive cells when three mutations of mini-BRCA2 occurred, respectively (Fig. 4c-e). Results from 4 independent transfection experiments (Fig. 4f, Supplementary Figs. 17-20) show an unequivocal pattern: the percentage of GFP positive cells significantly decreases by ~20% from the primitive ancestral version (BRCA2^2662M^) to the human version (BRCA2^2662T^). The magnitude of the effect of BRCA2^M2662T^ is also consistent with the known pathological mutation BRCA2^D2723H^. These experimental results, together with our models, suggest that during evolution to human the BRCA2^M2662T^ variant led to a decreased stability of BRCA2-associated protein complexes, specifically the binding of BRCA2 and RAD51. This is a restraint of recombinational DNA repair.

**Fig. 4.**
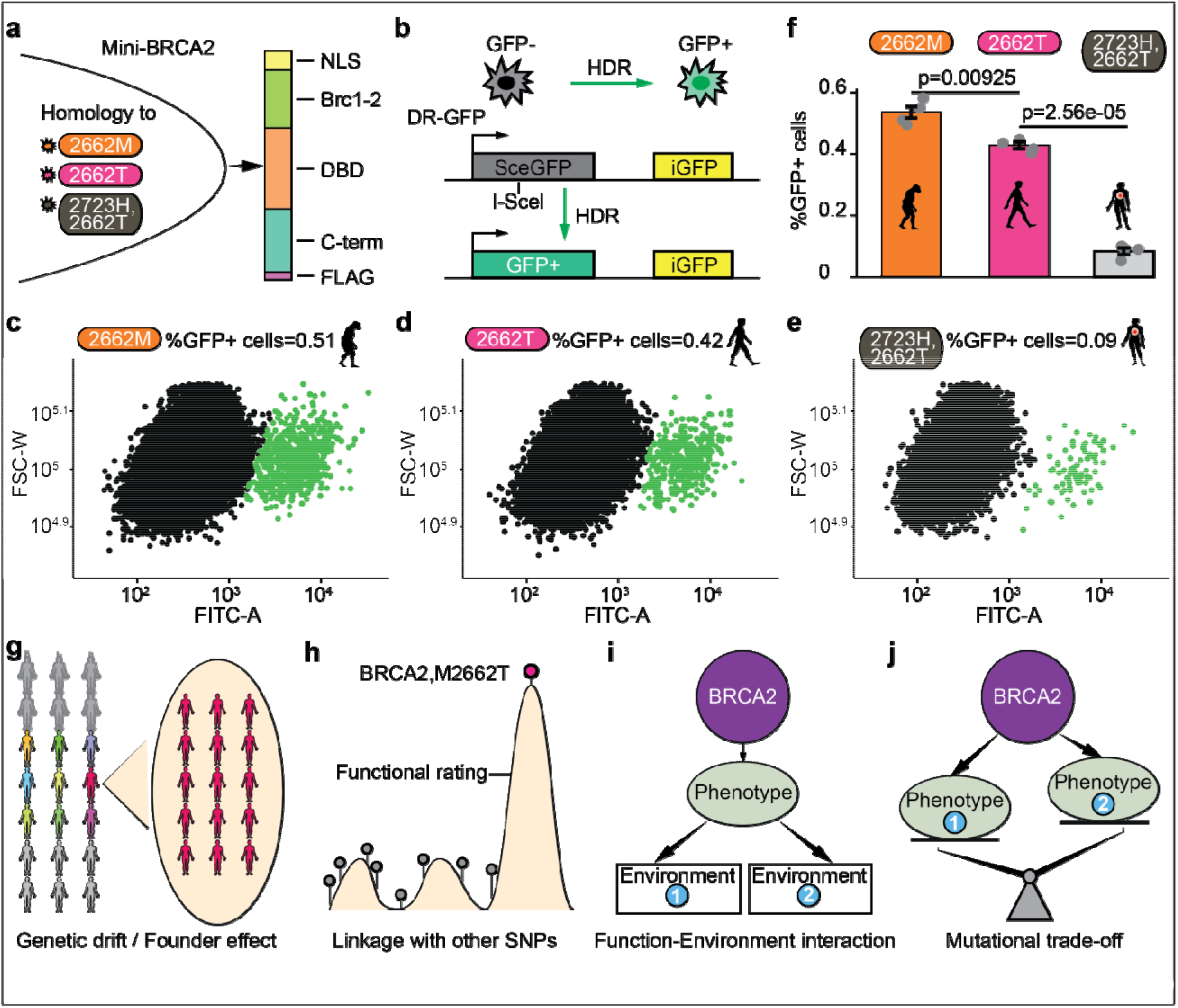
BRCA2^M2662T^ decreases the DSB repair ability of cells. **(a)** Mini-BRCA2 domain structures. Two mutations (BRCA2^2662M^ and BRCA2^2723H^) were introduced into DBD. **(b)** DR-GFP. The DR-GFP reporter consists of two defective GFP genes. Expression of I-SceI endonuclease results in a DSB at the I-SceI site in the SceGFP gene which can be repaired using the homologous sequence in the iGFP gene to generate GFP positive cells which are quantified by flow cytometry. **(c-e)** Representative flow cytometric analyses of V-C8 DR-GFP cells to detect cellular green fluorescence following DSB induction. **(f)** Comparison of DSB repair ability of mini-BRCA2 with different mutations. BRCA2^2662T^ significantly reduces the DSB repair ability of mini-BRCA2 when compared to BRCA2^2662M^. For BRCA2^2662T^ compared with BRCA2^2662M^, p-value = 0.00925; for BRCA2^2723H^ compared with BRCA2^2662T^, p-value = 2.56e-05. Error bars indicate standard deviation. **(g-j)** Four models to explain the emergence of BRCA2^M2662T^ in humans.

## Discussion

Our data provide comprehensive insights into several adaptive evolutionary features of cancer genes since divergence from the CHLCA. As previously reported, we find strong positive selection in genes within the non-homologous end joining (NHEJ) pathway^45–49^. Moreover, we have identified variants in cancer genes with unappreciated significance for a putative role in the human evolution such as *ALB*, *ZNF268, HERC4, EMG1, GNRHR or DNM1L*. While *BRCA2* specifically has been targeted during human evolution, we extend this observation showing that a single residue, T2662, has undergone positive selection and fixation in humans since the CHLCA. Moreover, we provide experimental evidence that positive selection for a polar amino acid at this site targets the DSS1 binding domain and consequently relaxes the DNA damage response by approximately ~20% compared to the ancestral methionine at this position.

Several evolutionary models are proposed to explain the occurrence of BRCA2^M2662T^. The first model is genetic drift/founder effect^50^ (Fig. 4g). However, one drawback is that genetic drift is often used to explain neutral variations^51^. Moreover, the founder effect does not explain why BRCA2^2662M^ and BRCA2^2662T^ are fixed in non-human primates and humans, respectively. A second model is linkage^51^ (Fig. 4h), i.e. other SNPs are linked to BRCA2/2662 which together confer more beneficial phenotypes. Our bioinformatics analyses for functional consequences indicate that there is no stronger candidate SNP than BRCA2/2662. However, our ability to distinguish the linkage relationship of these variations is currently limited and we did not assess structural changes in our dataset. In the third model, we assume the effect of BRCA2/M2662T depends on the interaction with environmental factors (Fig. 4i). That is to say, in a given environment, the reduction of BRCA2^M2662T^ DSB repair ability could confer higher fitness. However, it is hard to predict under what circumstances such a reversal would occur. The fourth model is a mutational trade-off^52,53^ (Fig. 4j). Here we suppose that BRCA2^M2662T^ exists as a trade-off where it enhances one aspect of performance while restraining another type of performance i.e. DSB repair. Although we cannot completely rule out the first three models, we favor the fourth given the multiple demonstrated roles of BRCA2. BRCA2 defects result in intrauterine growth retardation, microcephaly, and short thumb in humans^54,55^ and abnormal embryo development and kinked tails in mice^56^.

Lou et al.^47^ have suggested that NHEJ genes underwent rapid positive selection driven in part due to viruses that interact with NHEJ protein complexes during their lifecycle. Our findings do not dispute this possibility. However, we propose that additional selective pressure may have been to maintain both cellular and organismal/reproductive fitness in the context of social behaviors that increase DNA damage, for example the use of fire or the consumption of cooked meat 1.8-2.3 million years ago^57–59^. Of note, the importance of BRCA2/T2662 specifically is buttressed by our observation that a species of blind cavefish, *Astyanax mexicanus*, have independently acquired a threonine at the orthologous position compared to all other osteiciathians studied. While the functional consequences of a polar amino acid at this site likely differ among cavefish and humans, it is in keeping with a recent report that loss of function of mutations in DNA repair genes have contributed to the loss of photoreactivation DNA repair and evolution to blindness in *Phreatichthys andruzzii* ^60^.

While the genome has undergone neutral selection with respect to variants within the NHEJ pathway, we find that oncogenes as a class of cancer genes are much less tolerant of non-synonymous single nucleotide substitutions. This finding parallels fundamental features of oncogenes and tumor suppressor genes^1,3^. Thus, variants that change the protein structure of oncogenes may have provided little advantage during human evolution compared to those in tumor suppressor genes. These restraints on oncogene evolution do not appear limited to single nucleotide variants, as increasing gene size and proximity to tumor suppressor genes have also been implicated as a mechanism to limit oncogene somatic amplification^61^.

Our data indicate the power of merging evolutionary theory with cancer genomics to identify novel properties of cancer genes. This approach has the prospect to lay the ground work for a new integrative understanding of the biology and evolution of cancers and hopefully can be applied to other human health issues like neurodegenerative and cardiovascular diseases. Understanding the origins of these diseases should not start with their simple occurrence in humans but also use the powerful tools of contemporary evolutionary biology and systematics to understand their origins and their history in ours and related species.

## Materials and Methods

### Data collection

The sequencing data of 21 non-human primate genomes and 54 ancient human genomes and 2 present-day human genomes were downloaded from the SRA database^62^ (Sequence Read Archive, https://trace.ncbi.nlm.nih.gov/Traces/sra/) and ENA database (European Nucleotide Archive, https://www.ebi.ac.uk/ena) as “sra” format^27,63–82^. The detail information about these data including “Bioproject", “BioSample” and “Run ID” were shown in Supplementary Table 1. The total data amount is 3.6 Tb.

For non-human primate genomes, two genomes were available for chimpanzee (two of *Pan troglodytes*), bonobo (two of *Pan paniscus*), gorilla (two of *Gorilla gorilla*), orangutan (one each *Pongo abelii* and *Pongo pygmaeus*), gibbon (one each *Hylobates moloch* and *Symphalangus syndactylus*), macaque (two of *Macaca mulatta*), snub-nosed monkey (one each *Rhinopithecus strykeri* and *Rhinopithecus rolellana*), marmoset (two of *Callithrix jacchus*), and mouse lemur (two of *Microcebus murinus*), respectively. However, only one available genome was downloaded for squirrel monkey (*Saimiri boliviensis*), tarsius (*Carlito syrichta*) and bushbaby (*Otolemur garnettii*), respectively. The sequencing depth is about 30-fold for each genome.

For 54 ancient human genomes, the sequencing depth ranged from 1- to 30-fold coverage. Two representative ancient human genomes, Neanderthal and Denisovan, were included. The sequencing depth of Neanderthal and Denisovan is ~30-fold, respectively. Time estimation of ancient human genomes refer to Slatkin et al.^24^.

### Sequence quality checking and mapping

All analysis involving big data were performed with MSKCC’s high performance computing cluster. A filtering step was used to check sequence quality and remove low-quality reads. sratoolkit (version 2.6.3, https://www.ncbi.nlm.nih.gov/sra/docs/toolkitsoft/) were used to dump sequence reads from “sra” format to “fasta” format. bbmap (version 36.20, https://sourceforge.net/projects/bbmap/) was used to detect adapter sequences. If any adapter sequences were detected, fastx_toolkit (version 0.0.13, http://hannonlab.cshl.edu/fastx_toolkit/) fastx_clipper module was used to remove adapter sequences. Then fastx_toolkit fastq_quality_filter module was used to remove low-quality reads with parameter set “-q 30 -p 80 -Q 33”. At last FastQC (version 0.11.4, https://www.bioinformatics.babraham.ac.uk/projects/fastqc/) was used to double check to ensure that there is no adapter sequences and low-quality reads.

The human reference genome (GRCh38) was downloaded from the Ensembl database (ftp://ftp.ensembl.org/pub/release-84/fasta/homo_sapiens/dna/). First, the reference genome was indexed by using BWA^83^ (version 0.7.12, https://github.com/lh3/bwa). Then qualified sequence reads of 77 genomes were mapped to the indexed human reference genome using BWA. For 21 non-human primate genomes and 2 present-day human genomes, the parameter was options “-t 5 -q 15 -n 0.01”. For 54 ancient human genomes, the parameter was options “-t 5 -l 1024 -o 2 -n 0.02” which was described in Schubert et al.^84^. Other parameters were set to the default. This mapping process produces a battery of “sam” files.

After mapping, a series of samtools^85^ (version 1.3, https://github.com/samtools/samtools) orders, “view”, “sort”, “merge”, and “rmdup”, were used to convert file format, reorder reads, merge multiple bam files, and remove duplicate reads. We finally got a consolidated bam file for each genome to store all sequence information and used for subsequent analysis.

### SNPs/Indels calling and annotation

Firstly, samtools mpileup was used to transform bam file to mpileup file. Then SNPs and Indels were called using VarScan2^86^ (version 2.3.7, https://github.com/Jeltje/varscan2) according to the guidelines. For SNPs calling, the parameter was options “--min-coverage 2 --min-var-freq 0.20 --p-value 0.05”. For Indels calling, the parameter was options “--min-coverage 2 --min-var-freq 0.10 --p-value 0.10”. Other parameters were set to the default.

We performed an additional round of SNPs/Indels processing to filter on depth and quality. For Indels filtering, VarScan2 filter was used with parameter set “--min-reads 2 --min-var-freq 0.15 --p-value 0.05”. For SNPs filtering, Indels file of each genome was used as a filter to remove SNPs which are adjacent to Indels.

For SNPs/Indels (hereafter SNPs) annotation and effect prediction, SnpEff^87^ (version 4.2, https://github.com/pcingola/SnpEff) was used according to the guidelines. SNPs which lead to changes of proteins (for example, missense_variant, start_lost, stop_gained, and so on) were defined as non-silent SNPs in this study. Non-silent SNPs were identified for each genome.

### Construction of phylogenetic tree

Non-silent SNPs (n=1,561,780, Indels were excluded) of 2 old world monkeys (two of *Macaca mulatta*), 2 lesser apes (one each *Hylobates moloch* and *Symphalangus syndactylus*), 8 non-human great apes (two each of *Pan troglodytes*, *Pan paniscus*, *Gorilla gorilla*, and one each *Pongo abelii* and *Pongo pygmaeus*) and 54 ancient human genomes were used for this analysis. The phylogenetic analysis which is based on dynamic homologies were performed using POY5^88^ (version 5.0, https://github.com/amnh/poy5) in parallel on 40 cores of Intel Xeon Gold 6148 CPU at 2.4Ghz. The “search” option was used for 50 hours on each core for a total of 2000 CPU hours. A single most parsimonious tree of length 892,733 steps was found (showed in Supplementary Fig. 3a).

Another phylogenetic tree was inferred using the package APE^89^ (version 5.2, https://cran.r-project.org/web/packages/ape/) in R statistical environment (version 3.3.1, https://www.r-project.org/). The neighbor joining method was employed. The tree was bootstrapped by sampling the binary matrix of 1,561,780 non-silent SNPs 100 times. For visualization, FigTree (version 1.4.3, http://tree.bio.ed.ac.uk/software/figtree/) was used (showed in Supplementary Fig. 3b).

### Sequence similarity estimation by using mapping data

Genome annotation file (GTF, gene transfer format) of human reference genome was downloaded from Ensembl (ftp://ftp.ensembl.org/pub/release-84/gtf/homo_sapiens). Firstly, the longest transcripts were identified for each cancer genes^7^ (n=401). Then the exons corresponding to the longest transcripts were extracted. Thirdly, SNPs derived from mapping data were counted for each exon to estimate sequence similarity of exons.

### Sequence similarity estimation by using assembled protein sequences

Multiple alignment file of protein sequences was downloaded from UCSC (http://hgdownload.soe.ucsc.edu/goldenPath/hg38/multiz20way/). This file contains alignments of the protein sequence of the human reference genome (GRCh38) aligned to other 19 assemblies. From this multiple alignment file we extracted 12 non-human primate genomes (*Pan troglodytes*, *Pan paniscus*, *Gorilla gorilla*, *Pongo pygmaeus*, *Nomascus leucogenys*, *Macaca fascicularis*, *Rhinopithecus roxellana*, *Callithrix jacchus*, *Saimiri boliviensis*, *Tarsius syrichta*, *Otolemur garnettii*, *Microcebus murinus*). Then alignment corresponding to cancer genes and non-cancer genes were extracted, respectively. At last, SNPs and gaps of these alignment were counted for each exon to estimate sequence similarity.

### dN/dS of protein-coding genes

dN and dS values of human reference genome when compared with other 7 primate species were retrieved from Ensembl database by using BioMart (http://useast.ensembl.org/index.html). These 7 species are: *Pan troglodytes*, *Pongo abelii*, *Nomascus leucogenys*, *Macaca mulatta, Callithrix jacchus*, *Otolemur garnettii*, *Microcebus murinus*. The dataset “Ensembl Genes 84” were chosen and “Human Genes (GRCh38)” was used as reference. dN/dS was estimated for each gene. For visualization, Prism (version 7, https://www.graphpad.com/scientific-software/prism/) was used.

### McDonald-Kreitman test

Genome-wide data of 88 great apes were used in this analysis, including 9 present-day humans, 25 chimpanzees, 13 bonobos, 31 gorillas and 10 orangutans. The mean sequence depth was ~25-fold per individual. SNPs data were downloaded from the website of “Great Ape Genome Project”^27^ (http://biologiaevolutiva.org/greatape/data.html).

PopGenome^90^ (version 2.6.1, https://cran.r-project.org/web/packages/PopGenome/) package in R statistical environment was employed for McDonald-Kreitman test. Px (the number of non-synonymous polymorphisms), Ps (the number of synonymous polymorphisms), Dx (the number of non-synonymous substitutions), Ds (the number of synonymous substitutions) were determined in each test. We calculated adaptive selection index (AS) to better visualize the selection on different categories of genes. AS is calculated as AS=log_2_((Dx*Ps)/(Ds*Px)) as described in Li et al.^28^. In general, AS > 0 denote positive selection, AS < 0 denote opposite selection, and AS = 0 denote neutral selection. Significance levels were determined using one-sided Fisher exact test (https://www.socscistatistics.com/tests/Default.aspx).

### Emergent genetic variations for each branch and estimation of mutation rate

Only non-silent SNPs of cancer genes were counted in this analysis. We focus on the emergent SNPs of great apes (including human branch), lesser apes and old world monkeys because they are more closely related to human not hence as susceptible to bias caused by cross species mapping. Reference genome (GRCh38) and two additional genomes (SAMN00801888 and SAMN00001584, Supplementary Table 1) were used to represent present-day human genomes. Emergent SNPs were defined as SNPs that only found in a node and its child node, and not found in its parent node or sibling nodes.

In particular, for the 6 emergent SNPs between present-day humans and Neanderthal/ Denisovan, we estimated genotype frequencies in 8 non-human great ape genomes (two each of *Pan troglodytes*, *Pan paniscus*, *Gorilla gorilla*, and one each *Pongo abelii* and *Pongo pygmaeus*), 54 ancient human genomes and 2504 present-day human genomes. The genotype frequencies of 2504 present-day human genomes were retrieved from 1000 Genomes Project database (http://www.internationalgenome.org/) in combination with manual review.

Calibration times were queried from the TimeTree database^91^ (http://www.timetree.org) and showed in Supplementary Fig. 1. Mutation rate were defined as the number of emergent SNPs divided by divergence time. It is worth noting that the estimation of mutation rate is not suitable for present-day human and Neanderthal/Denisovan because there is no available divergence time for them in TimeTree database.

### Human-specific fixed SNPs filtered by 1000 Genomes Project database

CRAVAT^92^ (version 5.2.4, http://www.cravat.us/CRAVAT/) has a built-in database of 1000 Genomes Project database. Three hundred and ninety-four emergent SNPs were used in this analysis. The “vcf” file of SNPs was transformed to CRAVAT-format as described in CRAVAT guideline (http://www.cravat.us/CRAVAT/help.jsp). Genotype frequencies were queried and double checked manually. Only fixed SNPs (n=299) were retained after this analysis. These 299 SNPs were further compared with official SNPs file of 1000 Genomes Project database (ftp://ftp.1000genomes.ebi.ac.uk/vol1/ftp/release/20130502/) to ensure that all sites are fixed in present-day human population.

### Amino acid and Mechismo analysis

Classes of the 20 common amino acids refer to Pommie et al.^93^. The two “hydropathy” classes (hydrophobic, polar), three “volume” classes (large, medium, small) and three “charge” classes (positive, negative, uncharged) were used in this analysis. Detailed classification for each amino acid was showed in Fig. 1e.

We used Mechismo^30^ (http://mechismo.russelllab.org/) to identify SNPs which located in protein-protein or protein-DNA interactions. A list included 299 human-specific fixed SNPs were submit to Mechismo. “Low stringency” model (the default) was used. Five SNPs which has high confidence predictions were retained.

### Comparing to “Pathogenic” mutations in ClinVar

Variant summary file was download from ClinVar database^35^ (ftp://ftp.ncbi.nlm.nih.gov/pub/clinvar/tab_delimited/variant_summary.txt.gz). Indels and truncation SNPs were first removed, only those SNPs that result in amino acid substitution were retained. Then according to “clinical significance value”, SNPs with “Pathogenic” record were retained. Finally, the SNPs in cancer genes^7^ were extracted for further analysis. This filter step lead to a list including 327 known “Pathogenic” SNPs in 35 cancer genes.

We next sought to identify regions enriched for known “Pathogenic” SNPs across cancer genes. The average density of known “Pathogenic” SNPs was calculated with sliding windows of 20aa that had 90% overlap between adjacent windows. Further analysis shows that 6 human-specific fixed SNPs located in windows which has at least one ClinVar “Pathogenic” SNPs. This analysis is performed in R statistical environment. Package gtrellis^94^ (version 1.14.0, https://github.com/jokergoo/gtrellis) was used for visualizing density of “Pathogenic” SNPs.

### CHASM scoring

CHASM^95^ (version 3.1, http://www.cravat.us/CRAVAT/), which is a function module of CRAVAT, was used to identify and prioritize those 299 human-specific fixed SNPs most likely to generate functional changes of cancer genes. A list of 299 human-specific fixed SNPs was submitted to CHASM in the CRAVAT-format. Here, we chose “other” in the CHASM panel which represents the “pan-cancer” classifiers. CHASM score and p-value were queried and double checked manually. We required the variant to have a CHASM p-value < 0.01.

### Alignment of BRCA2-DSS1 domain around BRCA2/2662 of 98 osteichthyan species

Multiple alignment file of protein sequences of 98 osteichthyan species was downloaded from UCSC (http://hgdownload.soe.ucsc.edu/goldenPath/hg38/multiz100way/). Whole sequence of DSS1 and a 70aa fragment of BRCA2 that covered BRCA2/2662 were extracted for this analysis. According to the classification of UCSC, there are 12 primate species, 14 euarchontoglires species, 25 laurasiatheria species, 6 afrotheria species, 4 other mammal species, 14 aves species, 8 sarcopterygii species and 15 fish species included in this analysis.

### Identity of BRCA2 of 56 osteichthyan species

Whole genome protein sequences of 67 species were downloaded from Ensembl (https://useast.ensembl.org/info/data/ftp/index.html) in the format of fasta. The human BRCA2 (ENSP00000369497) was used as reference. Protein sequences of all other species were blast (version 2.5.0, https://blast.ncbi.nlm.nih.gov/Blast.cgi) with human BRCA2 with parameter set “-evalue 1e-3 -outfmt 6 -num_alignments 1”. After manual examination, the hit that has the best identity and coverage across 56 osteichthyan species were retained. These 56 sequences were used to calculate sequence identity of BRCA2:

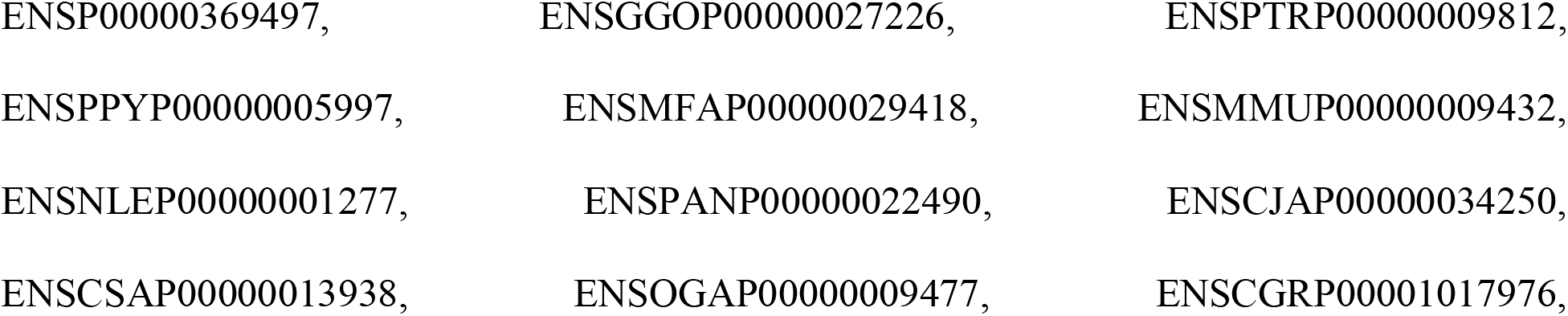
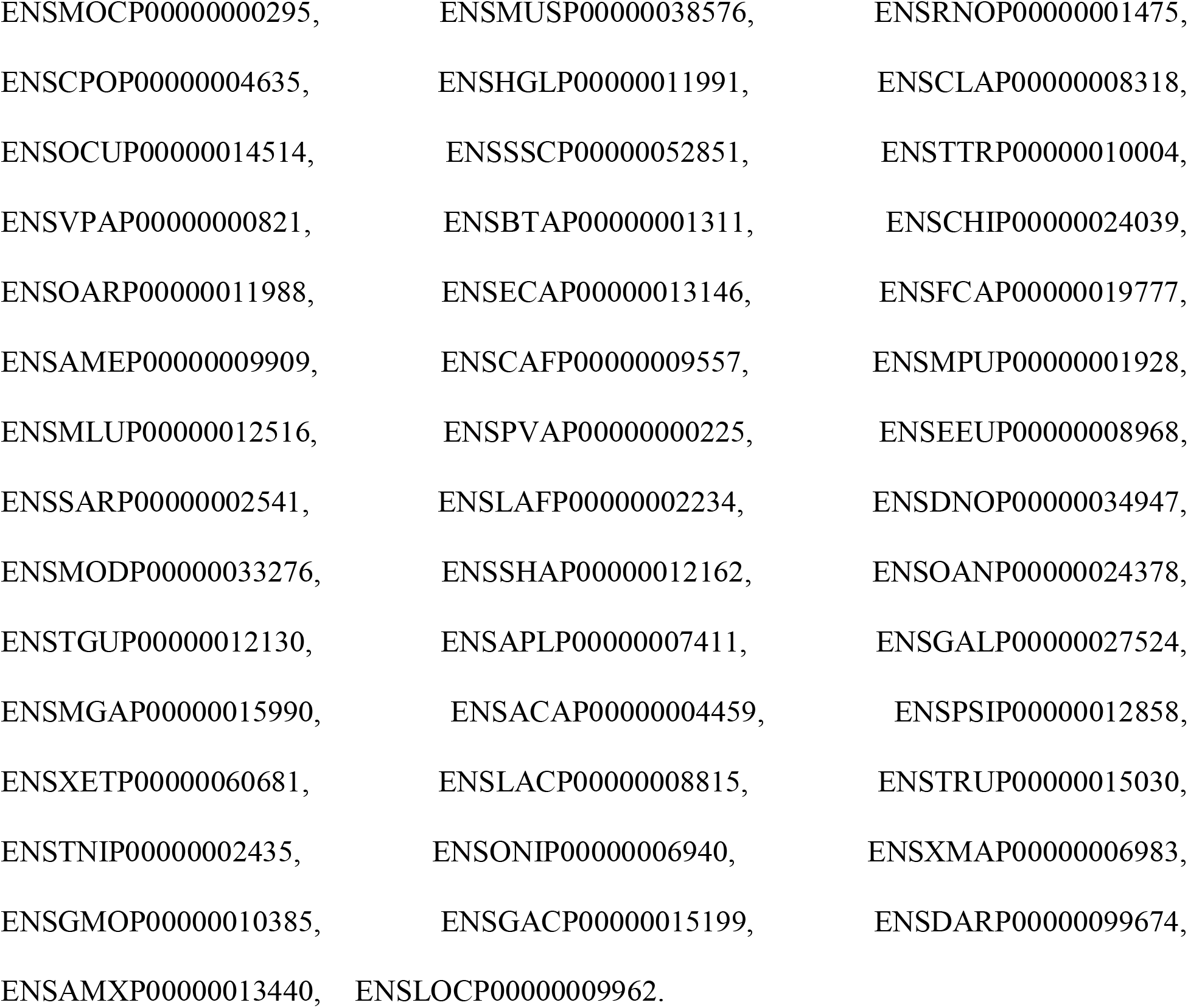

Multiple sequence alignment was performed by using ClustalX^96^ (version 2.1, http://www.clustal.org/clustal2/). Identity score were extracted using human BRCA2 as reference.

### Homologous modeling

The three-dimensional (3D) structure of the complex of BRCA2 (residues 2479-2818) and DSS1 (residues 6-23) were built by using YASARA Structure^97^ (version 11.3.2, http://www.yasara.org/). Homologous modeling was performed for the BRCA2-DSS1 complex with the two different amino acids (threonine; methionine) within the site of BRCA2 codon 2662. The “hm_build.mcr” macro of YASARA Structure with default parameters was used followed by an energy minimization process. Three protein structures were used as templates (PDB ID: 1MIU, 1IYJ, 1MJE). This step resulted in five models which were used for subsequent hydrophobic interaction analysis and molecular simulation.

The same modeling was also performed by using a larger complex of BRCA2 (residues 2373-3224) and DSS1 (residues 1-70). This model is used for structural visualization.

### Hydrophobic interaction analysis

In console mode of YASARA Structure, we used “ListIntRes” command to list codon 2662 and paired residues that are involved in hydrophobic interactions. The parameter was options “ListIntRes 2662, all, Type=Hydrophobic”. In simple terms, each hydrophobic atom pair contributes a value in the range [0..1] to the total interaction strength. Here, “0” denotes detectable interaction while “1” denotes optimal interaction. The total interaction strength is simply the sum of all the individual values.

### Molecular simulation

These two models resulted from homologous modeling were subjected to further refinement using the “md_refine.mcr” macro of YASARA Structure. We then performed molecular simulation for the two models. Using the standard macro “md_run.mcr”, YASARA Structure proceeded as follows: cleaning the structure, optimizing the hydrogen bonding network, creating a simulation cell, assigning force field parameters, filling the cell with water, placing counter ions, predicting pKa values and assigning protonation states, running an energy minimization, setting initial atom velocities according to a Boltzmann distribution. Simulation parameters were kept at the values defined by the macro: temperature 298 K; solvent density 0.997g/ml; AMBER14 force field. Snapshots of the trajectories were taken at every 100 picoseconds. The RMSD (Root Mean Square Deviation) of the backbone of the protein structure was calculated using the YASARA Structure. All RMSD values were obtained for all backbone atoms and were calculated with reference to the starting structure.

### Co-immunoprecipitation and western blotting analysis of DSS1 and BRCA2’s variants

Co-immunoprecipitation was performed using Dynabeads Co-Immunoprecipitation Kit (ThermoFisher Scientific, CAT# 14321D) following the protocol supplied by the manufacturer. 293T cells were co-transfected with myc-tagged DSS1 and Flag-tagged BRCA2’s variants. Forty-eight hours after transfection, cells were harvested, weighted and resuspended in 1:9 ratio of cell mass to extraction buffer with protease inhibitors. For each immunoprecipitation, 1.5 mg of anti-FLAG antibody-coupled Dynabeads magnetic beads was added to 750 μl of cell lysate, incubated on a rotator at 4°C for 25 minutes. After washing on magnet to remove unbound protein, the immunoprecipitated protein complexes were eluted from Dynabeads in 60 μl of elution buffer. Western Blotting was performed using a Simple Western Wes machine (Proteinsimple). The equal amount of each samples was loaded into the capillary automatically and separated by size as they migrate through a stacking and separation matrix. The separated proteins were then immobilized to the capillary. The target proteins were identified using a primary antibody and immunoprobed using an HRP-conjugated secondary antibody and chemiluminescent substrate. The resulting chemiluminescent signal was captured and processed in Compass software. Sample data was presented as a lane view in a virtual-blot like image as well as a graph view with quantitative signal intensity (area). To detect DSS1 protein, a 2-40 kDa Separation Matrix (Proteinsimple, CAT# SM-W012) and anti-DSS1 antibody (Proteintech, CAT# 13639-1-AP) were used. For detecting BRCA2’s variants, the cell lysate and the FLAG-immunoprecipitated protein complexes were separated on a 66-440 kDa Separation Matrix (Proteinsimple, CAT# SM-W008) and detected using anti-FLAG (Cell Signaling Technology, D6W5B, CAT# 14793S) and anti-BRCA2 (Cell Signaling Technology, D9S6V, CAT# 10741S) antibodies.

### CRISPR/Cas9 knockdown

Cell line 293T (human immortalized embryonic kidney cells, female origin) was purchased from ATCC (American Type Culture Collection) and maintained in Dulbecco’s Modified Eagle Medium (Gibco, CAT# 11-965-092) containing 10% heat inactivated fetal bovine serum (BenchMark, CAT# 100-106), 10ug/ml cellmaxin (GenDEPOT, CAT# C3319-020) at 37°C and 5% CO2. Knockdown experiment was performed by using Alt-R® CRISPR/Cas9 System (IDT, IL, USA) following the manufacturer’s protocol. The Alt-R® CRISPR/Cas9 crRNAs were designed by using Alt-R® Custom CRISPR/Cas9 crRNA Design Tool (https://www.idtdna.com/site/order/designtool/index/CRISPR_CUSTOM) and purchased from IDT. To knockdown BRCA2, a crRNA that targets the second exon of BRCA2 was used. The crRNA was incubated with tracrRNA (IDT, CAT# 1072534), and then Alt-R® S.p.Cas9 Nuclease (IDT, CAT# 1081059) to prepare Cas9:crRNA:tracrRNA ribonucleoprotein (RNP) complex, the RNP complex was then delivered into 293T cells using Lipofectamine CRISPRMAX Cas9 Transfection Reagent (Invitrogen, CAT# CMAX00008) according to the manufacturer’s protocol. 48 hours after transfection, the cells were harvested and subjected to T7 endonuclease I (T7EI) mismatch detection assays using the Alt-R® Genome Editing Detection Kit (IDT, CAT# 1075932). The limiting dilution approach was used to isolate individual cell clones following CRISPR/Cas9 gene editing. Isolated single cell clones were expanded and frozen for further validation and banking. Sequence alterations of BRCA2 gene at exon 2 were confirmed by Sanger sequencing using CRISP-ID analysis^98^ (version 1.1, http://crispid.gbiomed.kuleuven.be/) and come up with the consensus manually.

Expression of BRCA2 at protein level was confirmed by western blotting using NuPAGE® Large Protein Blotting Kit (Invitrogen, CAT# LP0001) following the protocol supplied by the manufacturer. Cell lysates were prepared by using RIPA lysis buffer (Thermo Fisher Scientific, CATCAT# 89900) supplemented with protease inhibitor cocktail (Sigma-Aldrich, CAT# P8340). Equal amounts of protein (50 ug) were separated on SDS-polyacrylamide gel (Thermo Fisher Scientific, CATCAT# EA0378BOX) and transferred onto nitrocellulose membranes (Invitrogen, CATCAT# LC2001). Each membrane was hybridized with primary antibody followed by horseradish peroxidase (HRP)-linked IgG and visualized by the enhanced chemiluminescence (ECL) system (GeneTex, CATCAT# GTX14698). The signaling was captured by using FUJIFILM X-A2 digital camera (FUJIFILM, USA), the resulting images were analyzed using ImageJ (version 1.52a, https://imagej.nih.gov/ij/). The antibodies used for this assay were: BRCA2 (Cell Signaling Technology, CATCAT# 10741), β-TUBULIN (Cell Signaling Technology, CATCAT# 2128), and anti-rabbit IgG, HRP-linked antibody (Cell Signaling Technology, CATCAT# 7074).

### Plasmids constructs

Human BRCA2 (GenBank NM_000059.3; 228-10481) cDNA was cloned into the BamHI-NotI of the pcDNA3.1(+)-myc-His expression vector (Invitrogen, CAT# V80020) yielding pcDNA3.1-BRCA2T (BRCA2/2662T). BRCA2 mutation expression vector pcDNA3.1-BRCA2M (BRCA2/2662M) was created by using a site-directed mutagenesis strategy. To construct vectors expressing BRCA2/2662T or BRCA2/2662M fused with the GAL4 DNA-binding domain (GBD), BRCA2/2662T and BRCA2/2662M cDNA fragment was subcloned into the XbaI-NotI sites of pBIND vector (Promega, CAT# E2440), generating pBIND-BRCA2-2662T and pBIND-BRCA2-2662M. To construct vector expressing RAD51 fused with the VP16-activation domain (VAD), RAD51 cDNA fragment (GenBank NM_002875.4; 300-1316) was inserted into the XbaI-NotI sites of pACT vector (Promega, CAT# E2440), producing pACT-RAD51.

### Mammalian two-hybrid assay

Mammalian two-hybrid assay was performed using CheckMate™ Mammalian Two-Hybrid System (Promega, CAT# E2440) following the protocol supplied by the manufacturer. The following vectors were used in this assay. The VAD-expression vector was pACT-RAD51 and GBD-expression vectors used were pBIND-BRCA2-2662T and pBIND-BRCA2-2662M. pG5luc (Promega, CAT# E2440) was used as a reporter in which the firefly luciferase (luc+) gene was driven by a major late promoter of adenovirus connected to five GAL4 binding sites. VAD (pACT) and GBD (pBIND) cloning vectors were used as basal control for the background level of luciferase. The activity of the *Renilla* luciferase that is expressed in GBD expression vector pBIND, was used for normalizing the transfection efficiency. pBIND-Id and pACT-MyoD (Promega, CAT# E2440) that express two proteins (GAL4:Id and VP16:MyoD fusion proteins) known to interact in vivo, were used as a positive control. The VAD expression vector, GBD expression vector and pG5luc vector were co-transfected into 293T cells using Lipofectamine 2000 Transfection Reagent (Invitrogen, CAT# 11668019) according to the manufacturer’s protocol. The total amount of transfected DNA was adjusted by adding pcDNA3.1(−) empty (Invitrogen, CAT# V79520) vector. Expression of GAL4:BRCA2 and VP16:RAD51 at protein level was confirmed by western blotting as described above. The antibody used for VP16:RAD51 was anti-Rad51 antibody (abcam, CAT# ab63801). Thirty-six hours after transfection, cells were harvested and tested for luciferase activity using a Dual-Luciferase® Reporter Assay System (Promega, CAT# E1980). Luciferase activity was determined with the Cytation 5 Cell Imaging Multi-Mode Reader (BioTek, VT, USA) and Gen5 (version 2.09, https://www.biotek.com/products/software-robotics-software/gen5-microplate-reader-and-imager-software/). Firefly luciferase activities were normalized by *Renilla* luciferase activities. Quadruplicate experiments were done for each transfection. Prism was used for visualization. To test for significant differences, two-way ANOVA analysis in R statistical environment was used.

### DSB Repair Assays

The BRCA2-deficient V-C8 hamster cell line contains a single integrated copy of DR-GFP reporter. For DSB repair assays, 5×10^6^ exponentially growing cells were collected and resuspended in 650 μl Opti-MEM media (Invitrogen) in a 0.4-cm cuvette (Biorad) with 50 μg each of the I-SceI expression vector pCBASce (or the empty pCAGGS vector) and the expression vector for mini-BRCA2. Cells were transfected by electroporation (280 V and 1000 μF) with a Biorad Gene Pulser II. DSB repair was measured by counting the number of GFP-positive cells using Aurora spectral analyzer (Cytek Biosciences) or FACSAria sorter (BD) 48 h after electroporation.

### Statistical analysis

For statistical analysis of dN/dS, boxplots were made by using Prism (version 7, https://www.graphpad.com/scientific-software/prism/). Maximum, upper quartile, median, lower quartile, minimum and mean values were shown in Supplementary Fig. 4d. To test significance levels of adaptive selection index in McDonald-Kreitman test, one-sided Fisher exact test was used (https://www.socscistatistics.com/tests/Default.aspx). To test for significant differences of results of co-immunoprecipitation assay of BRCA2 and DSS1, one sample t-test in R statistical environment was used. To test for significant differences of results of mammalian two-hybrid assay and DSB repair assay, two-way ANOVA analysis in R statistical environment was used.

## Acknowledgments

Supported by NIH grants R01 CA179991, R35 CA220508 and funding from the MSK Functional Genomics Initiative (C.A.I.-D.), NCI Cancer Center Support Grant P30 CA08748 to MSKCC, Cycle for Survival and Kleberg foundation.

## Author contributions

J.H., Y.Z., A.P.M.-M., W.C.W., M.A.N. and C.A.I.-D. designed various portions of the study; J.H., W.C.W., and A.P.M.-M. performed analyses; M.J. and T.W. provided key reagents for functional studies, J.H., and Y.Z. performed experiments; J.H., Y.Z., W.C.W., M.A.N., and C.A.I.-D. wrote the manuscript; W.C.W., M.A.N., M.J. and A.P.M.-M. revised the manuscript. C.A.I.-D. supervised the program.

## Data and materials availability

All data that support the findings of this study are included in this published article and supplementary materials. All data generated or analyzed during this study are available from the corresponding author upon request.

## Supplementary Materials

Supplementary Figs. 1-20

Supplementary Tables 1-12

