## SupplementaryInformation for "Selection for Decreased BRCA2 Functional Activity in *Homo sapiens* After Divergence from the Chimpanzee-Human Last Common Ancestor"

**This file includes:**

Supplementary Figs. 1-20

Supplementary Tables 1-3, 8, 12

Captions for Supplementary Tables 4-7, 9-11

**Other Supplementary Materials for this manuscript include the following:**

Supplementary Tables 4-7, 9-11 (.xlsx)

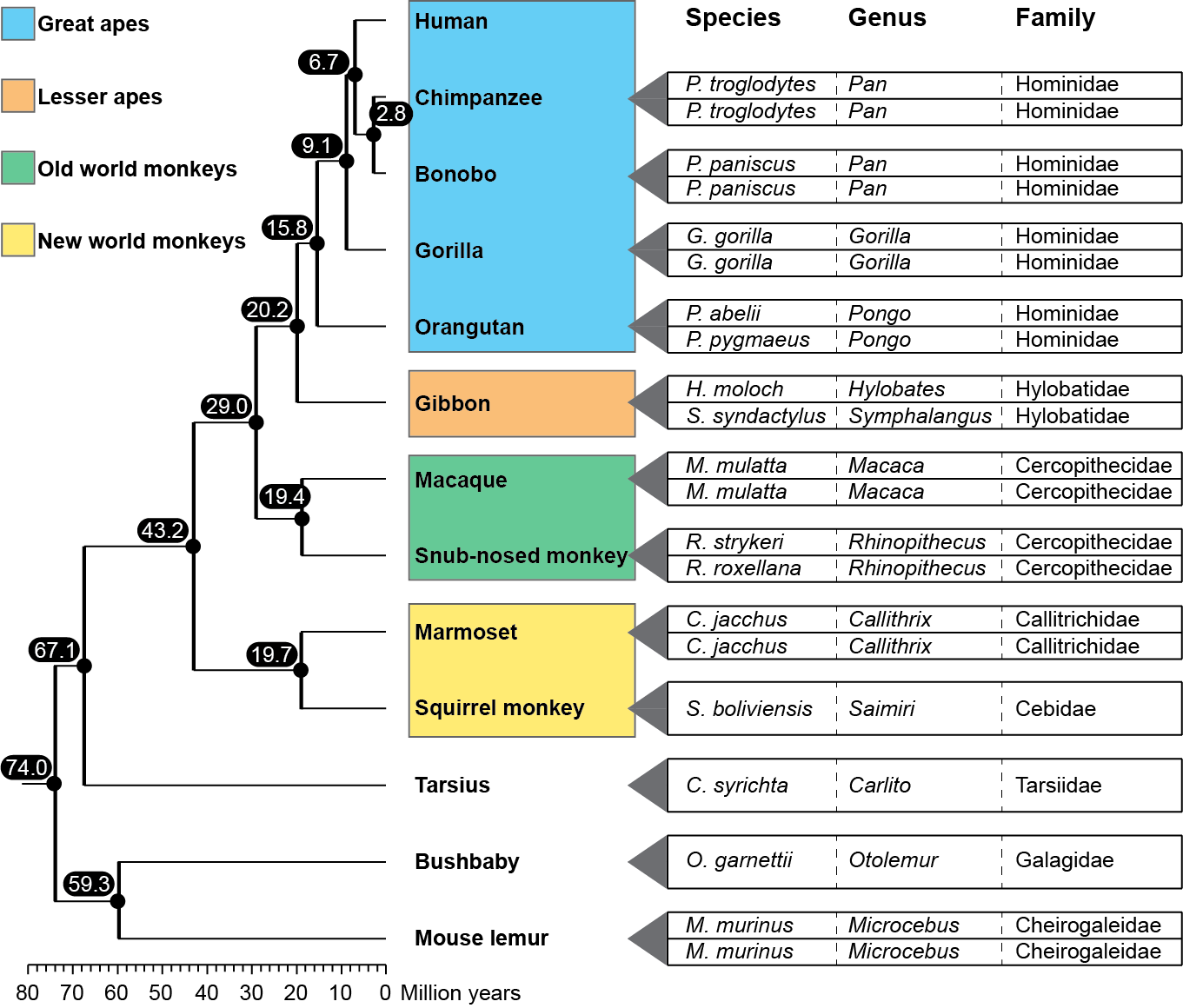

**Supplementary Fig. 1 | Taxonomic and phylogenetic relationships of 21 primate genomes.** The phylogenetic relationship of primates referred to Enard et al.^22^. Twelve groups of species are shown on the left edges. Generally, two genomes were used for each group. However, there is only one available genome for each of squirrel monkey, tarsius and bushbaby. Details for each genome are list on the right edges. The separation time of species was obtained from Timetree (<http://www.timetree.org/>).

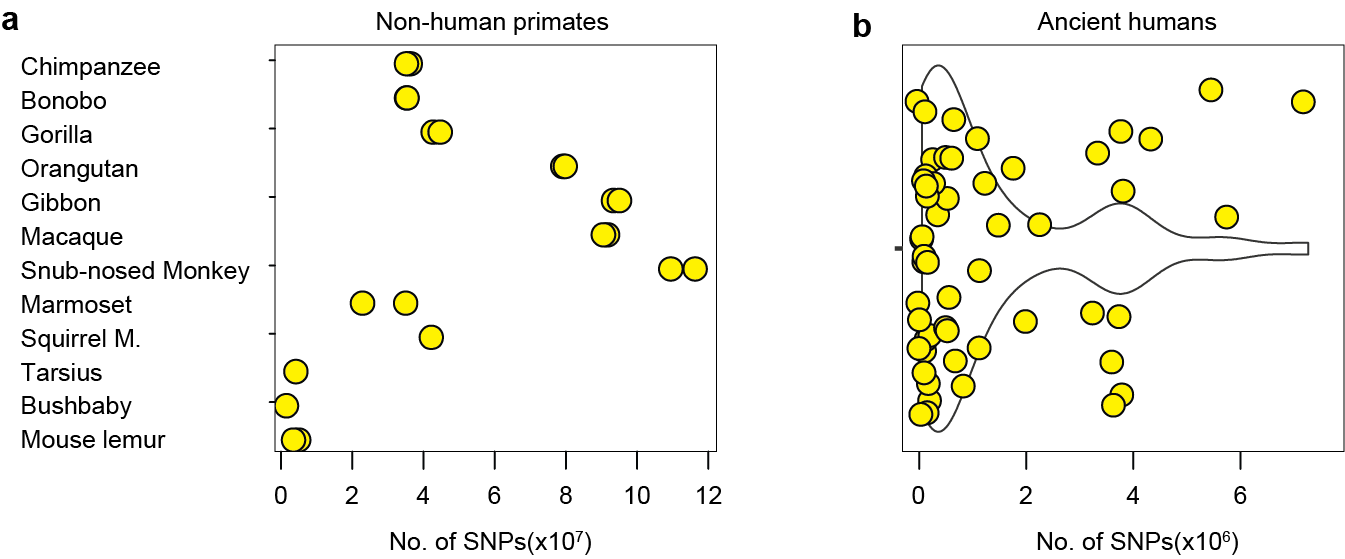

**Supplementary Fig. 2 | Construction of a variations database of primate whole genomes. (a)** Number of SNPs in 21 genomes of 12 non-human primate groups. **(b)** Number of SNPs in 54 ancient human genomes.

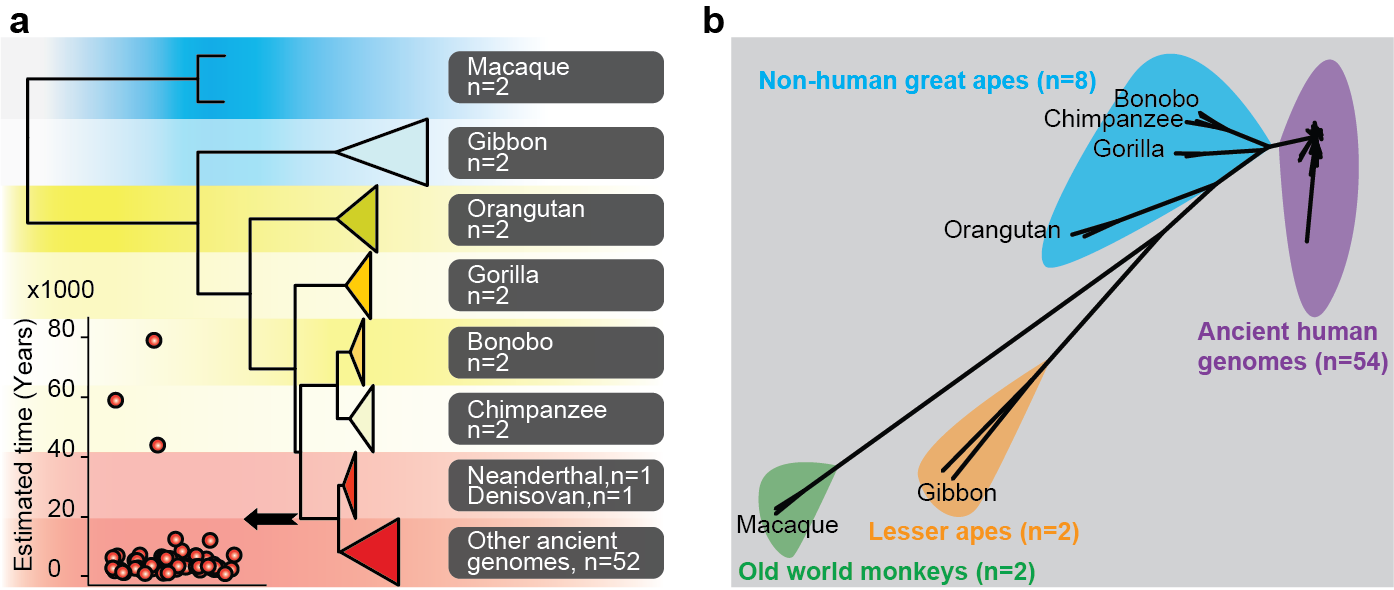

**Supplementary Fig. 3 | Phylogenetic reconstruction. (a)** Phylogenetic tree using non-silent SNPs of old world monkeys (Macaque, n=2), lesser apes (Gibbon, n=2), non-human great apes (n=8) and ancient human genomes (n=54). In the lower left, estimated time of ancient human genomes was shown. **(b)** Phylogenetic reconstruction using APE package based on 1,561,780 non-silent SNPs.

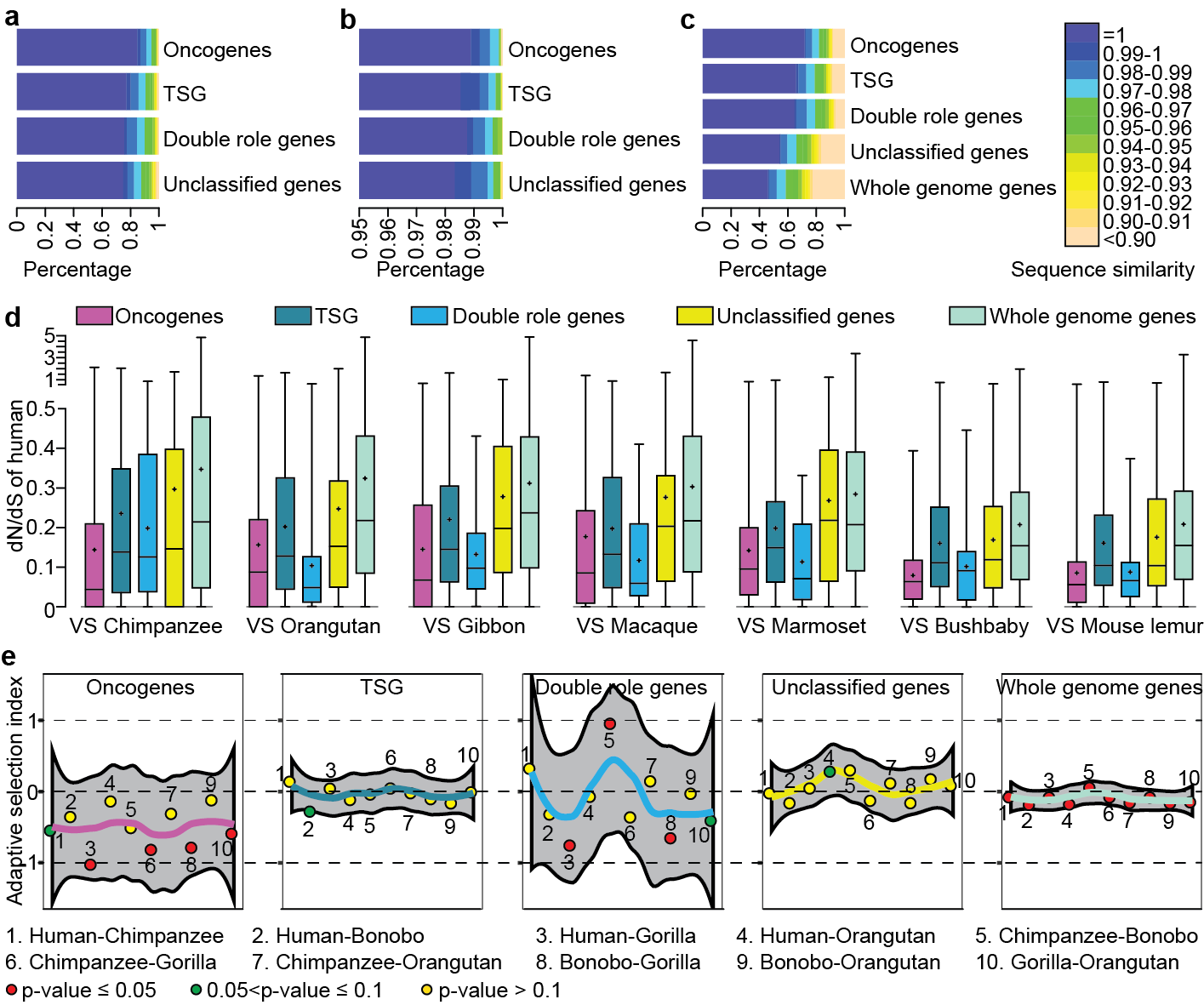

**Supplementary Fig. 4 | Evolutionary features of cancer genes. (a)** Exon sequence similarity for cancer genes using non-silent SNPs of 21 non-human primate genomes. TSG: Tumor Suppressor Genes. **(b)** Exon sequence similarity for cancer genes using non-silent SNPs of 54 ancient human genomes. **(c)** Exon sequence similarity for cancer genes and whole genome genes using protein sequences derived from 12 assembled non-human primate genomes. Panels (a) to (c) share the same color bar. **(d)** Comparison of dN/dS for cancer genes and whole genome genes between human reference genome and other seven species of primates. Boxplots depict maximum, upper quartile, median, lower quartile and minimum. "+" denote mean values. **(e)** McDonald–Kreitman test for cancer genes and whole genome genes using genomic SNPs of 9 present-day humans and 79 non-human great apes. Adaptive selection index was shown to measure selection pressure.

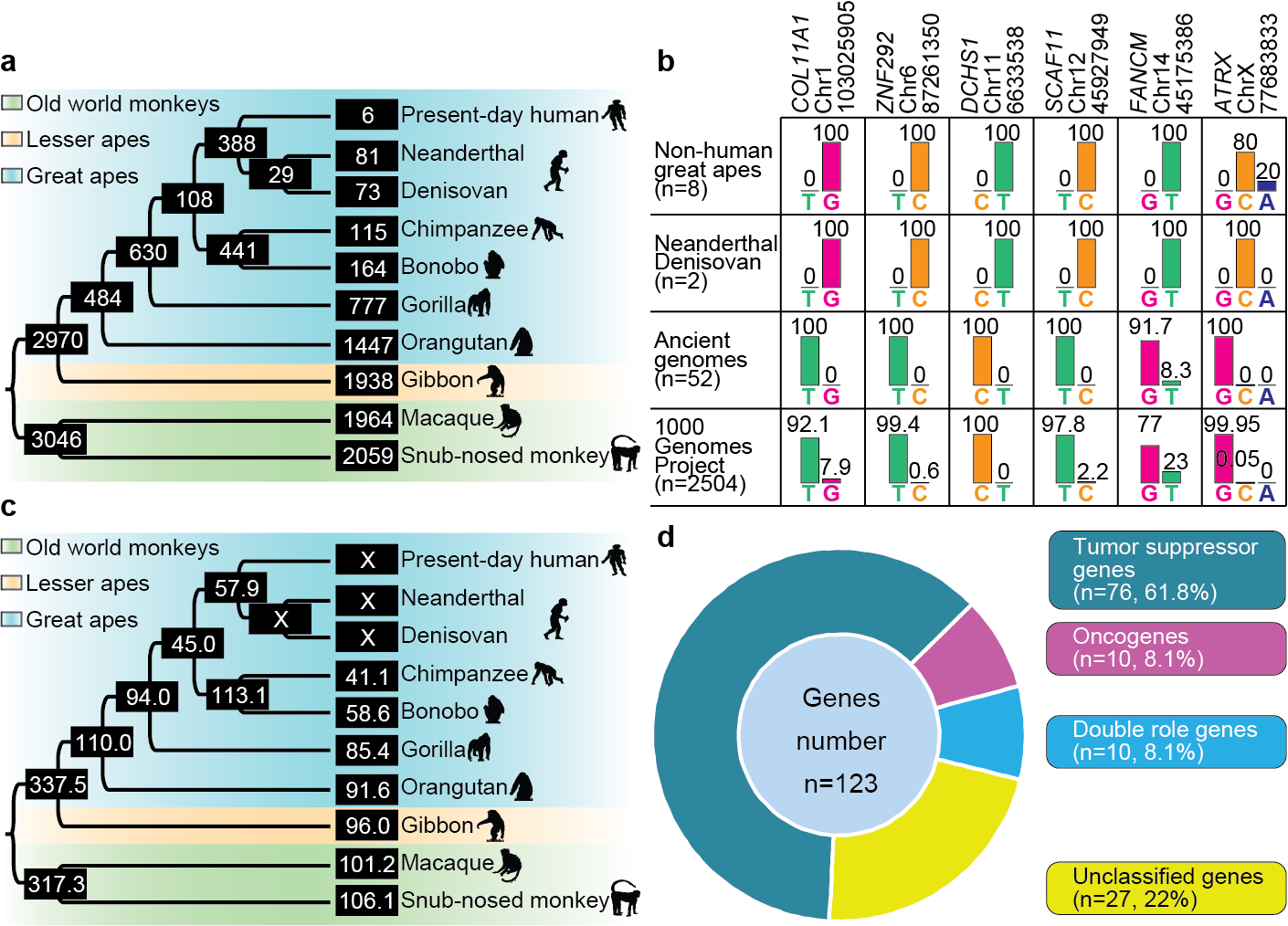

**Supplementary Fig. 5 |** **Emergent genetic variations during evolution of primates. (a)** Emergent SNPs for each branch. The figures in the box denote the number of SNPs. **(b)** Present-day human accumulated six SNPs when compared with Neanderthal/Denisovan. Genotype frequency was estimated in eight non-human great apes (two each of *Pan troglodytes*, *Pan paniscus*, *Gorilla gorilla*, and one each *Pongo abelii* and *Pongo pygmaeus*), Neanderthal/Denisovan (n=2), other ancient human genomes (n=52) and 2504 present-day humans in the 1000 Genomes Project. **(c)** Estimated mutation rate for cancer genes using emergent genetic variations of each branch. The figures in the box denote the estimated mutation rate. "×" indicates the time of divergence between the two branches is not available from TimeTree. **(d)** Distribution of 123 cancer genes with human-specific fixed SNPs in the four categories of cancer genes.

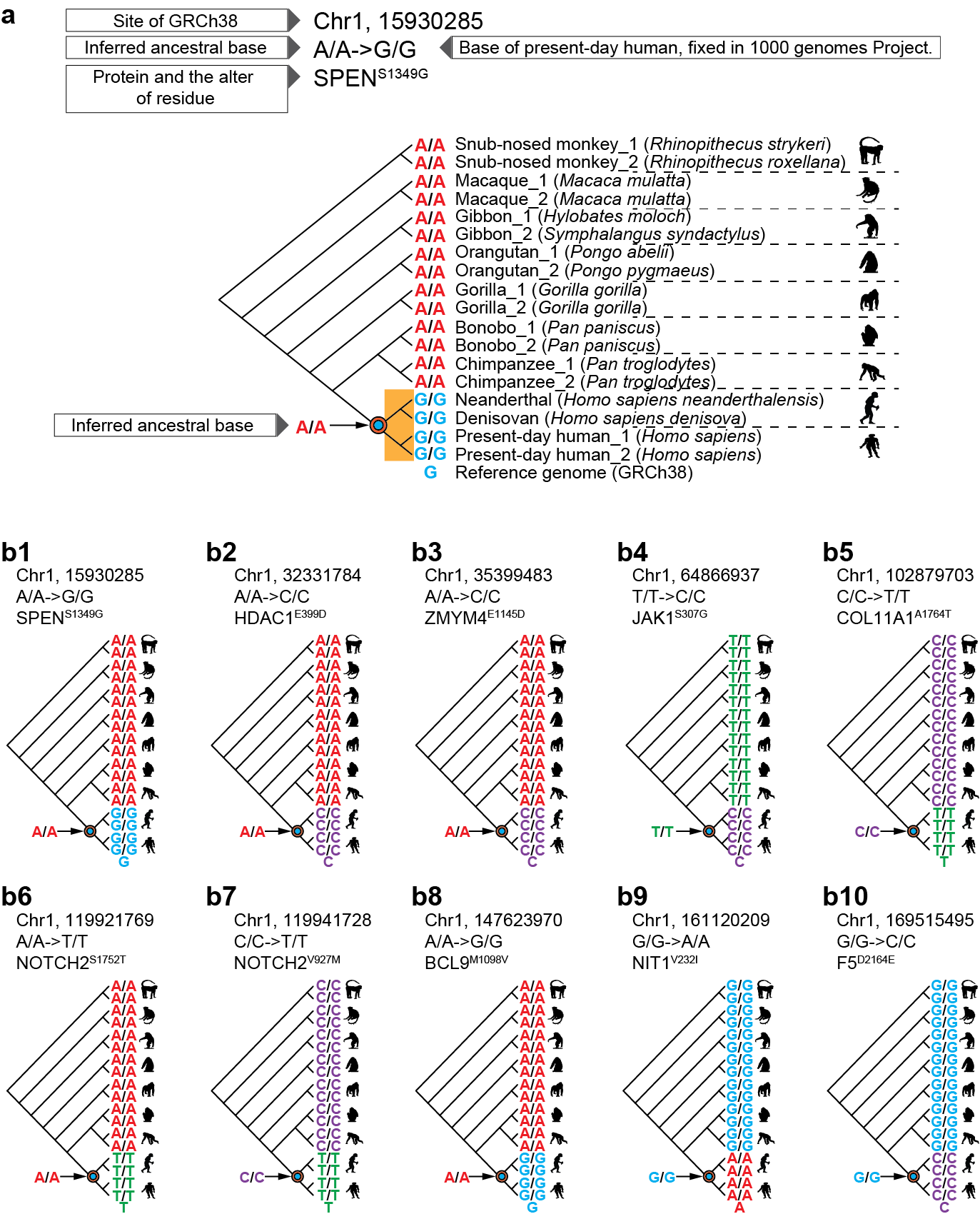

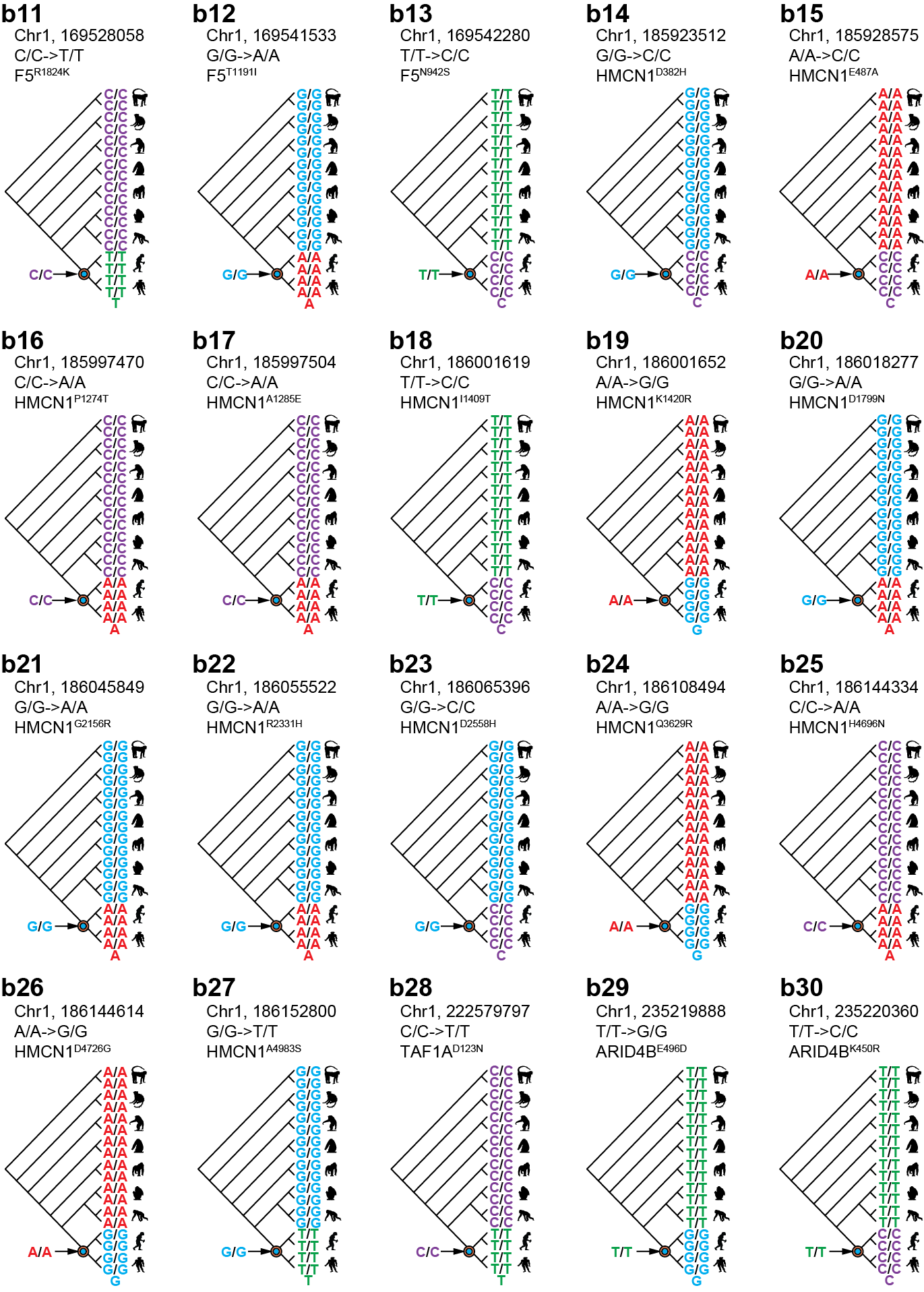

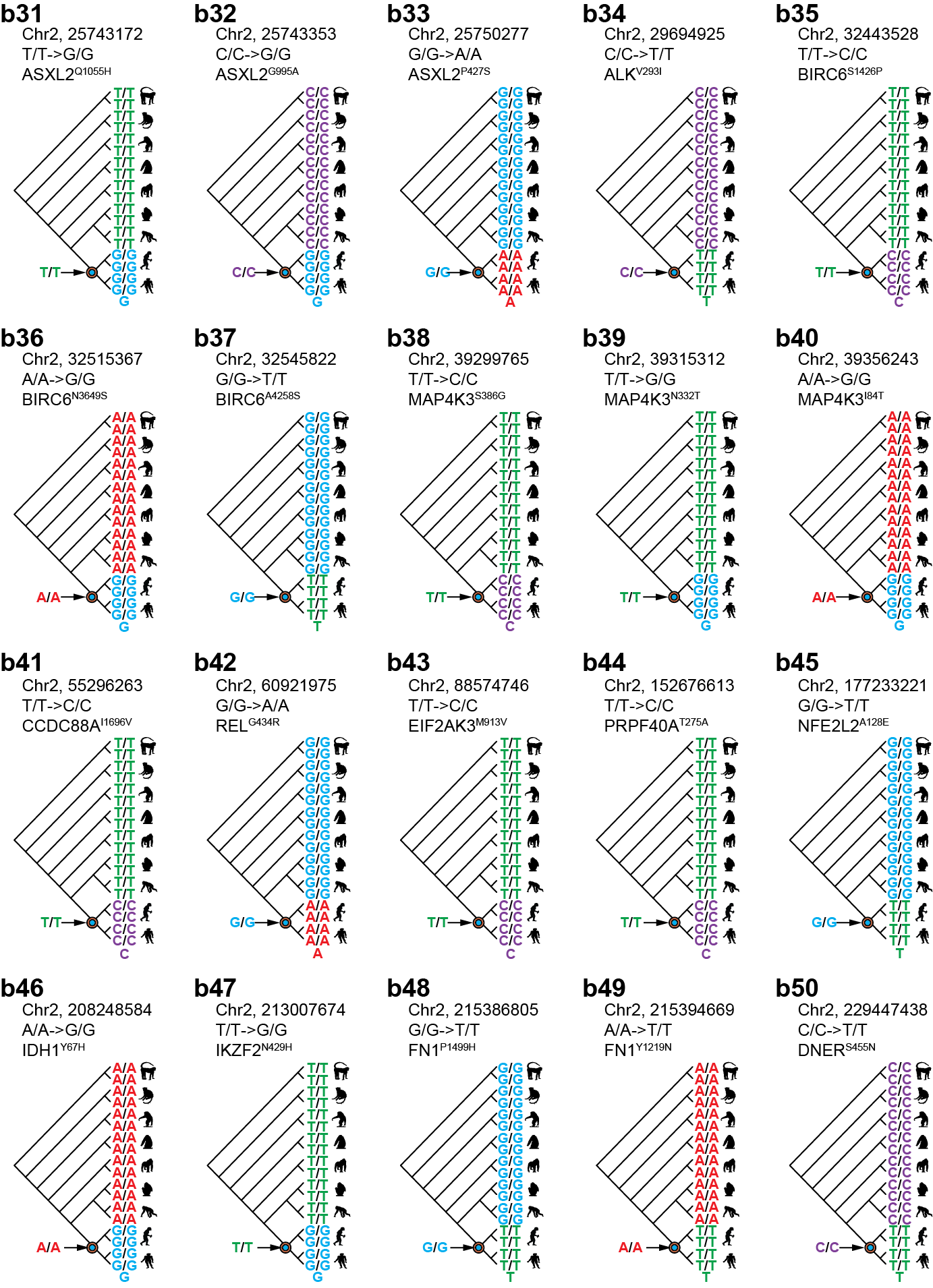

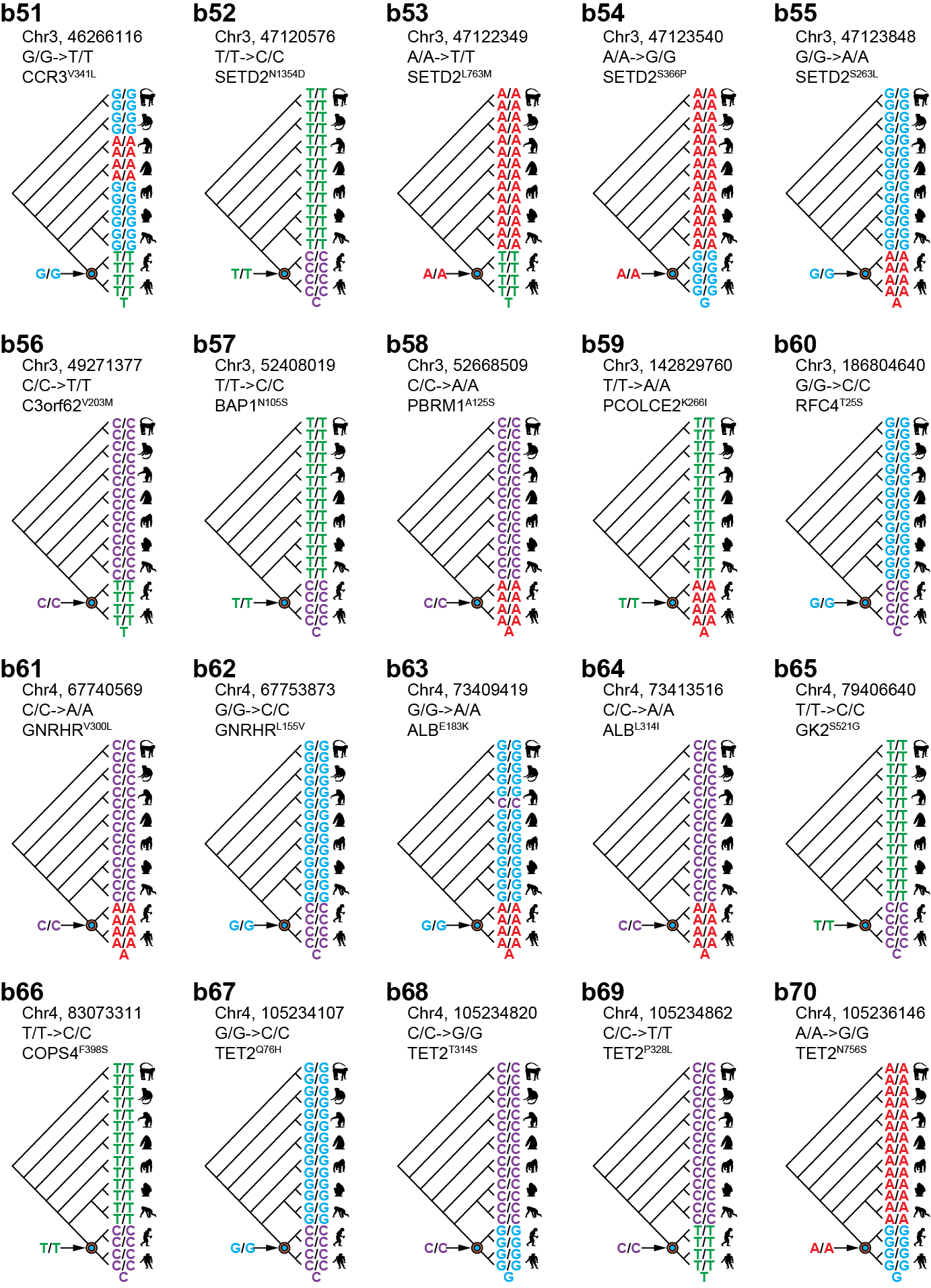

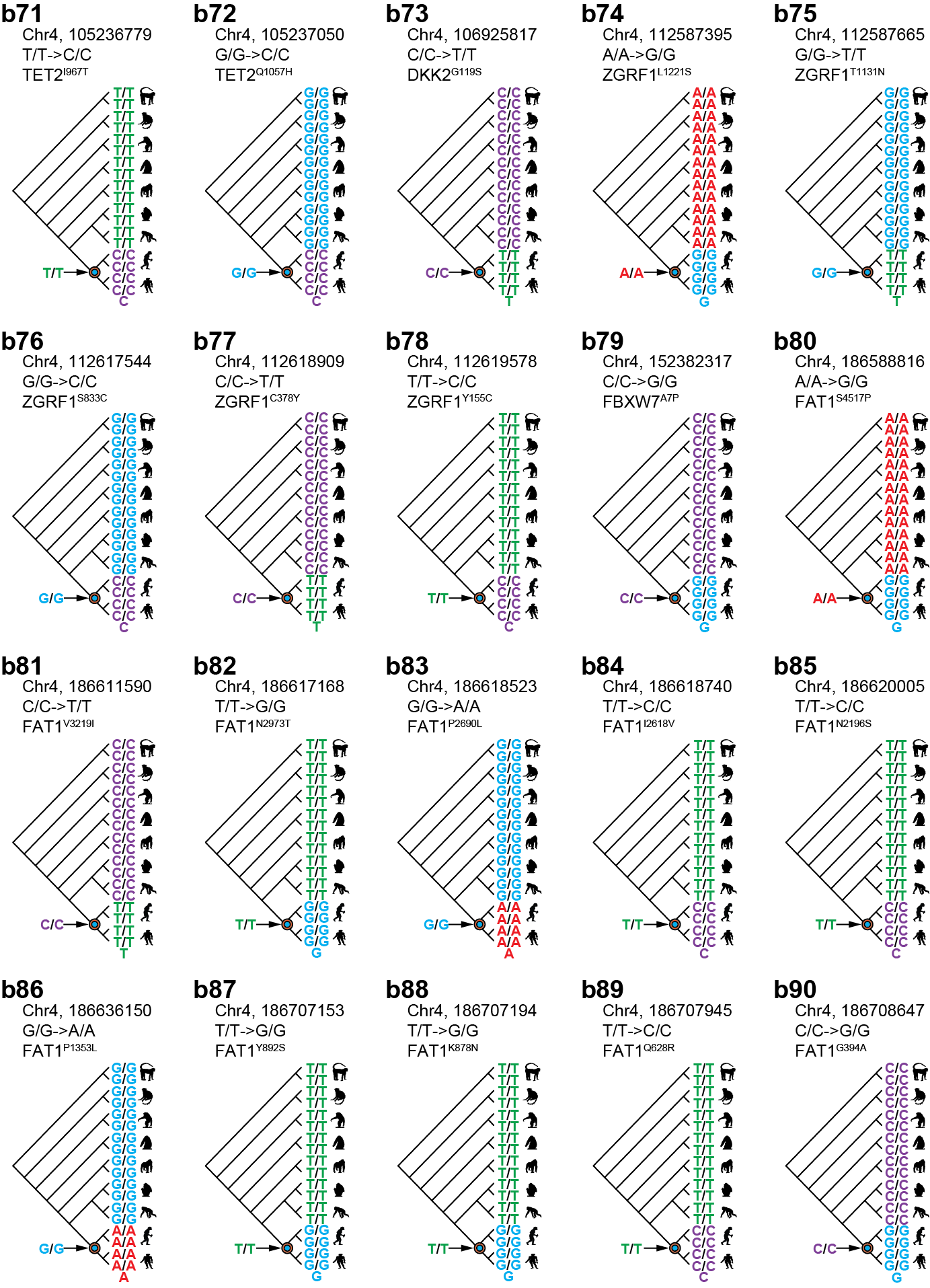

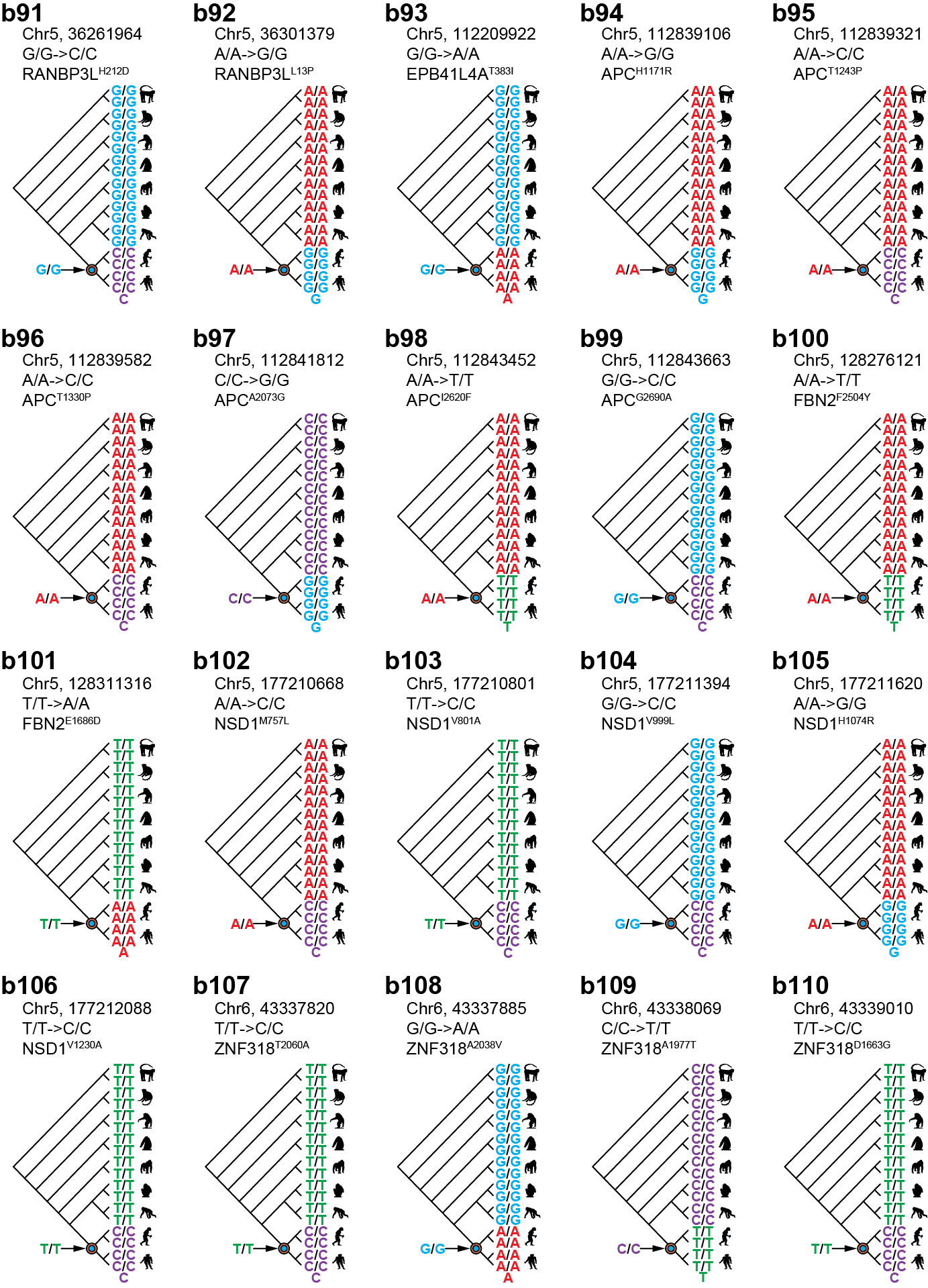

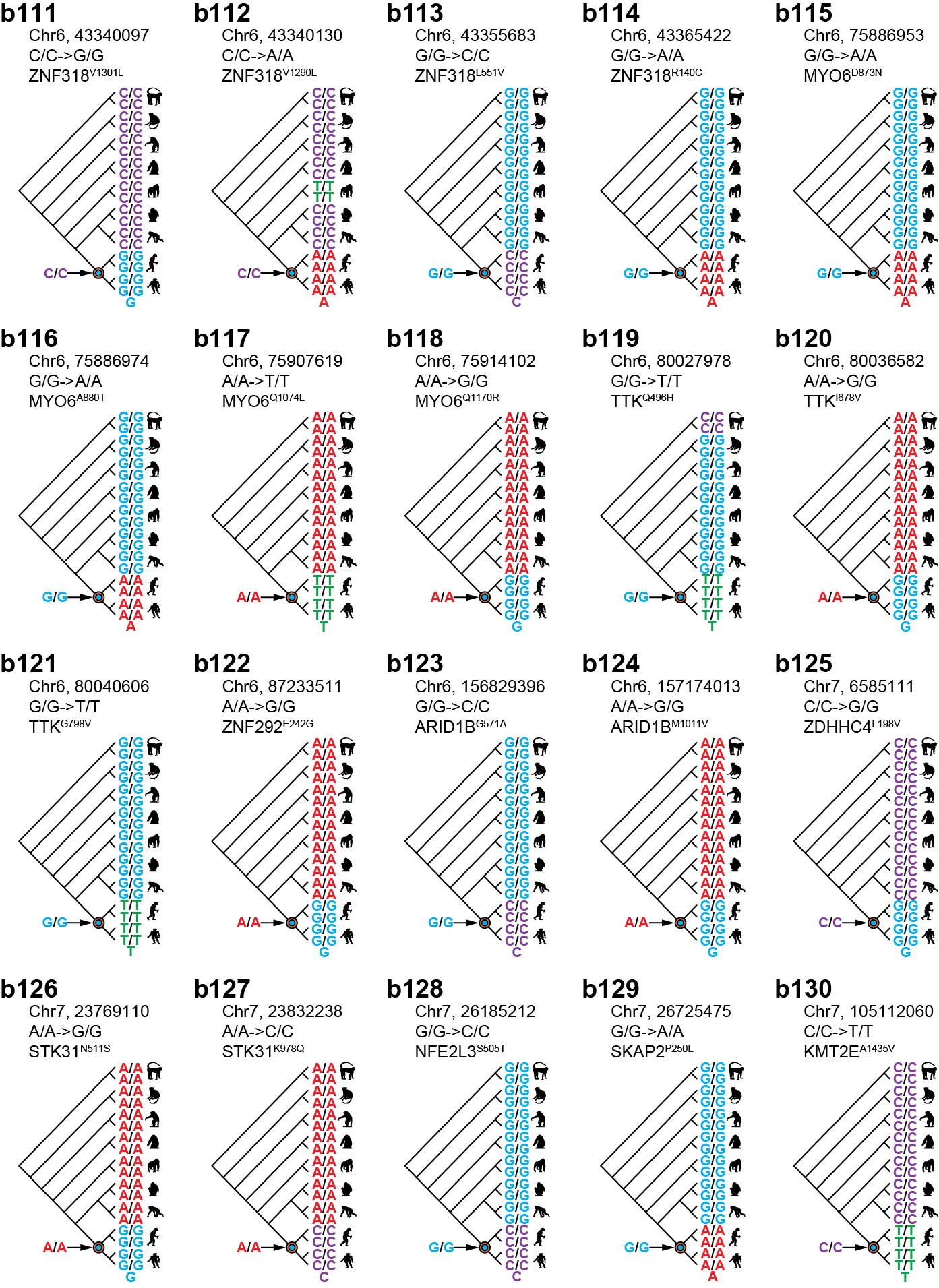

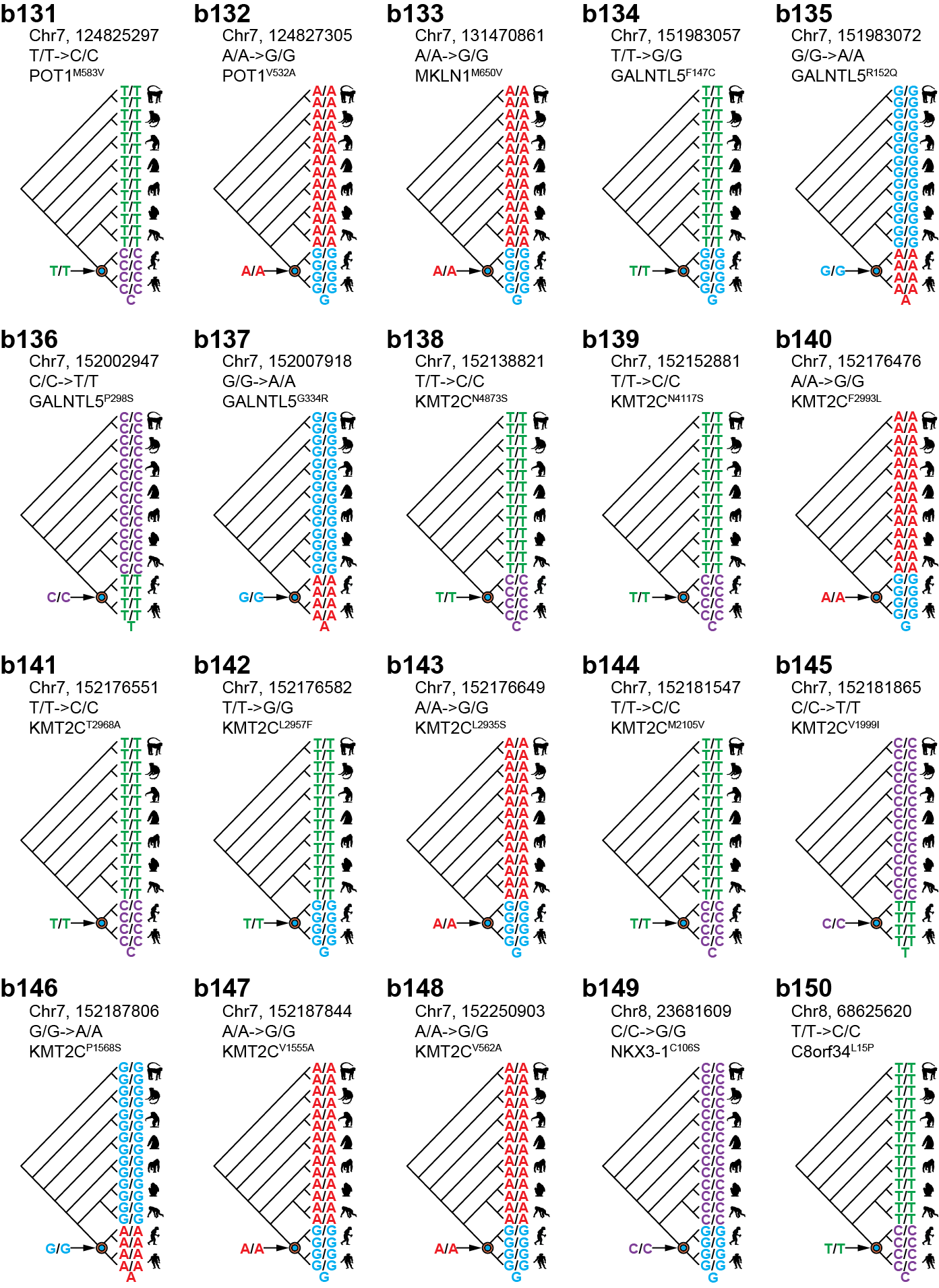

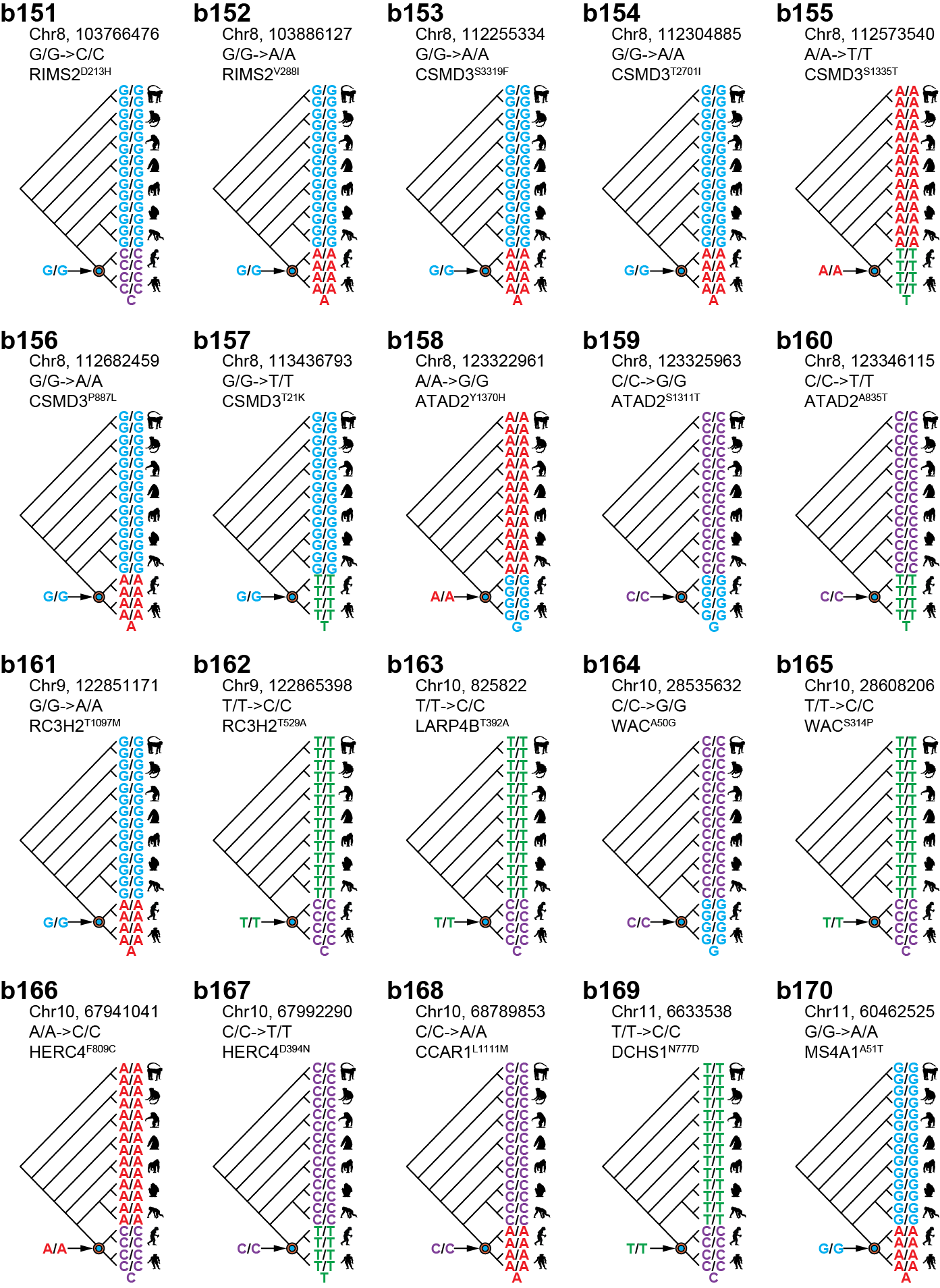

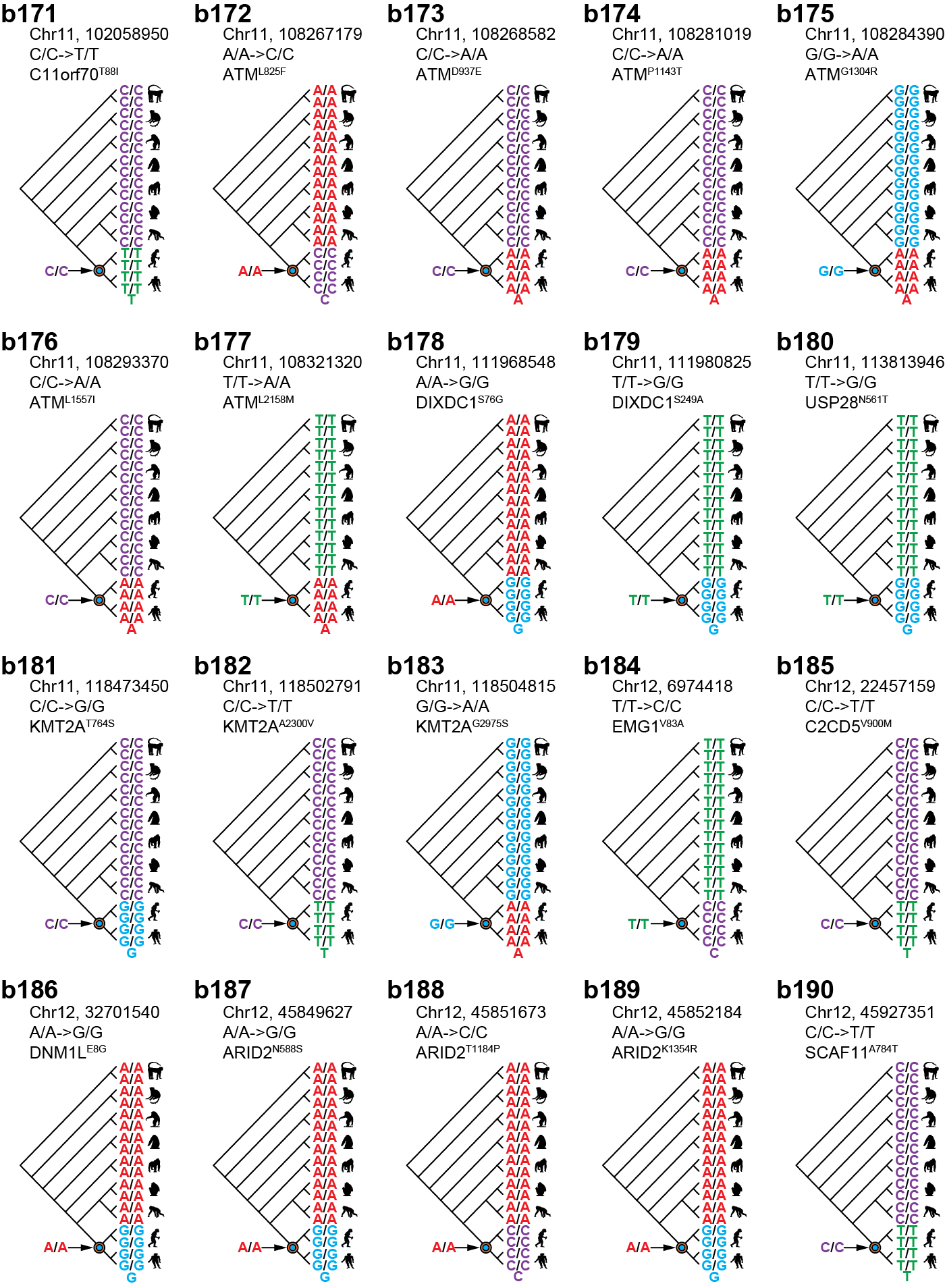

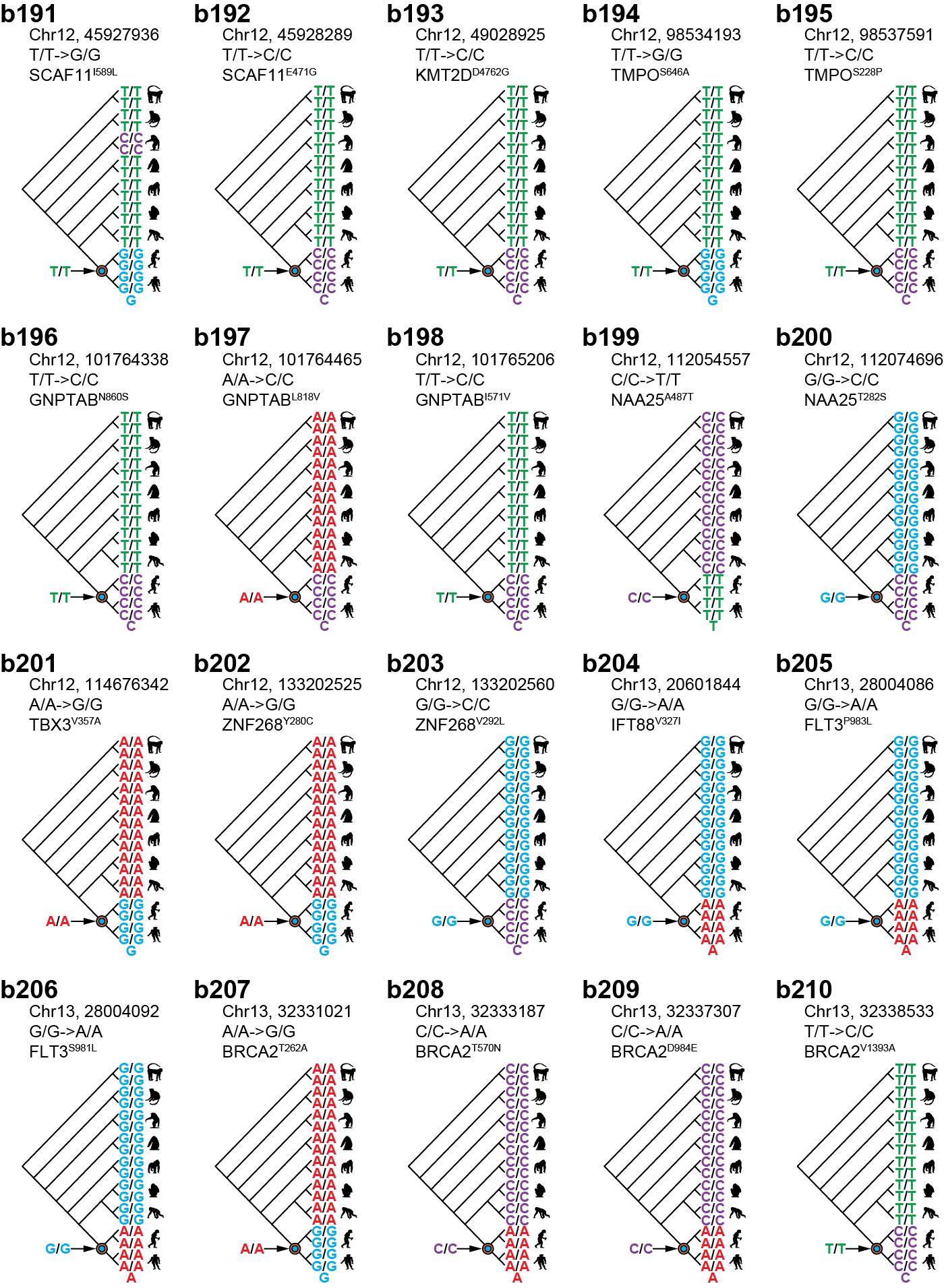

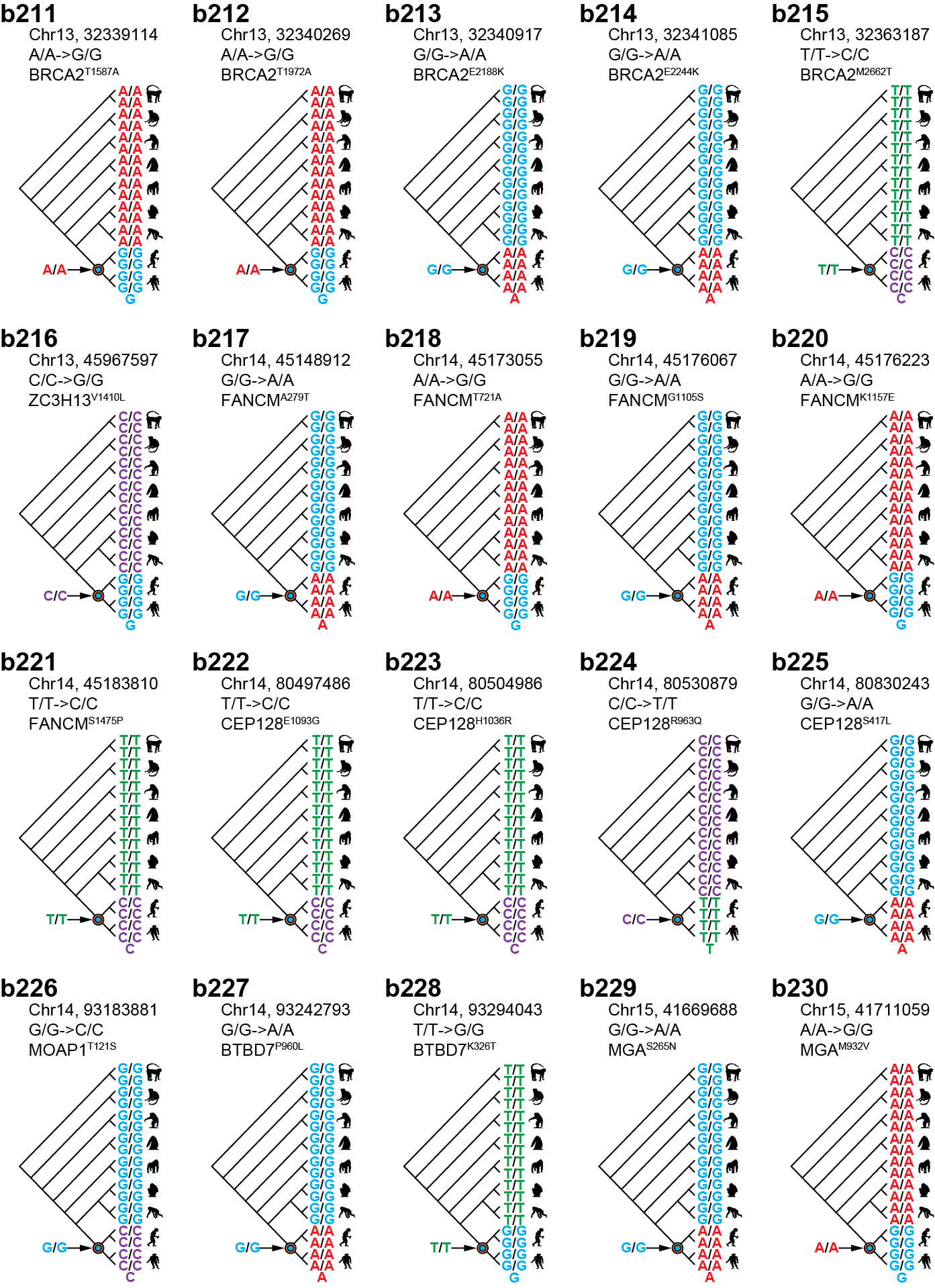

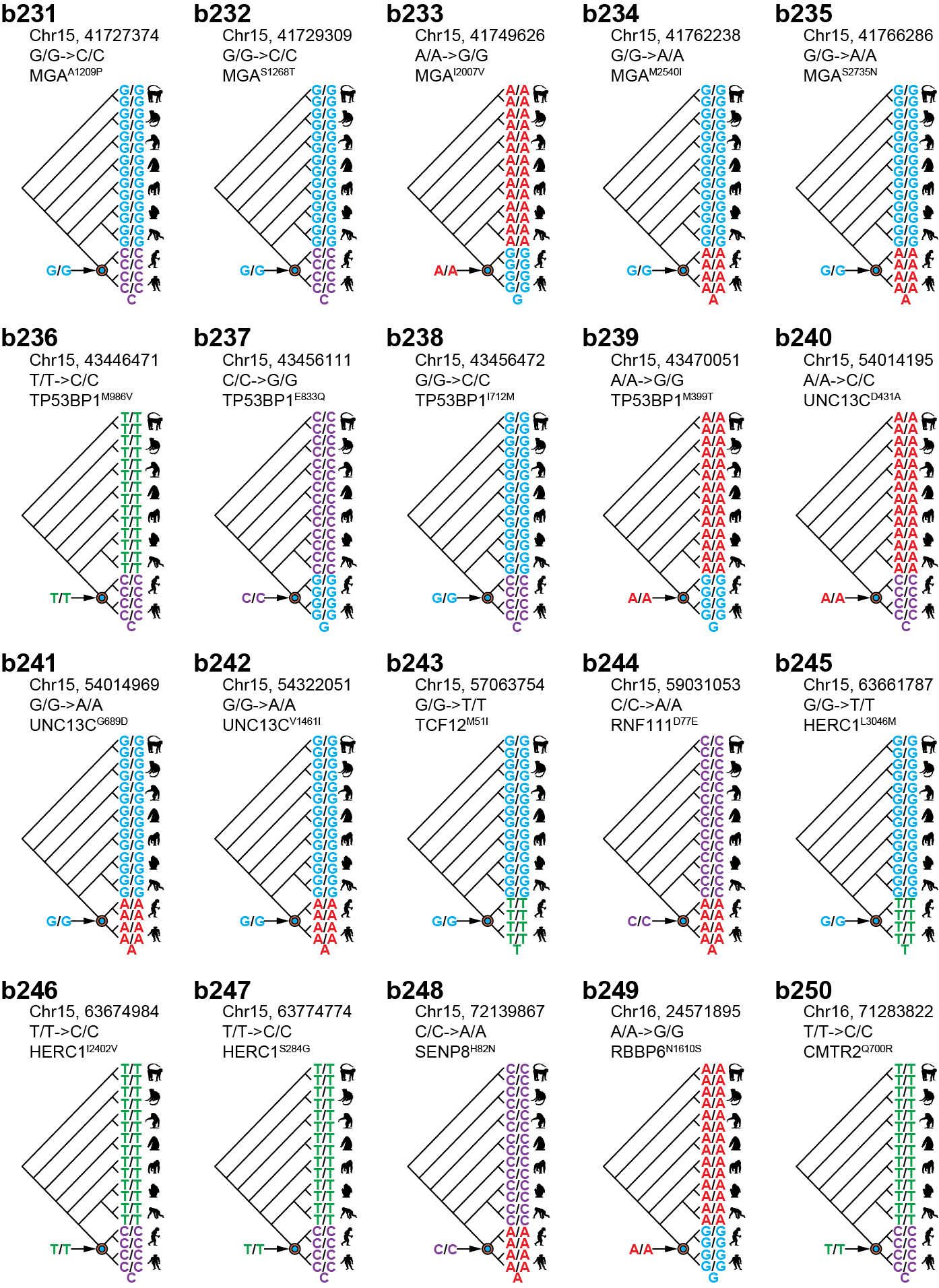

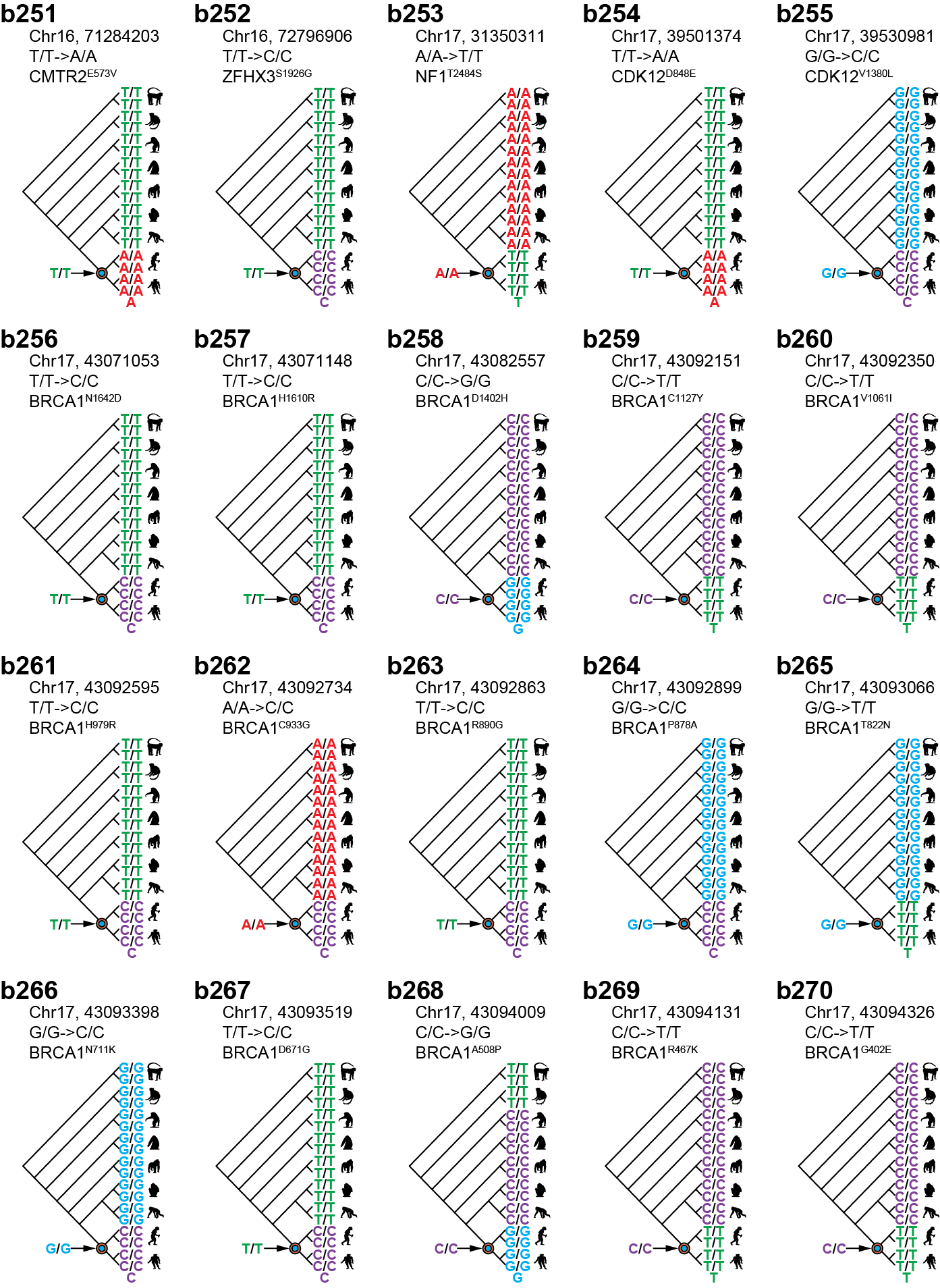

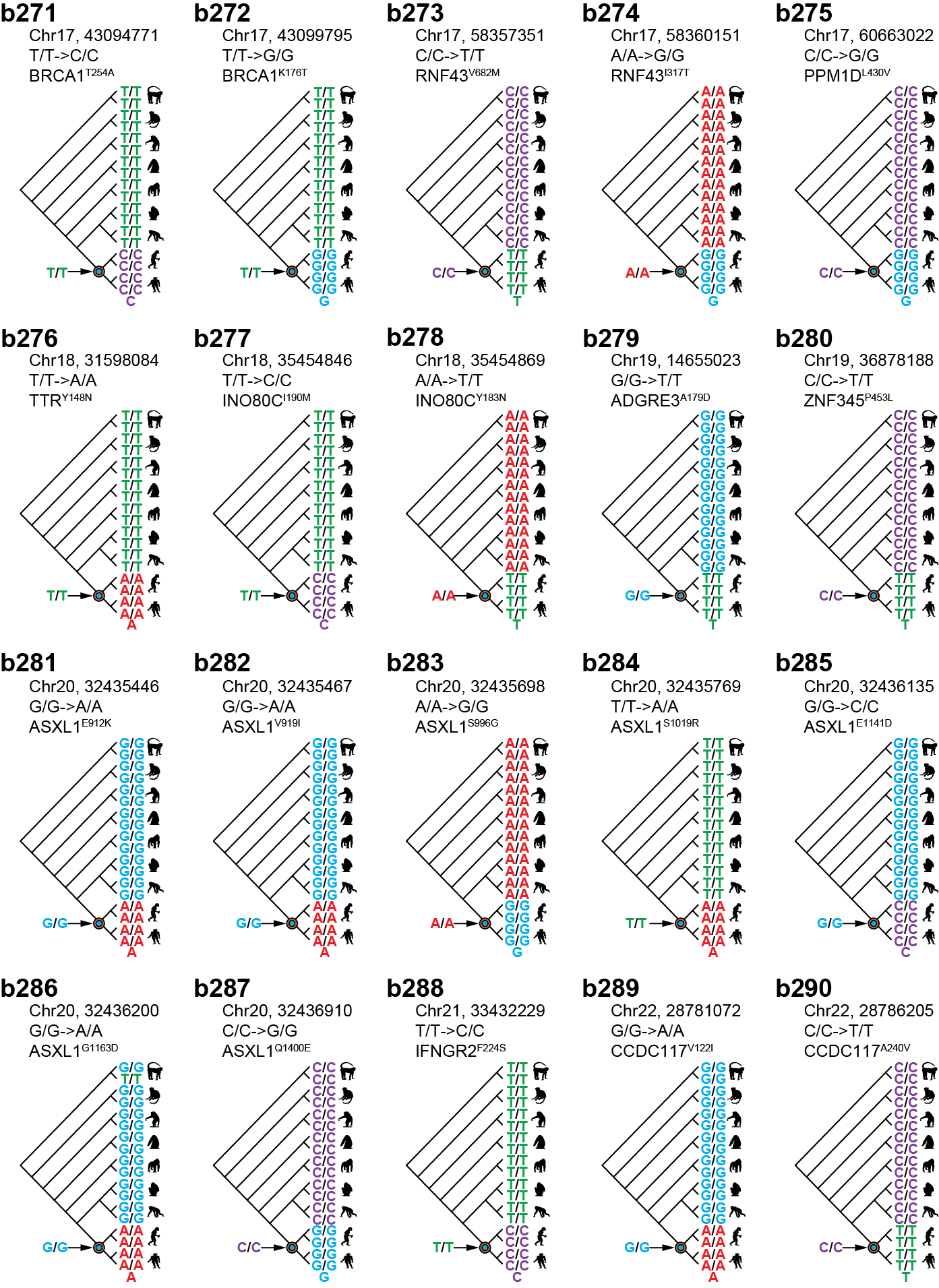

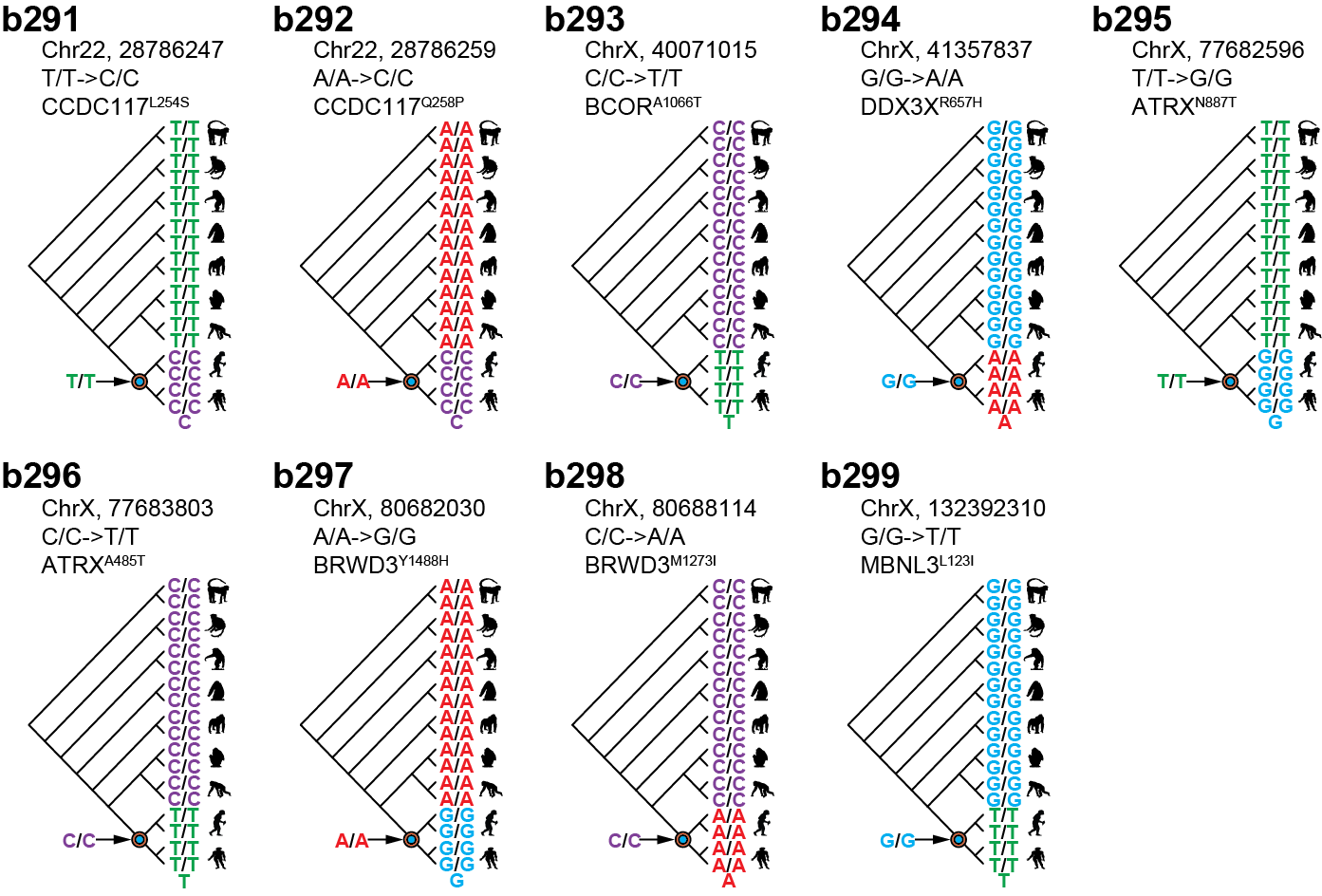

**Supplementary Fig. 6 | Infer ancestral genotype of human after divergence from the CHLCA. (a)** Eighteen genomes and human reference genome (GRCh38) that were used to infer ancestral base of human. **(b)** 299 human-specific fixed SNPs and the ancestral base.

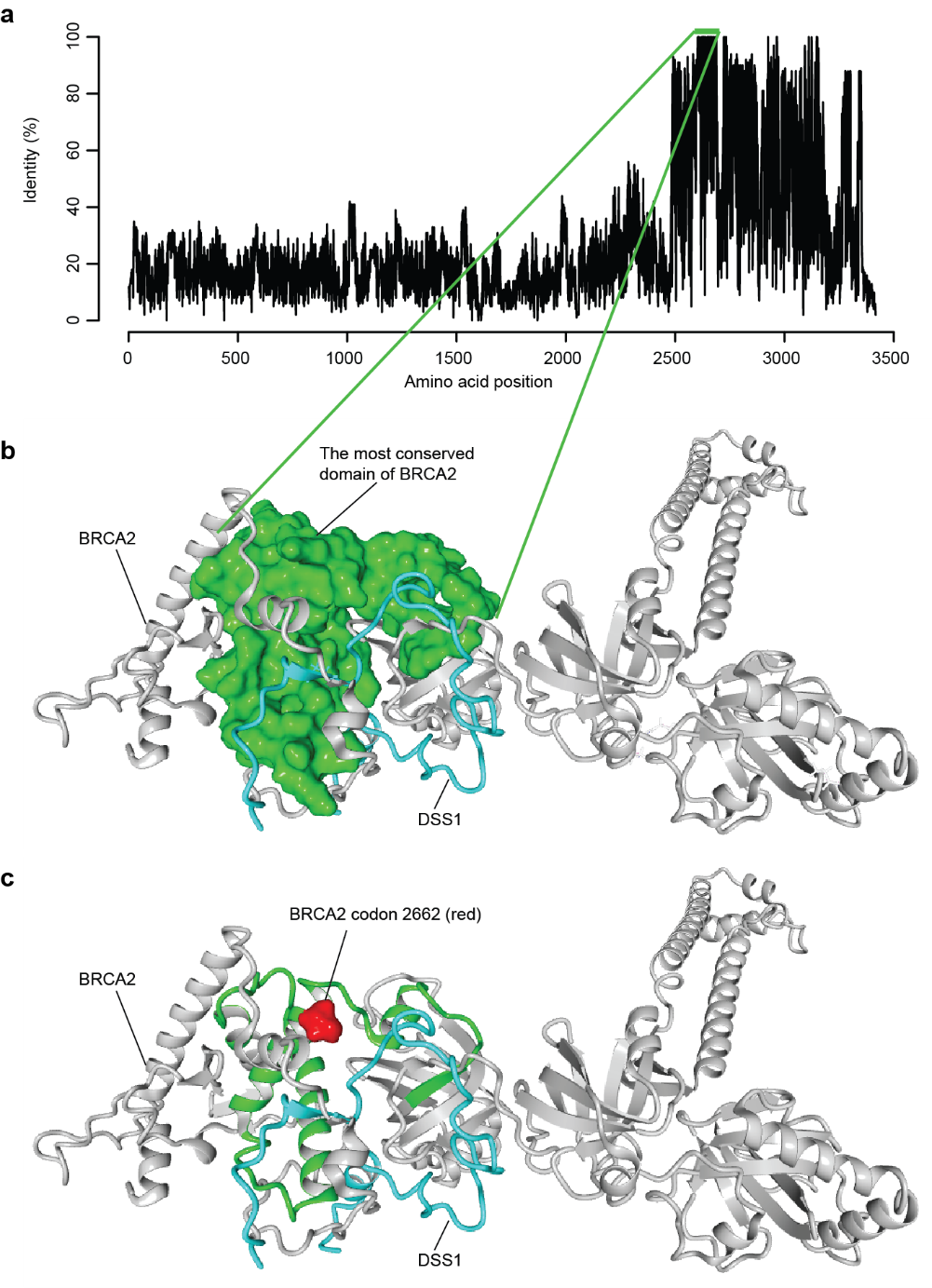

**Supplementary Fig. 7 | The most conserved domain of BRCA2 and the location of BRCA2 codon 2662.** **(a)** Sequence identity of whole protein sequence of BRCA2 of 56 osteichthyan species. The green line denotes the most conserved domain of BRCA2 (from codon 2603 to 2688). **(b)** Structure of the most conserved domain of BRCA2 (green region). This domain wrapped around N-terminal of DSS1 (cyan protein). **(c)** Location of BRCA2 codon 2662 (red). Codon 2662 is at the center of the most conserved domain of BRCA2.

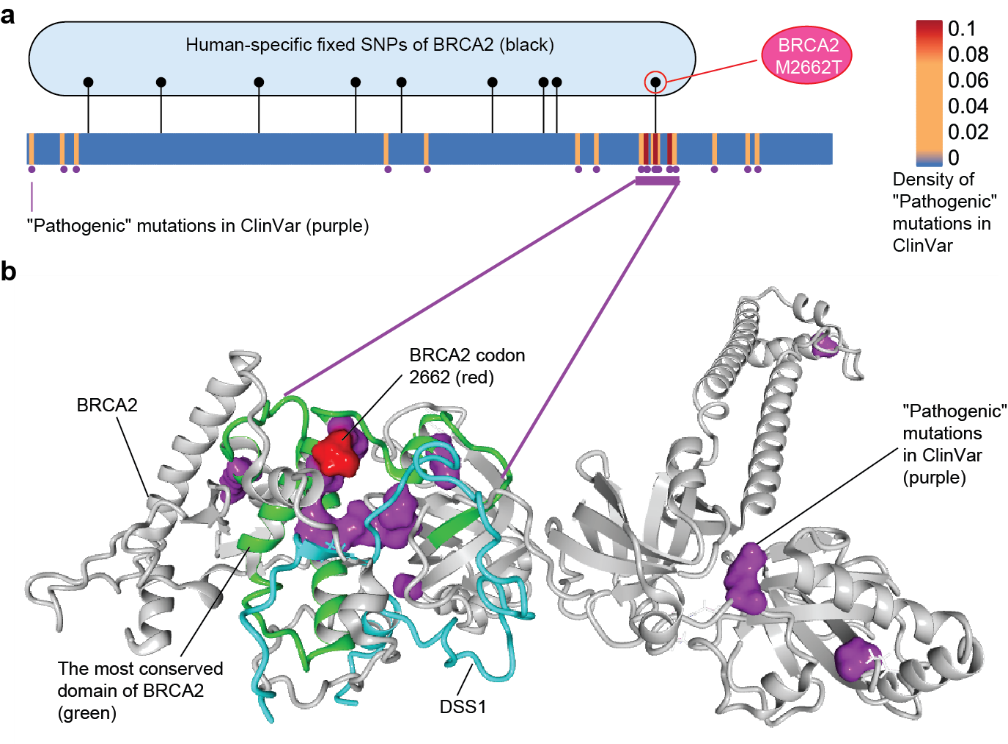

**Supplementary Fig. 8 | A comparison between human-specific fixed SNPs and known "Pathogenic" mutations in ClinVar of BRCA2.** **(a)** Nine human-specific fixed SNPs of BRCA2 (black dots) and 19 known "Pathogenic" mutations in ClinVar (purple dots). The purple line denotes the region with clustered "Pathogenic" mutations (n=9). **(b)** Structural view for BRCA2 codon 2662 (red), known "Pathogenic" mutations in ClinVar (purple dots) and the most conserved domain of BRCA2 (green). Codon 2662 is glued to the cluster of known "Pathogenic" mutations, located at the most conserved domain.

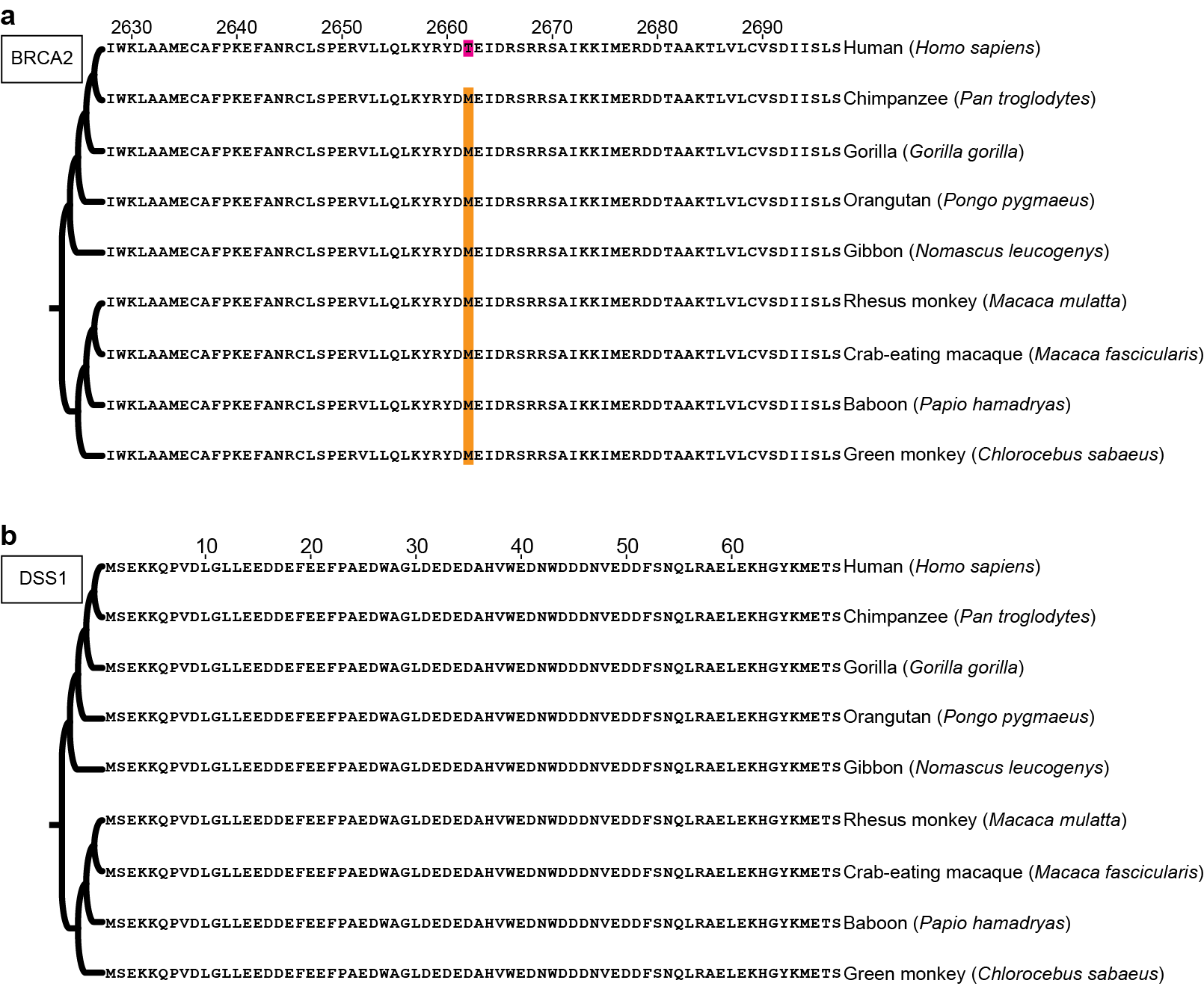

**Supplementary Fig. 9 | BRCA2^M2662T^ is the sole SNP found in BRCA2-DSS1 domain among great apes, lesser apes and old world monkeys.** **(a)** Alignment of a fragment of BRCA2 (from 2628 aa to 2697 aa). BRCA2^2662M^ was identified as the ancestral genotype because all non-human great apes, lesser apes and old world monkeys have M amino acid in the homologous position. BRCA2^2662T^ was the emergent genetic variant occurred in human after divergence from CHLCA. No other variants were found. **(b)** Alignment of whole protein sequence of DSS1. No variants were found in DSS1.

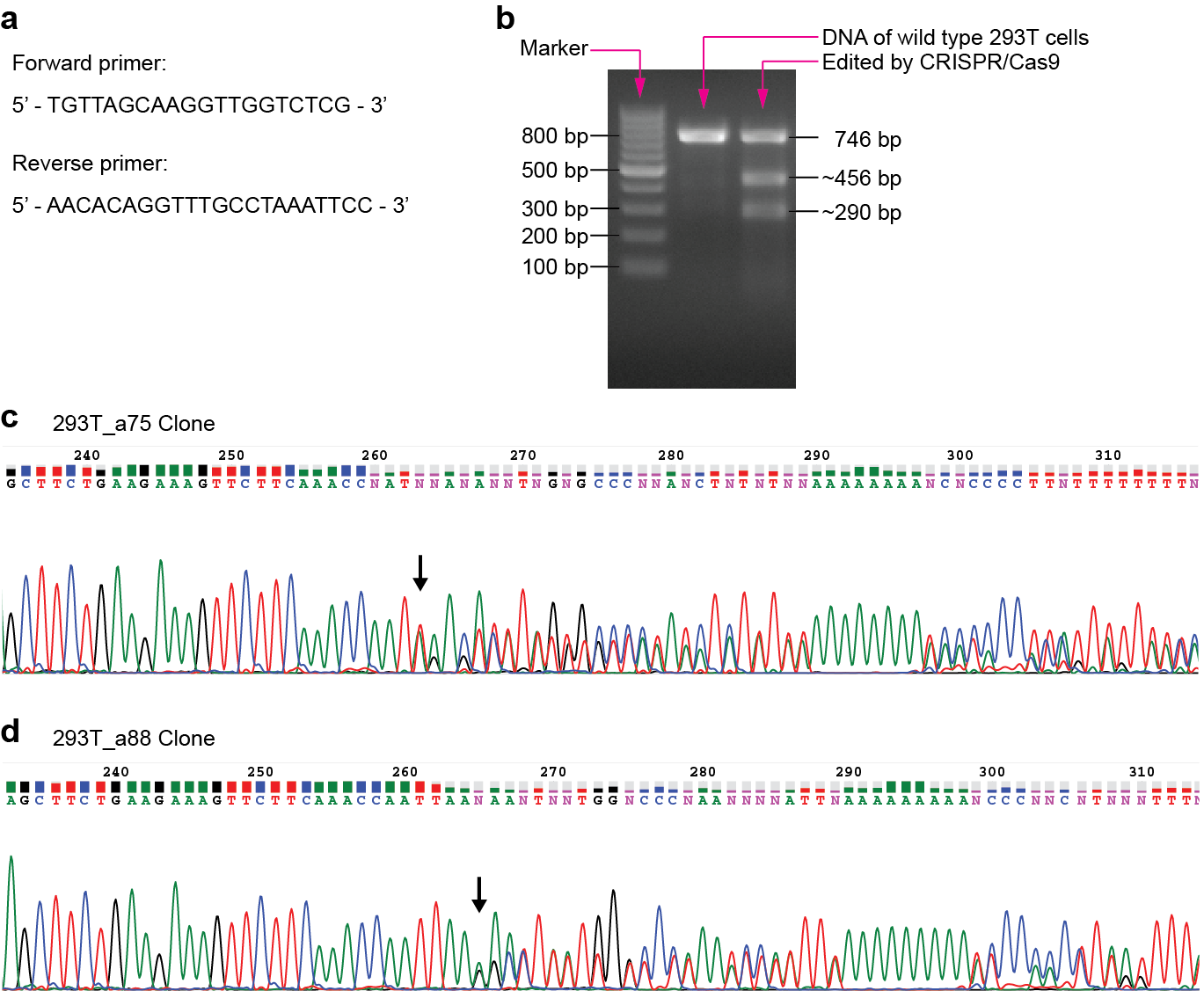

**Supplementary Fig. 10 | Identified positive clones after *BRCA2* gene editing.** **(a)** PCR primers used to amplify target fragment that covered the cut site. **(b)** Electropherogram after digest by T7 Endonuclease I (T7EI). **(c)** and **(d)** show the result of Sanger sequencing of 293T_a75 and 293T_a88 clones. The black arrow denotes the start of heterozygous peaks.

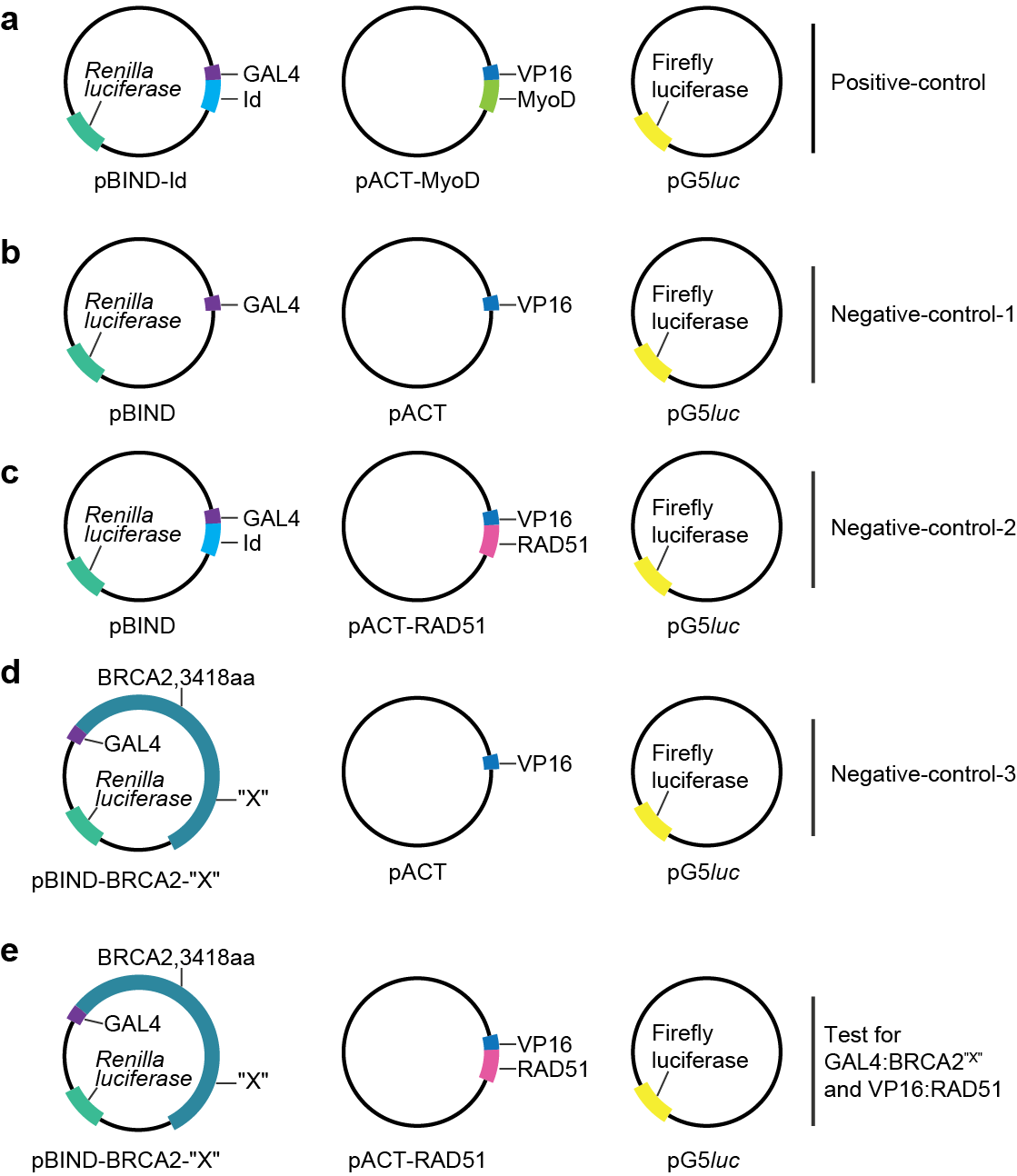

**Supplementary Fig. 11 | Schematic diagram for design of BRCA2-RAD51 two-hybrid test. (a)** The positive control reaction. The pBIND-Id and pACT-MyoD vectors encode GAL4:Id and VP16:MyoD fusion proteins, respectively. The binding of Id and MyoD lead to the binding of GAL4 and VP16, then activates the expression of pG5*luc*. The pG5*luc* vector expresses firefly luciferase. In addition, the pBIND-Id vector expresses the *Renilla* luciferase, which is used to normalize for differences in transfection efficiency. The pcDNA3.1(-) vector is used to maintain a constant mass of DNA for each transfection reaction. **(b)** Negative control: the background level of firefly luciferase expression from the pG5*luc* vector in the presence of the pACT and pBIND vectors. **(c)** Negative control: the background level of firefly luciferase expression from the pG5*luc* vector in the presence of the pACT-RAD51 and pBIND vectors. **(d)** Negative control: the background level of firefly luciferase expression from the pG5*luc* vector in the presence of the pACT and pBIND-BRCA2-"X" vectors. "X" denotes various mutations. **(e)** Tests for interaction of the GAL4:BRCA2^"X"^ fusion protein with the VP16:RAD51 fusion protein. "X" denotes various mutations.

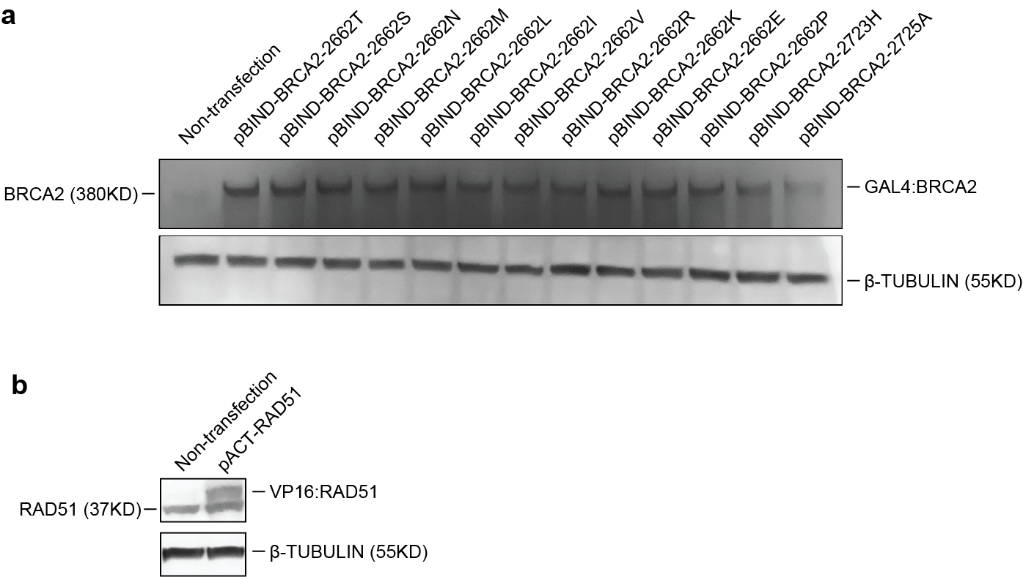

**Supplementary Fig. 12 | Expression of plasmids which were used in two-hybrid test. (a)** Expression of plasmids pBIND-BRCA2-"X". "X" denotes various mutations. 293T_a88 clone was used. **(b)** Expression of plasmid pACT-RAD51. 293T_a88 clone was used.

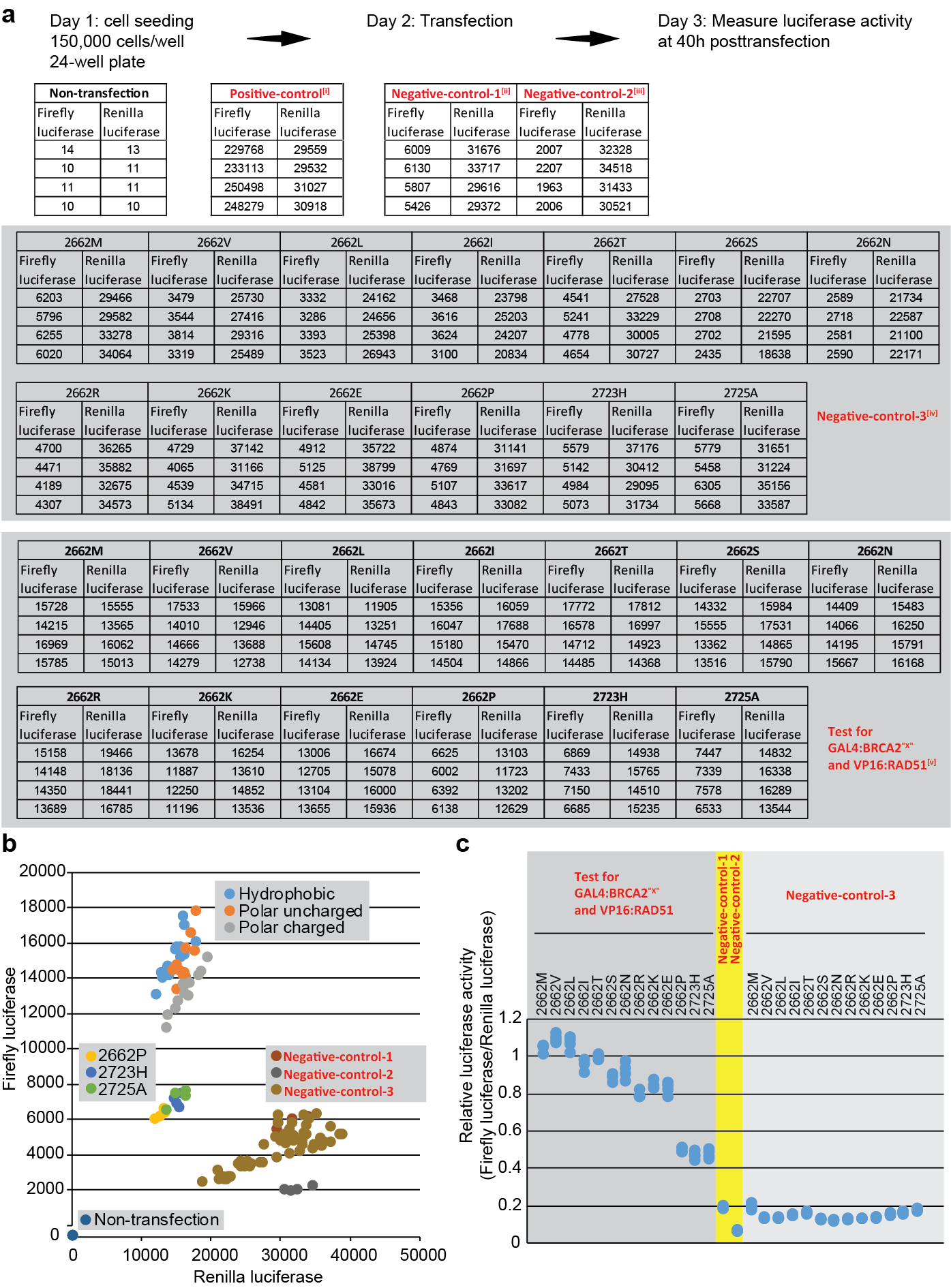

**Supplementary Fig. 13 | Test for BRCA2-RAD51 two-hybrid in 293T_a88 clone. (a)** Raw data of BRCA2-RAD51 two-hybrid test. The test included both positive and negative controls. [i] Positive-control, pBIND-Id + pACT-MyoD + pG5luc, related to Supplementary Fig. 11a. [ii] Negative-control-1, pBIND + pACT + pG5luc, related to Supplementary Fig. 11b. [iii] Negative-control-2, pBIND + pACT-RAD51 + pG5luc, related to Supplementary Fig. 11c. [iv] Negative-control-3, pBIND-BRCA2-"X" + pACT + pG5luc, related to Supplementary Fig. 11d. "X" denotes various mutations. [v] Test for GAL4:BRCA2^"X"^ and VP16:RAD51: pBIND-BRCA2-"X" + pACT-RAD51 + pG5luc, related to Supplementary Fig. 11e. "X" denotes various mutations. **(b)** A scatter plot showed very low firefly luciferase activity in the negative control group. **(c)** Relative luciferase activity of BRCA2-RAD51 two-hybrid. Positive-control and Non-transfection were not included.

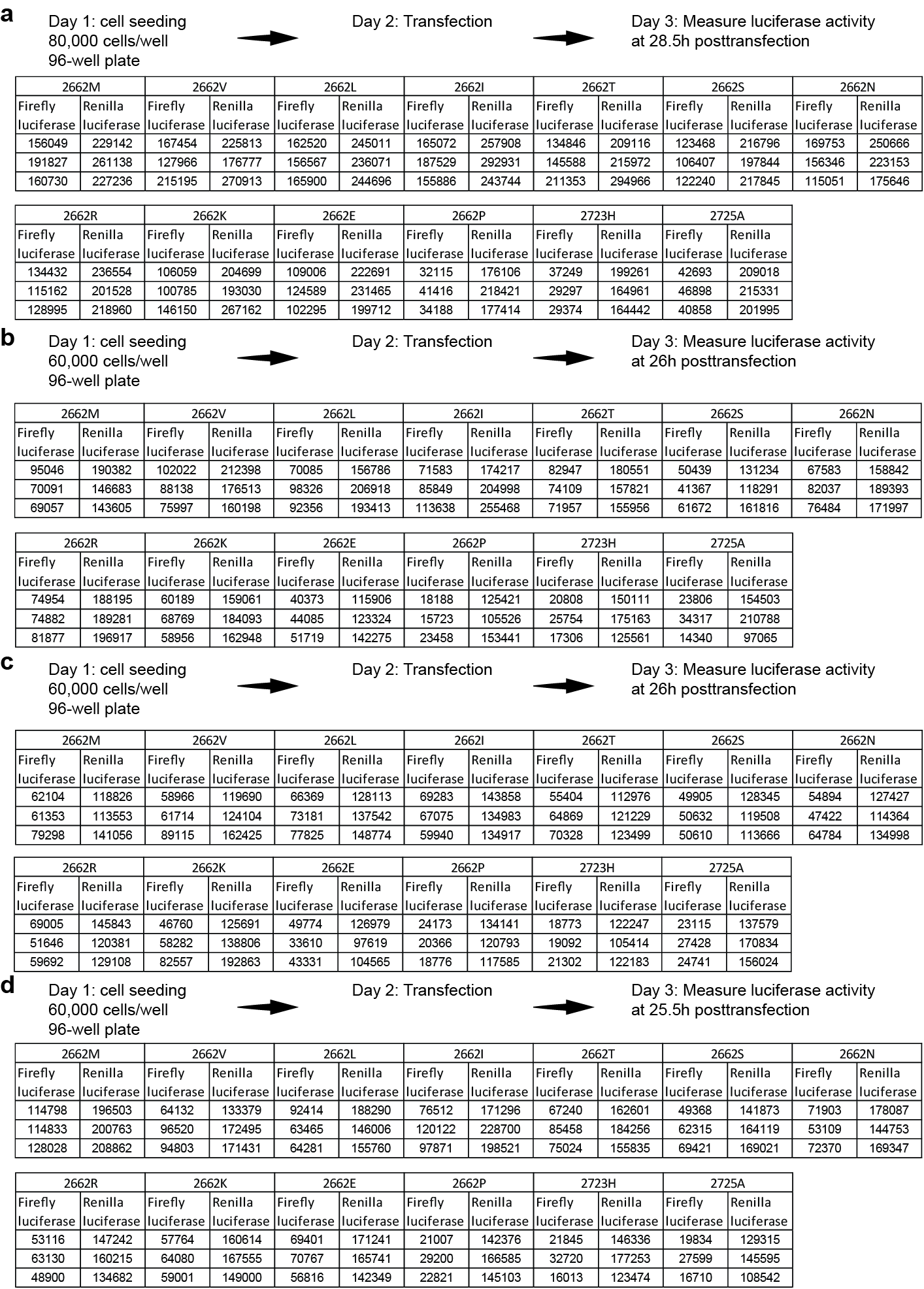

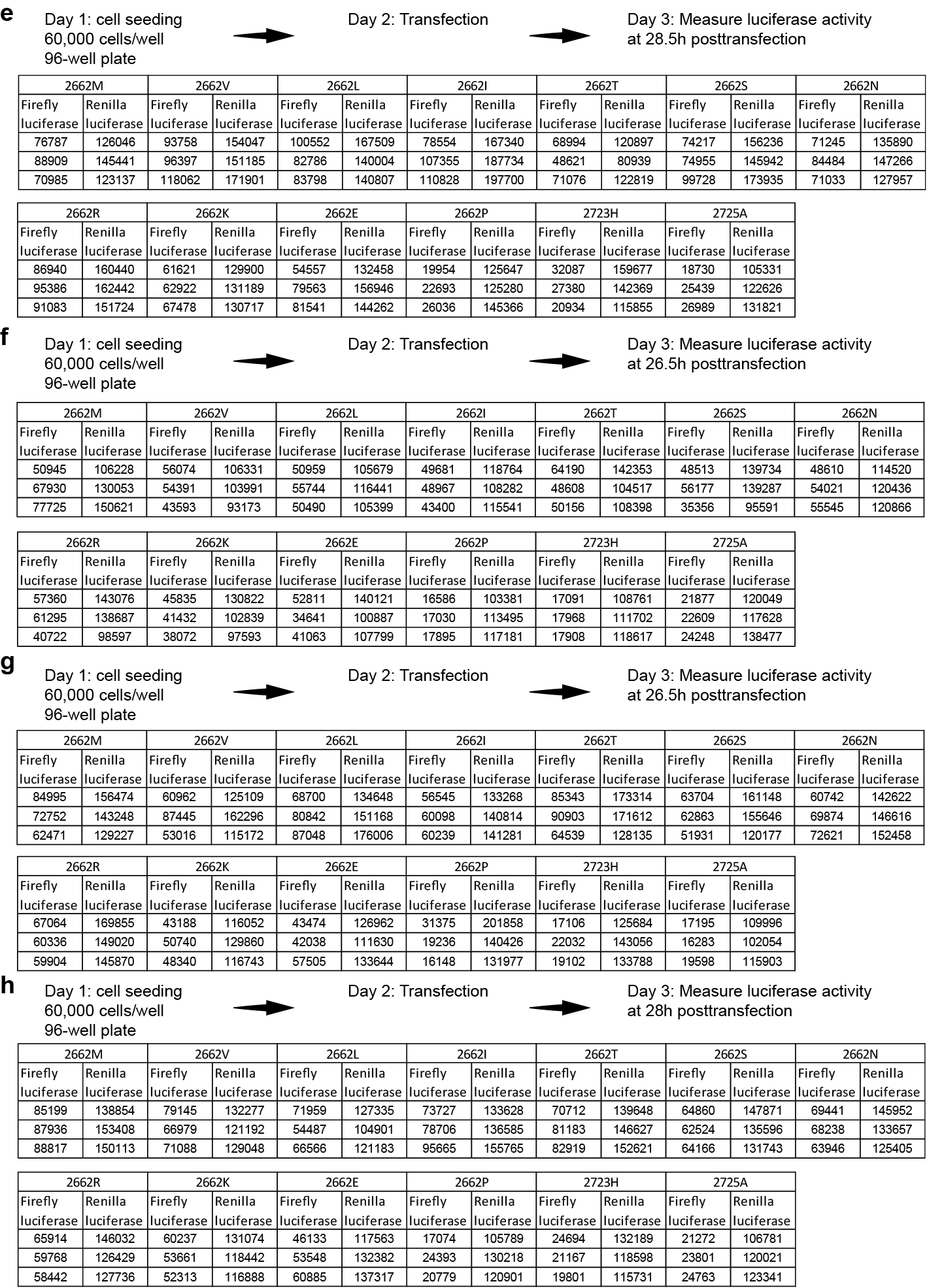

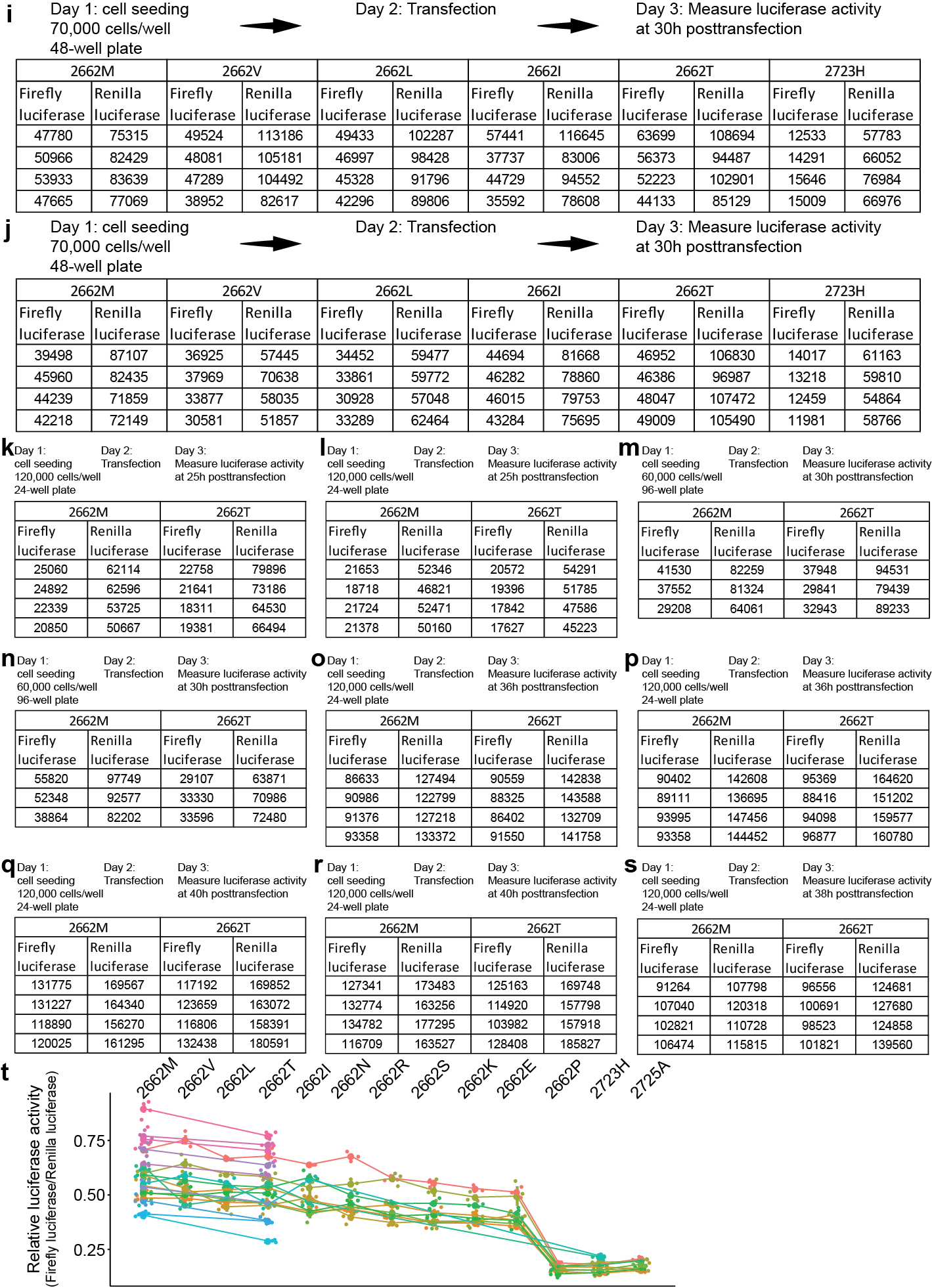

**Supplementary Fig. 14 | Result of BRCA2-RAD51 two-hybrid in 293T_a88 clone. (a-s)** Raw data of BRCA2-RAD51 two-hybrid in 293T_a88 clone. **(t)** Relative luciferase activity of 13 BRCA2 mutations.

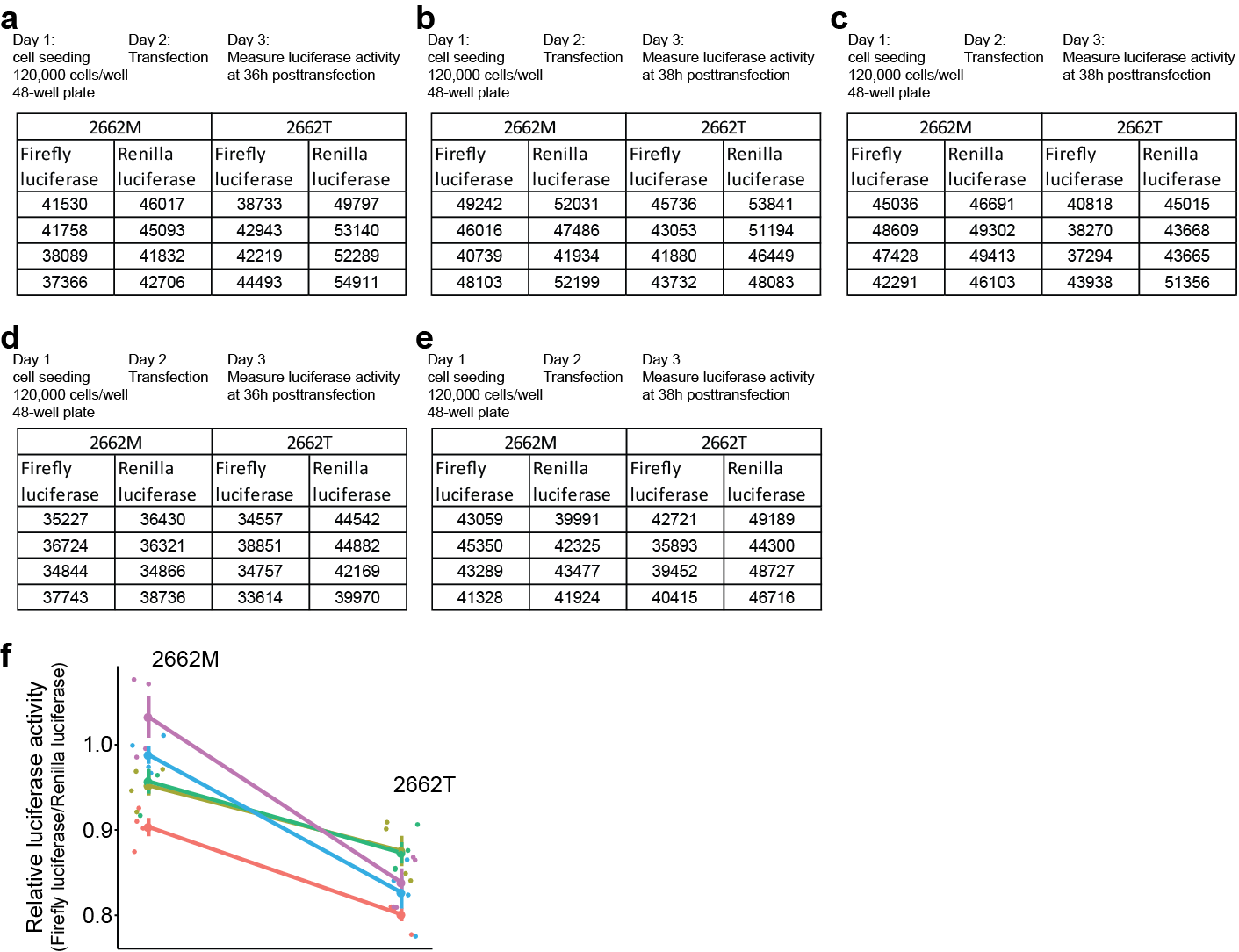

**Supplementary Fig. 15 | Result of BRCA2-RAD51 two-hybrid in 293T_a75 clone. (a-e)** Raw data of BRCA2-RAD51 two-hybrid in 293T_a75 clone. **(f)** Relative luciferase activity of 2 BRCA2 mutations.

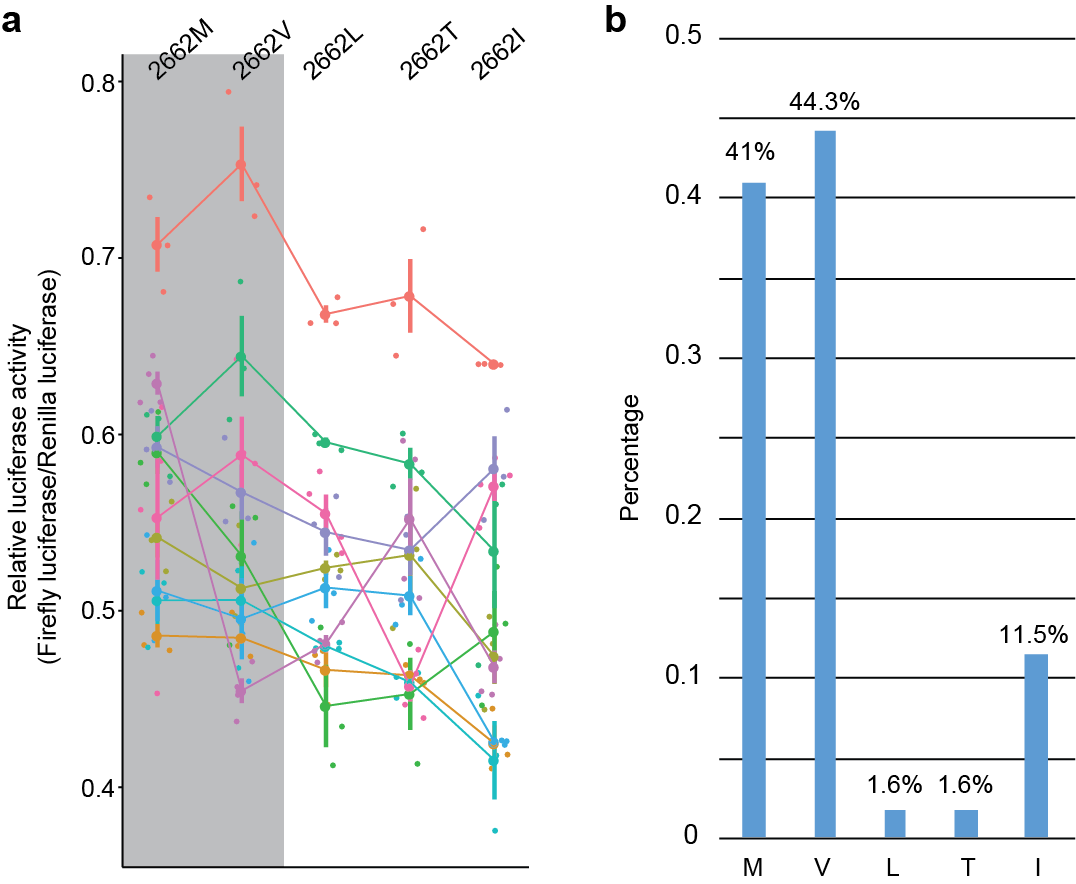

**Supplementary Fig. 16 | BRCA2 activity versus amino acid frequency. (a)** Relative luciferase activity of BRCA2-RAD51 two-hybrid with the five different amino acids within the site of BRCA2 codon 2662. Data from Supplementary Fig. 14 a-j were used. **(b)** The frequency of five amino acids in mammals. Related to Fig. 2b.

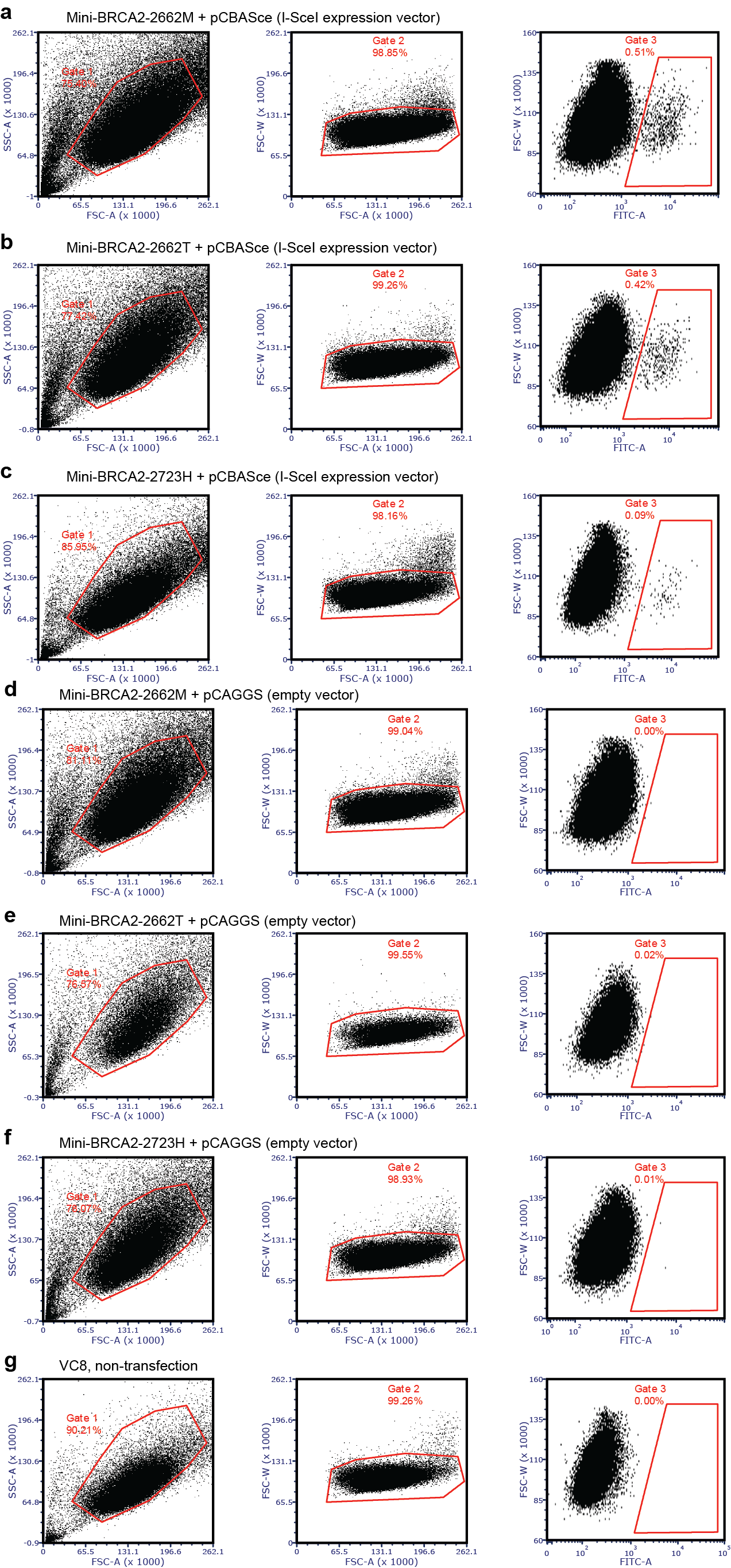

**Supplementary Fig. 17 | The first flow cytometric analysis of V-C8 DR-GFP cells to detect cellular green fluorescence following DSB induction.** V-C8 cells containing a chromosomal DR-GFP reporter were cotransfected with expression vectors for the I-SceI endonuclease and the mini-BRCA2. V-C8 cells were assayed by flow cytometry 48h after transfection. **(a-c)** Result of flow cytometric analysis of V-C8 cells demonstrates decreased DSB repair after transient expression of mini-BRCA2^2662T^ as compared with mini-BRCA2^2662M^. Related to Fig. 4 c-f. **(d-g)** Negative controls. GFP-positive cells were rare in the absence of I-SceI expression.

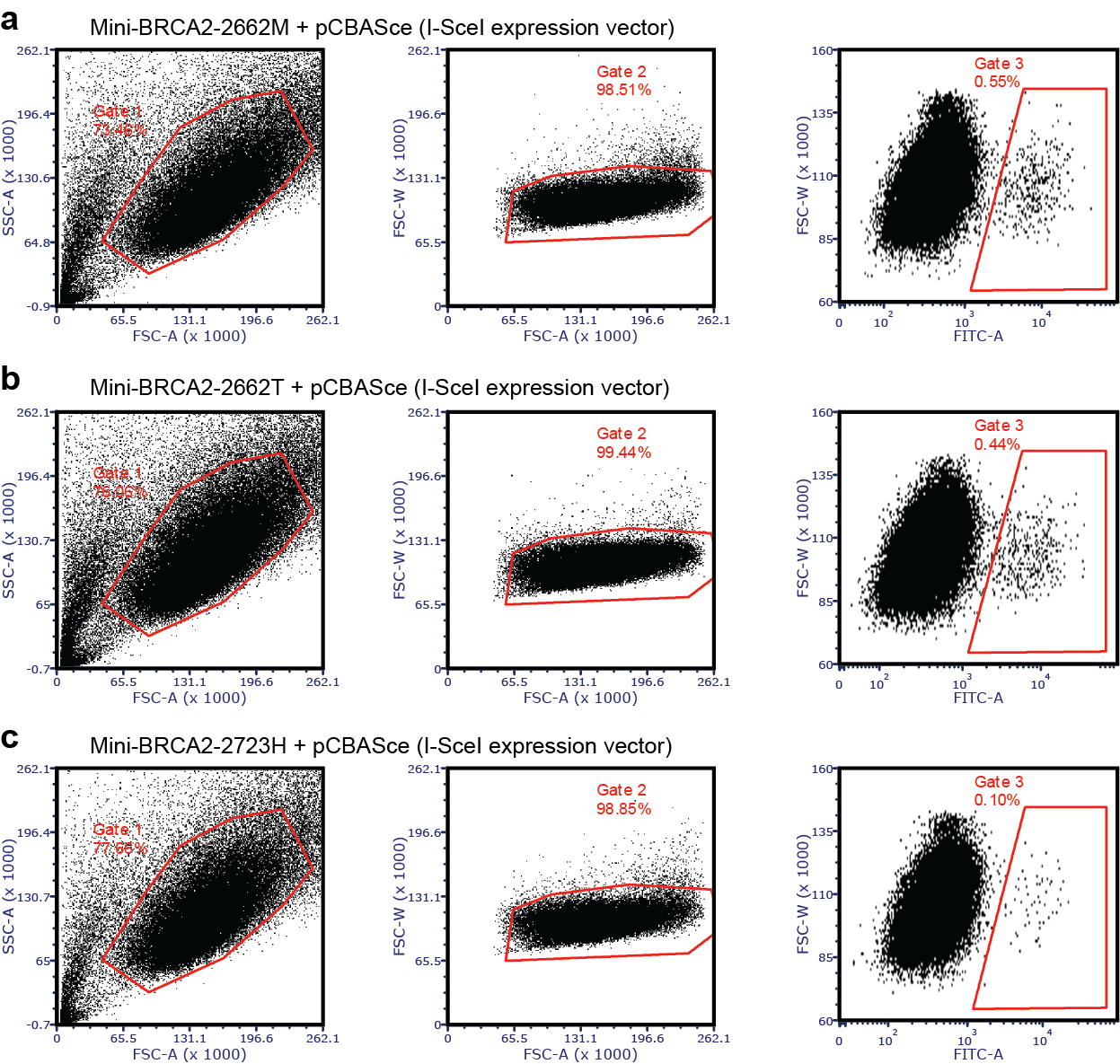

**Supplementary Fig. 18 | The second flow cytometric analysis of V-C8 DR-GFP cells to detect cellular green fluorescence following DSB induction.** Related to Fig. 4 f.

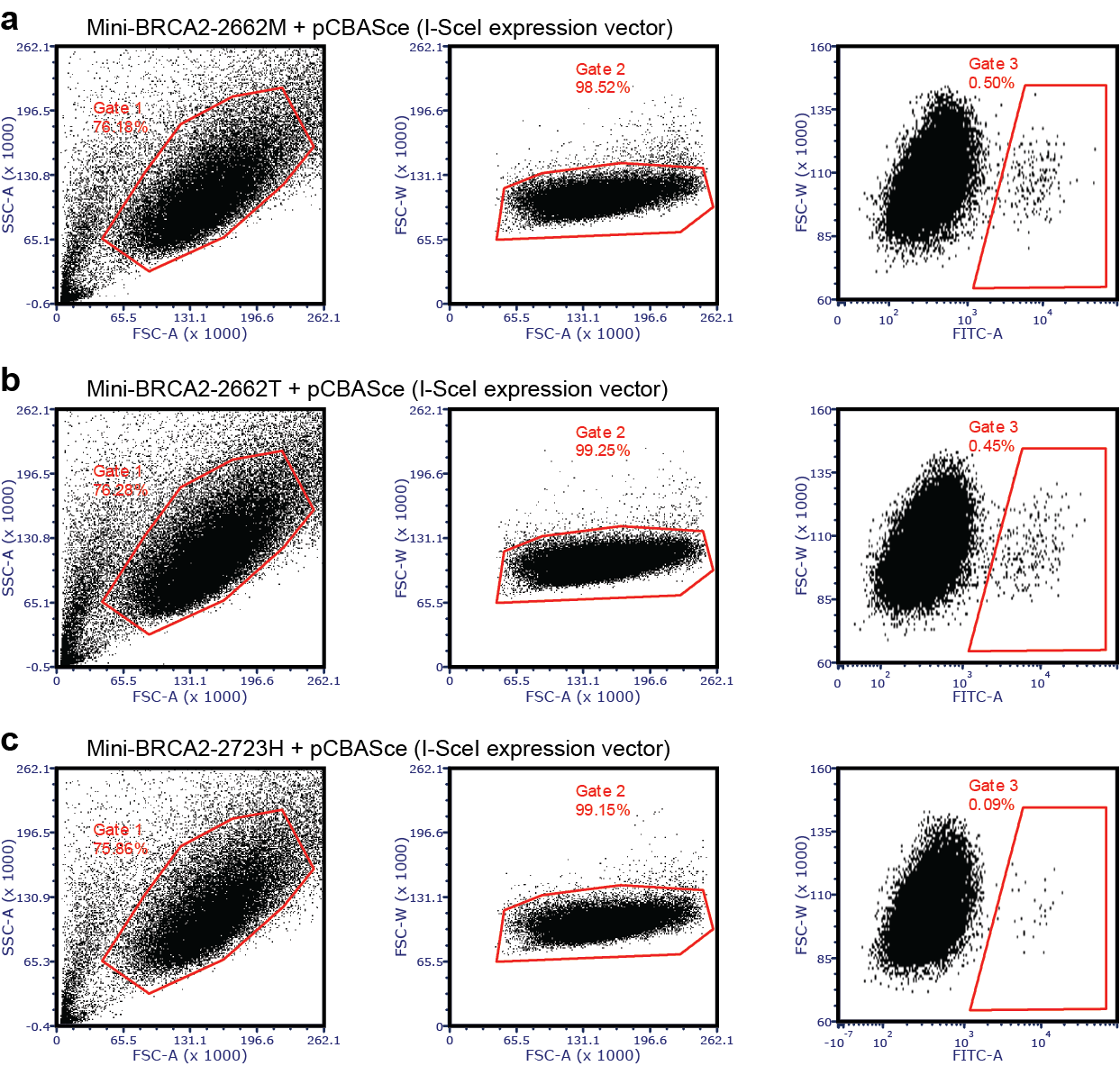

**Supplementary Fig. 19 | The third flow cytometric analysis of V-C8 DR-GFP cells to detect cellular green fluorescence following DSB induction.** Related to Fig. 4 f.

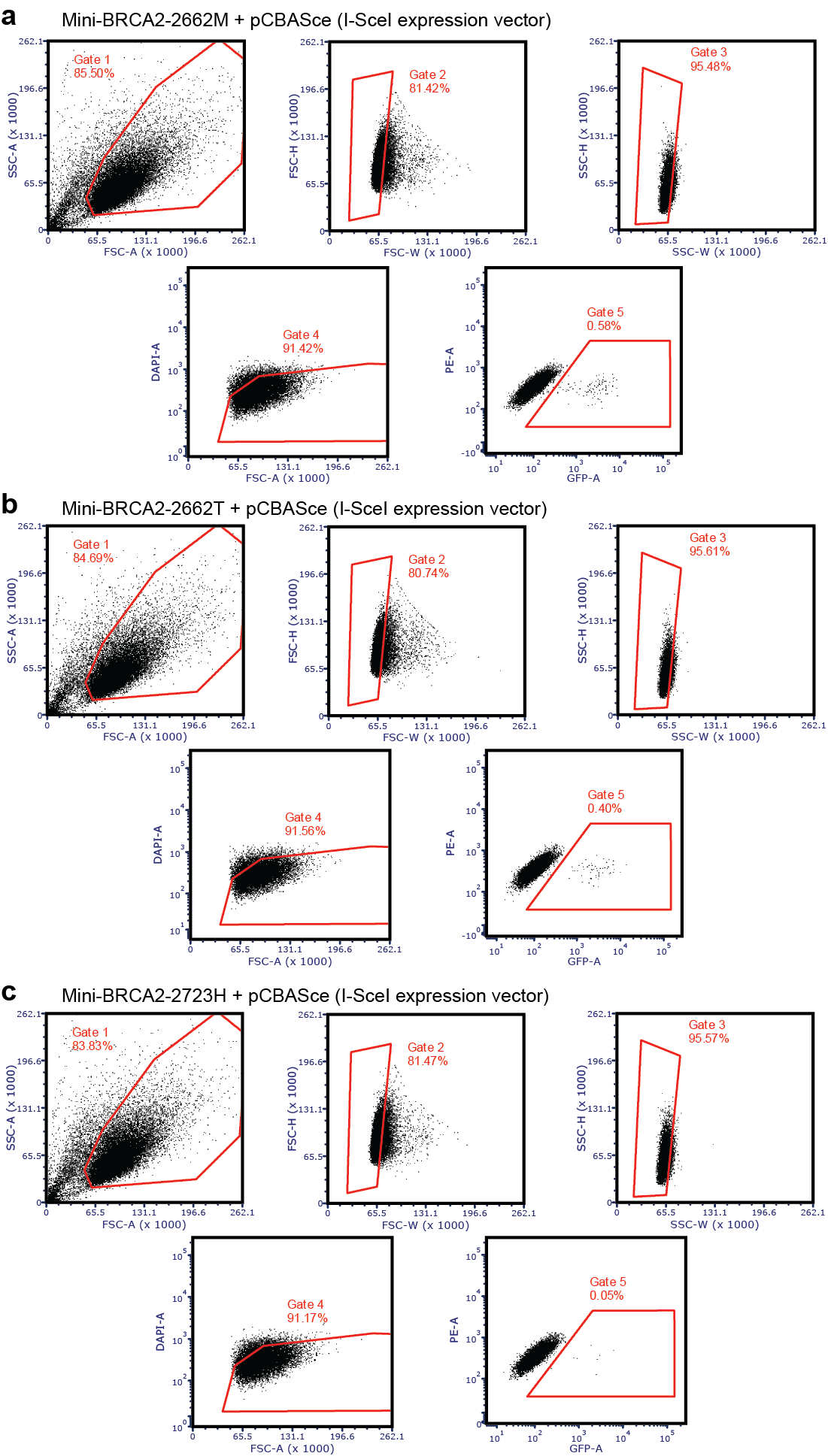

**Supplementary Fig. 20 | The fourth flow cytometric analysis of V-C8 DR-GFP cells to detect cellular green fluorescence following DSB induction.** Related to Fig. 4 f.

**Supplementary Table 1.**

Detailed information for the data of 21 non-human primate genomes, 54 ancient human genomes and 2 present-day human genomes used in this study. Estimated times for ancient human genomes refer to Slatkin et al.^24^.

| **Common Name used in this study** | **Scientific Name** | **BioSample**  (and estimated times for ancient human genomes) | **BioProject** | **Run ID** |
| --- | --- | --- | --- | --- |
| Chimpanzee_1 | *Pan troglodytes* | SAMN01920535 | PRJNA189439 | SRR748066; SRR748067; SRR748068; SRR748069; SRR748070 |
| Chimpanzee_2 | *Pan troglodytes* | SAMN01920536 | PRJNA189439 | SRR748071; SRR748072; SRR748073; SRR748074; SRR748075 |
| Bonobo_1 | *Pan paniscus* | SAMN01920507 | PRJNA189439 | SRR740781; SRR740782; SRR740783; SRR740784; SRR740787; SRR740790; SRR740792; SRR740793 |
| Bonobo_2 | *Pan paniscus* | SAMN01920508 | PRJNA189439 | SRR740794; SRR740795; SRR740796; SRR740797; SRR740798; SRR740799; SRR740800; SRR740801 |
| Gorilla_1 | *Gorilla gorilla* | SAMN01920476 | PRJNA189439 | SRR748109; SRR748110; SRR748111; SRR748112 |
| Gorilla_2 | *Gorilla gorilla* | SAMN01920489 | PRJNA189439 | SRR748076; SRR748077; SRR748078; SRR748079; SRR748080 |
| Orangutan_1 | *Pongo abelii* | SAMN01920545 | PRJNA189439 | SRR748041; SRR748042; SRR748044; SRR748045 |
| Orangutan _2 | *Pongo pygmaeus* | SAMN01920547 | PRJNA189439 | SRR747999; SRR748000; SRR748001; SRR748002; SRR748003 |
| Gibbon_1 | *Hylobates moloch* | SAMN02850874 | PRJNA232723 | SRR1390886; SRR1390887; SRR1390888; SRR1390889; SRR1390890; SRR1390891; SRR1390892; SRR1390893; SRR1390894; SRR1390895; SRR1390896; SRR1390897; SRR1390898; SRR1390899; SRR1390900; SRR1390901; SRR1390902; SRR1390903; SRR1390904; SRR1390905; SRR1390906; SRR1390907; SRR1390908; SRR1390909; SRR1390910; SRR1390911; SRR1390912; SRR1390913; SRR1390914; SRR1390915; SRR1390916; SRR1390917; SRR1390918; SRR1390919; SRR1390920; SRR1390921; SRR1390922; SRR1390923; SRR1390924; SRR1390925; SRR1390926; SRR1390927; SRR1390928; SRR1390929; SRR1390930; SRR1390931; SRR1390932; SRR1390933; SRR1390934; SRR1390935; SRR1390936; SRR1390937 |
| Gibbon_2 | *Symphalangus syndactylus* | SAMN02388245 | PRJNA232723 | SRR1390831; SRR1390832; SRR1390833; SRR1390834; SRR1390835; SRR1390852; SRR1390869 |
| Macaque_1 | *Macaca mulatta* | SAMN03200174 | PRJNA239805 | SRR1790406; SRR1790407; SRR1790409; SRR1790411; SRR1790412; SRR1790413; SRR1790415; SRR1790416 |
| Macaque_2 | *Macaca mulatta* | SAMN03200175 | PRJNA239805 | SRR1790419; SRR1790422; SRR1790423; SRR1790424; SRR1790425; SRR1790427; SRR1790428; SRR1790429 |
| Snub-nosed monkey_1 | *Rhinopithecus strykeri* | SAMN03076575 | PRJNA261768 | SRR1588563; SRR1588566 |
| Snub-nosed monkey_2 | *Rhinopithecus rolellana* | SAMN03076573 | PRJNA261768 | SRR1588550; SRR1588560; SRR1588561 |
| Marmoset_1 | *Callithrix jacchus* | SAMN02744554 | PRJNA20401 | SRR1274859; SRR1274860; SRR1274861; SRR1274862 |
| Marmoset_2 | *Callithrix jacchus* | SAMN02744555 | PRJNA20401 | SRR1274963; SRR1274964; SRR1274965; SRR1274966 |
| Squirrel monkey | *Saimiri boliviensis* | SAMN00672664 | PRJNA67945 | SRR315550; SRR315552; SRR315553; SRR315556; SRR315560 |
| Tarsius | *Carlito syrichta* | SAMN04994160 | PRJNA20339 | SRR3502859; SRR3502861; SRR3502863; SRR3502864; SRR3502866; SRR3502867; SRR3502868; SRR3502869; SRR3502870; SRR3502871; SRR3502917; SRR3502921; SRR3922571; SRR3922572; SRR3922616; SRR3922617; SRR3922618; SRR3922619; SRR3922620; SRR3922621; SRR3922622; SRR3922623; SRR3922625; SRR3922626 |
| Bushbaby | *Otolemur garnettii* | SAMN00103053 | PRJNA16955 | SRR034397; SRR034398; SRR034399; SRR034400; SRR034401; SRR034402; SRR034403; SRR034404; SRR065344; SRR065345; SRR065346; SRR065347; SRR065348; SRR065349; SRR065350; SRR065351; SRR065352; SRR065353; SRR065354; SRR065355; SRR065356; SRR065357; SRR065358; SRR065359; SRR065360; SRR065361 |
| Mouse Lemur_1 | *Microcebus murinus* | SAMN02900033 | PRJNA19967 | SRR1537460; SRR1537471 |
| Mouse Lemur_2 | *Microcebus murinus* | SAMN02900024 | PRJNA19967 | SRR1537461; SRR1537462; SRR1537463 |
| Neanderthal | *Homo sapiens neanderthalensis* | SAMEA2052074  (50000-70000 years bp) | PRJEB1265 | ERR229910; ERR229911; ERR229912; ERR229913; ERR229914 |
| Denisovan | *Homo sapiens denisova* | SAMEA1487354  (80000 years bp) | PRJEB3092 | ERR141688; ERR141689; ERR141690; ERR141691; ERR141692; ERR141693; ERR141694; ERR141695; ERR141696; ERR141697; ERR141698; ERR141699; ERR141700 |
| Ust_Ish | *Homo sapiens* | SAMEA2656891  (45000 years bp) | PRJEB6622 | ERR566093 |
| Kotias | *Homo sapiens* | SAMEA3609088  (9529-9895 years bp) | PRJEB11364 | ERR1078321; ERR1078322; ERR1078323; ERR1078324; ERR1078325 |
| Satsurblia | *Homo sapiens* | SAMEA3609089  (13132-13380 years bp) | PRJEB11364 | ERR1078326; ERR1078327; ERR1078328; ERR1078329; ERR1078330 |
| Bichon | *Homo sapiens* | SAMEA3609090  (13560-13770 years bp) | PRJEB11364 | ERR1078331; ERR1078332; ERR1078333; ERR1078334; ERR1078335; ERR1078336; ERR1078337; ERR1078338; ERR1078339; ERR1078340; ERR1078341; ERR1078342; ERR1078343; ERR1078344; ERR1078345; ERR1078346; ERR1078347; ERR1078348; ERR1078349; ERR1078350; ERR1078351 |
| Bar_Ear_1 | *Homo sapiens* | SAMEA3672571  (8200-8400 years bp) | PRJEB11848 | ERR1144005 |
| Bar_Ear_2 | *Homo sapiens* | SAMEA3672572  (8000-8200 years bp) | PRJEB11848 | ERR1144001 |
| Kle_Lat | *Homo sapiens* | SAMEA3672573  (6000-6200 years bp) | PRJEB11848 | ERR1352893 |
| Pal_Lat | *Homo sapiens* | SAMEA3672574  (~6400 years bp) | PRJEB11848 | ERR1144003 |
| Rev_Ear | *Homo sapiens* | SAMEA3672575  (8200-8500 years bp) | PRJEB11848 | ERR1144004 |
| Motala | *Homo sapiens* | SAMEA2697132  (8000 years bp) | PRJEB6272 | ERR577498 |
| Stuttgart | *Homo sapiens* | SAMEA2697125  (7000 years bp) | PRJEB6272 | ERR577491 |
| Loschbour | *Homo sapiens* | SAMEA2697124  (8000 years bp) | PRJEB6272 | ERR577490 |
| Cov_Bon | *Homo sapiens* | SAMN03470461  (7400 years bp) | PRJNA280812 | SRR1964490 |
| La_Bra | *Homo sapiens* | SAMN02437317  (7000 years bp) | PRJNA230689 | SRR1045122; SRR1045127 |
| Lat_Bro | *Homo sapiens* | SAMN02680267  (3100-3300 years bp) | PRJNA240906 | SRR1186791 |
| Lat_Cop | *Homo sapiens* | SAMN02687908  (4700-4900 years bp) | PRJNA240906 | SRR1187905 |
| Iro_Pre | *Homo sapiens* | SAMN02680194  (2800-3000 years bp) | PRJNA240906 | SRR1186714 |
| Ear_Neo | *Homo sapiens* | SAMN02680193  (7600-7800 years bp) | PRJNA240906 | SRR1187910 |
| Mid_Neo_Alf | *Homo sapiens* | SAMN02680266  (7000-7300 years bp) | PRJNA240906 | SRR1186790 |
| Mid_Neo_Lat | *Homo sapiens* | SAMN02687905  (7000-7200 years bp) | PRJNA240906 | SRR1187897 |
| Mid_LBK | *Homo sapiens* | SAMN02687906  (7000-7300 years bp) | PRJNA240906 | SRR1187912 |
| Lat_Neo | *Homo sapiens* | SAMN02687907  (6400-6500 years bp) | PRJNA240906 | SRR1187902 |
| Mota | *Homo sapiens* | SAMN04115941  (4500 years bp) | PRJNA295861 | SRR2544592 |
| Iri_Bal | *Homo sapiens* | SAMEA3696307  (5000-5400 years bp) | PRJEB11995 | ERR1162625; ERR1162626; ERR1162627; ERR1162628; ERR1162629; ERR1162630; ERR1162631; ERR1162632; ERR1162633; ERR1162634 |
| Iri_Rat_1 | *Homo sapiens* | SAMEA3696317  (~4000 years bp) | PRJEB11995 | ERR1162635; ERR1162636; ERR1162637; ERR1162638; ERR1162639; ERR1162640; ERR1162641; ERR1162642; ERR1162643; ERR1162644 |
| Iri_Rat_2 | *Homo sapiens* | SAMEA3696327  (3700-4000 years bp) | PRJEB11995 | ERR1162645; ERR1162646; ERR1162647; ERR1162648 |
| Iri_Rat_3 | *Homo sapiens* | SAMEA3696331  (3500-3700 years bp) | PRJEB11995 | ERR1162649; ERR1162650; ERR1162651; ERR1162652 |
| Bat_Afa | *Homo sapiens* | ERS699134  (3000-5000 years bp) | PRJEB9021 | ERR844313 |
| Kyt_And_1 | *Homo sapiens* | ERS699125  (3000-5000 years bp) | PRJEB9021 | ERR844304 |
| Kyt_And_2 | *Homo sapiens* | ERS699129  (3000-5000 years bp) | PRJEB9021 | ERR844308 |
| Bed_Nor | *Homo sapiens* | ERS699073  (3000-5000 years bp) | PRJEB9021 | ERR844252 |
| Qxie_Iro | *Homo sapiens* | ERS699079  (3000-5000 years bp) | PRJEB9021 | ERR844258 |
| Kyt_Iro | *Homo sapiens* | ERS699128  (3000-5000 years bp) | PRJEB9021 | ERR844307 |
| Ver_Iro | *Homo sapiens* | ERS699164  (3000-5000 years bp) | PRJEB9021 | ERR844343 |
| Sar_Iro | *Homo sapiens* | ERS699165  (3000-5000 years bp) | PRJEB9021 | ERR844344 |
| Sab_Kar | *Homo sapiens* | ERS699119  (3000-5000 years bp) | PRJEB9021 | ERR844298 |
| Arb_Kar_1 | *Homo sapiens* | ERS699121  (3000-5000 years bp) | PRJEB9021 | ERR844300 |
| Arb_Kar_2 | *Homo sapiens* | ERS699122  (3000-5000 years bp) | PRJEB9021 | ERR844301 |
| Arb_Kar_3 | *Homo sapiens* | ERS699123  (3000-5000 years bp) | PRJEB9021 | ERR844302 |
| Bys_Kar_1 | *Homo sapiens* | ERS699124  (3000-5000 years bp) | PRJEB9021 | ERR844303 |
| Bys_Kar_2 | *Homo sapiens* | ERS699126  (3000-5000 years bp) | PRJEB9021 | ERR844305 |
| Kap_Mez | *Homo sapiens* | ERS699138  (3000-5000 years bp) | PRJEB9021 | ERR844317 |
| Bol_Sin | *Homo sapiens* | ERS699096  (3000-5000 years bp) | PRJEB9021 | ERR844275 |
| Prz_Une | *Homo sapiens* | ERS699077  (3000-5000 years bp) | PRJEB9021 | ERR844256 |
| Erd_Vat | *Homo sapiens* | ERS699111  (3000-5000 years bp) | PRJEB9021 | ERR844290 |
| Ulan_Yam | *Homo sapiens* | ERS699145  (3000-5000 years bp) | PRJEB9021 | ERR844324 |
| Eng_Rom_1 | *Homo sapiens* | SAMEA3587709  (~2000 years bp) | PRJEB11004 | ERR1043141; ERR1043142; ERR1043143; ERR1043144 |
| Eng_Rom_2 | *Homo sapiens* | SAMEA3587710  (~2000 years bp) | PRJEB11004 | ERR1043145; ERR1043146; ERR1043147 |
| Eng_Rom_3 | *Homo sapiens* | SAMEA3587711  (~2000 years bp) | PRJEB11004 | ERR1043148; ERR1043149; ERR1043150 |
| Eng_Rom_4 | *Homo sapiens* | SAMEA3587712  (~2000 years bp) | PRJEB11004 | ERR1043151; ERR1043152; ERR1043153 |
| Eng_Rom_5 | *Homo sapiens* | SAMEA3587714  (~2000 years bp) | PRJEB11004 | ERR1043157; ERR1043158; ERR1043159 |
| Eng_Ang | *Homo sapiens* | SAMEA3587716  (~2000 years bp) | PRJEB11004 | ERR1043160; ERR1043161; ERR1043162 |
| Present-day_human_1 | *Homo sapiens* | SAMN00801888 | PRJEB2890 | ERR091571; ERR091572; ERR091573 |
| Present-day_human_2 | *Homo sapiens* | SAMN00001584 | PRJEB2378 | ERR024187; ERR024188; ERR024189; ERR024190; ERR024191; ERR024192; ERR024193; ERR024194; ERR024195; ERR024196; ERR024197; ERR024198; ERR024199; ERR024200; ERR024201; ERR024202 |

**Supplementary Table 2.**

Result of mapping, SNPs calling and non-silent SNPs for each genome.

| **Common Name used in this study** | **Scientific Name** | **Mapping** | **All SNPs** | **Non-silent SNPs** |
| --- | --- | --- | --- | --- |
| Chimpanzee_1 | *Pan troglodytes* | 89.53% | 37810545 | 82217 |
| Chimpanzee_2 | *Pan troglodytes* | 87.12% | 36703543 | 75746 |
| Bonobo_1 | *Pan paniscus* | 87.63% | 36776962 | 74018 |
| Bonobo_2 | *Pan paniscus* | 89.90% | 36904448 | 75062 |
| Gorilla_1 | *Gorilla gorilla* | 86.71% | 44190742 | 80378 |
| Gorilla_2 | *Gorilla gorilla* | 84.64% | 46197570 | 87468 |
| Orangutan_1 | *Pongo abelii* | 82.41% | 80616502 | 146302 |
| Orangutan_2 | *Pongo pygmaeus* | 82.74% | 81300268 | 153891 |
| Gibbon_1 | *Hylobates moloch* | 84.11% | 94803953 | 261874 |
| Gibbon_2 | *Symphalangus syndactylus* | 74.28% | 96499907 | 265163 |
| Macaque_1 | *Macaca mulatta* | 31.21% | 93180671 | 361043 |
| Macaque_2 | *Macaca mulatta* | 31.32% | 92011431 | 360063 |
| Snub-nosed monkey_1 | *Rhinopithecus strykeri* | 47.31% | 110920511 | 344118 |
| Snub-nosed monkey_2 | *Rhinopithecus rolellana* | 48.56% | 117746595 | 364849 |
| Marmoset_1 | *Callithrix jacchus* | 13.84% | 36508769 | 267110 |
| Marmoset_2 | *Callithrix jacchus* | 12.92% | 24386949 | 185005 |
| Squirrel monkey | *Saimiri boliviensis* | 13.79% | 43698865 | 323269 |
| Tarsius | *Carlito syrichta* | 1.11% | 5684918 | 126587 |
| Bushbaby | *Otolemur garnettii* | 1.20% | 3023913 | 107328 |
| Mouse Lemur_1 | *Microcebus murinus* | 2.12% | 6489696 | 153442 |
| Mouse Lemur_2 | *Microcebus murinus* | 2.07% | 5007888 | 166707 |
| Neanderthal | *Homo sapiens neanderthalensis* | 99.80% | 5543402 | 15885 |
| Denisovan | *Homo sapiens denisova* | 73.92% | 5835869 | 17574 |
| Ust_Ish | *Homo sapiens* | 99.79% | 4418127 | 12827 |
| Kotias | *Homo sapiens* | 82.60% | 3902096 | 18382 |
| Satsurblia | *Homo sapiens* | 19.49% | 249853 | 625 |
| Bichon | *Homo sapiens* | 81.38% | 7266729 | 43379 |
| Bar_Ear_1 | *Homo sapiens* | 99.93% | 2343729 | 6919 |
| Bar_Ear_2 | *Homo sapiens* | 99.97% | 1213591 | 3078 |
| Kle_Lat | *Homo sapiens* | 99.76% | 587247 | 1768 |
| Pal_Lat | *Homo sapiens* | 99.77% | 246416 | 712 |
| Rev_Ear | *Homo sapiens* | 99.98% | 227506 | 630 |
| Motala | *Homo sapiens* | 99.92% | 702077 | 2755 |
| Stuttgart | *Homo sapiens* | 99.76% | 3860456 | 11690 |
| Loschbour | *Homo sapiens* | 99.79% | 3721844 | 10348 |
| Cov_Bon | *Homo sapiens* | 99.25% | 175250 | 1771 |
| La_Bra | *Homo sapiens* | 99.95% | 1219597 | 6162 |
| Lat_Bro | *Homo sapiens* | 99.95% | 3877871 | 11386 |
| Lat_Cop | *Homo sapiens* | 99.98% | 124731 | 372 |
| Iro_Pre | *Homo sapiens* | 99.97% | 184973 | 476 |
| Ear_Neo | *Homo sapiens* | 99.96% | 185428 | 648 |
| Mid_Neo_Alf | *Homo sapiens* | 99.95% | 3825197 | 11178 |
| Mid_Neo_Lat | *Homo sapiens* | 99.97% | 99387 | 284 |
| Mid_LBK | *Homo sapiens* | 99.97% | 149409 | 537 |
| Lat_Neo | *Homo sapiens* | 99.97% | 139658 | 398 |
| Mota | *Homo sapiens* | 99.96% | 3329825 | 9431 |
| Iri_Bal | *Homo sapiens* | 84.71% | 3689879 | 14887 |
| Iri_Rat_1 | *Homo sapiens* | 75.49% | 3426319 | 12113 |
| Iri_Rat_2 | *Homo sapiens* | 48.57% | 363194 | 1184 |
| Iri_Rat_3 | *Homo sapiens* | 73.89% | 90332 | 335 |
| Bat_Afa | *Homo sapiens* | 99.99% | 915640 | 5112 |
| Kyt_And_1 | *Homo sapiens* | 100.00% | 293571 | 2138 |
| Kyt_And_2 | *Homo sapiens* | 99.99% | 1319757 | 6400 |
| Bed_Nor | *Homo sapiens* | 99.95% | 1572065 | 8468 |
| Qxie_Iro | *Homo sapiens* | 99.99% | 740380 | 6010 |
| Kyt_Iro | *Homo sapiens* | 99.99% | 288039 | 1905 |
| Ver_Iro | *Homo sapiens* | 99.99% | 53223 | 186 |
| Sar_Iro | *Homo sapiens* | 99.99% | 68876 | 280 |
| Sab_Kar | *Homo sapiens* | 99.99% | 1849147 | 10615 |
| Arb_Kar_1 | *Homo sapiens* | 99.99% | 1181968 | 5506 |
| Arb_Kar_2 | *Homo sapiens* | 100.00% | 621162 | 3904 |
| Arb_Kar_3 | *Homo sapiens* | 99.99% | 2078086 | 9599 |
| Bys_Kar_1 | *Homo sapiens* | 100.00% | 215463 | 1474 |
| Bys_Kar_2 | *Homo sapiens* | 99.99% | 221642 | 1414 |
| Kap_Mez | *Homo sapiens* | 99.99% | 587035 | 3321 |
| Bol_Sin | *Homo sapiens* | 99.88% | 618838 | 4956 |
| Prz_Une | *Homo sapiens* | 99.96% | 768674 | 6404 |
| Erd_Vat | *Homo sapiens* | 99.95% | 243256 | 1248 |
| Ulan_Yam | *Homo sapiens* | 99.99% | 649795 | 6021 |
| Eng_Rom_1 | *Homo sapiens* | 32.21% | 345161 | 1481 |
| Eng_Rom_2 | *Homo sapiens* | 53.33% | 195472 | 713 |
| Eng_Rom_3 | *Homo sapiens* | 67.58% | 264638 | 960 |
| Eng_Rom_4 | *Homo sapiens* | 51.37% | 202184 | 731 |
| Eng_Rom_5 | *Homo sapiens* | 76.51% | 438985 | 1498 |
| Eng_Ang | *Homo sapiens* | 61.61% | 181117 | 591 |
| Present-day_human_1 | *Homo sapiens* | 98.44% | 4648419 | 13247 |
| Present-day_human_2 | *Homo sapiens* | 98.72% | 5323034 | 13569 |

**Supplementary Table 3.**

Cancer genes (n=401) used in this study.

| **Categories** | **Genes** |
| --- | --- |
| Oncogenes  (n=78) | *AKT1; ALK; AR; ARL15; ATXN2; BCLAF1; BRAF; BTRC; CCND1; CDK4; CHD4; CLEC4C; CTNNB1; DICER1; DIRC2; DIS3; DKK2; DNAJB5; EGFR; EIF1AX; EIF2S2; ERBB2; ERBB3; EZH2; FBN2; FGFR2; FGFR3; FLT3; GK2; GOT2; GPX7; HNRNPA1; HRAS; IDH1; KAT8; KCNMB4; KLF4; KRAS; KRT15; LAMTOR1; MAP2K1; MAP2K2; MAPK1; MAX; MB21D2; MED12; MTOR; MYC; MYD88; NFE2L2; NKX3-1; NRAS; PCBP1; PIK3CA; PIK3CB; PPARD; PPP2R1A; PPP6C; PRKCI; PTPN11; RAC1; RAF1; RHEB; RHOA; RIT1; RRAS2; SF3B1; SKAP2; SMAD3; SNCB; SOS1; SPOP; TARDBP; TTR; TUBA3C; U2AF1; XPO1; ZNF600* |
| Tumor suppressor genes  (n=211) | *ACVR1B; ACVR2A; AJUBA; ALB; ALDOB; AMBRA1; AMER1; AMOT; ANKRD46; APC; ARHGAP35; ARID1A; ARID1B; ARID2; ARID4B; ASXL1; ASXL2; ATAD2; ATG14; ATG5; ATM; ATRX; AXIN1; B2M; BAP1; BCL9; BCOR; BRCA1; BRCA2; BRD7; BRE; BTBD7; C2CD5; C8orf34; CASP8; CBFB; CCAR1; CCDC88A; CD1D; CDH1; CDK12; CDKN1A; CDKN1B; CDKN2A; CEBPA; CHD2; CHD8; CHRDL1; CIC; COPS4; CREBBP; CSDE1; CTCF; CTDNEP1; CUL2; DACH1; DDX5; DHX15; DNER; DNM1L; DNMT3A; EED; EIF2AK3; ELF3; EMG1; EP300; EPB41L4A; EPHA2; ERRFI1; EXO5; FANCM; FAT1; FBXW7; FOXA1; FUBP1; GATA3; GGCT; GIGYF2; GNPTAB; GPS2; GSE1; HERC1; HERC4; HLA-A; HLA-B; HSP90AB1; ID3; IFNGR2; INPPL1; IWS1; JAK1; JAK2; KANSL1; KBTBD7; KDM5C; KDM6A; KEAP1; KMT2A; KMT2B; KMT2C; KMT2D; KMT2E; LARP4B; MAP2K4; MAP3K1; MAP4K3; MBD1; MBD6; MBNL1; MED23; MEN1; MGA; MLLT4; MOAP1; MORC4; MYO1B; MYO6; NAA15; NAA25; NCOA2; NCOR1; NEK9; NF1; NF2; NFE2L3; NIPBL; NIT1; NOTCH1; NOTCH2; NPM1; NSD1; PBRM1; PHF6; PIK3R1; POLA2; PPM1D; PRPF40A; PSIP1; PTEN; RANBP3L; RAPGEF6; RASA1; RB1; RBBP6; RBM10; RBM26; RC3H2; RERE; RNF111; RNF43; RPL11; RPL5; RUNX1; SCAF11; SEC22A; SENP3; SENP8; SETD2; SFPQ; SIN3A; SMAD2; SMAD4; SMARCA4; SMARCB1; SOX9; SP3; SPEN; SPSB2; STAG2; STK11; SUFU; TBL1XR1; TBX3; TCF12; TCF7L2; TET2; TEX11; TFDP2; TGFBR2; THRAP3; TMCO2; TMED10; TMEM30A; TMPO; TNRC6B; TP53; TP53BP1; TRAF3; TRIP12; TTK; UBE2D3; UNC13C; USP28; USP9X; VHL; VPS33B; WAC; WDR33; WDR47; WT1; WWP1; ZC3H13; ZFHX3; ZFP36L1; ZFP36L2; ZMYM3; ZMYM4; ZNF234; ZNF292; ZNF318; ZNF750* |
| Double role genes  (n=20) | *BIRC6; BRWD3; CDC73; CEP128; CNOT1; COL11A1; CSMD3; CUL1; DCHS1; DDX3X; FN1; HGF; HMCN1; IDH2; INPP4A; JUN; MYCN; PTN; RIMS2; UBR5* |
| Unclassified genes  (n=92) | *ABCA7; ARMCX1; BIRC8; BLVRA; C11orf70; C12orf57; C3orf62; CAMKV; CAPG; CBX4; CCDC117; CCM2; CCNC; CCR3; CD79B; CDCP1; CELF1; CENPB; CHEK2; CHUK; CMTR2; CNN2; CNOT4; COX7B2; CYB5B; DCUN1D1; DHX16; DIXDC1; EIF4A1; EIF4A2; ADGRE3; EPS8; F5; FCER1G; FGFR1; FUS; GALNTL5; GLIPR2; GNRHR; GOLM1; GRK1; GZMA; HDAC1; HIST1H2BO; IFT88; IKZF2; INO80C; KATNAL1; FAM234B; LPAR2; LYN; MBNL3; MKLN1; MS4A1; MSI1; MYL6; NAP1L2; NAP1L4; NME4; PCOLCE2; POT1; PRKACA; PTH2; PTMS; RAB18; REL; RFC4; RQCD1; RXRA; SARM1; SETD1B; SF3A3; SMARCC2; SOX4; STK31; TAF1A; TAS2R30; TM9SF1; TMEM107; TNFRSF9; TRIM8; TSC1; UNKL; UPP1; USO1; VN1R2; YOD1; ZDHHC4; ZGRF1; ZNF268; ZNF345; ZNF800* |

**Supplementary Table 4.**

Exon sequence similarity for cancer genes using non-silent SNPs of 21 non-human primate genomes and 54 ancient human genomes.

**Supplementary Table 5.**

Exon sequence similarity for cancer genes and whole genome genes using protein sequences derived from 12 assembled non-human primate genomes. panTro4: *Pan troglodytes*; panPan1: *Pan paniscus*; gorGor3: *Gorilla gorilla*; ponAbe2: *Pongo pygmaeus*; nomLeu3: *Nomascus leucogenys*; macFas5: *Macaca fascicularis*; rhiRox1: *Rhinopithecus roxellana*; calJac3: *Callithrix jacchus*; saiBol1: *Saimiri boliviensis*; tarSyr2: *Tarsius syrichta*; otoGar3: *Otolemur garnettii*; micMur1: *Microcebus marinus*.

**Supplementary Table 6.**

dN/dS of human reference genome when compared with other 7 primate species. These 7 species are: chimpanzee (*Pan troglodytes*), orangutan (*Pongo abelii*), gibbon (*Nomascus leucogenys*), macaque (*Macaca mulatta*), marmoset (*Callithrix jacchus*), bushbaby (*Otolemur garnettii*), mouse lemur (*Microcebus murinus*).

**Supplementary Table 7.**

Result of McDonald-Kreitman test. Px: the number of non-synonymous polymorphisms; Ps: the number of synonymous polymorphisms; Dx: the number of non-synonymous substitutions; Ds: the number of synonymous substitutions; AS Index: adaptive selection index.

**Supplementary Table 8.**

Potential functional consequences of 299 human-specific fixed SNPs.

| **Chromosome** | **Site (GRCh38)** | **Human base** | **Inferred ancestral base** | **Gene name** | **Human residue** | **Codon** | **Inferred ancestral residue** | **Change of hydrophobicity?** | **Change of charge?** | **Change of volume?** | **Located at interfaces?** | **Close to "Pathogenic" mutations of ClinVar?** | **CHASM p-value** | **CHASM score** |
| --- | --- | --- | --- | --- | --- | --- | --- | --- | --- | --- | --- | --- | --- | --- |
| 1 | 15930285 | G | A | *SPEN* | G | 1349 | S | Yes |  |  |  |  | 0.1804 | 0.466 |
| 1 | 32331784 | C | A | *HDAC1* | D | 399 | E |  |  |  |  |  | 0.2358 | 0.43 |
| 1 | 35399483 | C | A | *ZMYM4* | D | 1145 | E |  |  |  |  |  | 0.355 | 0.366 |
| 1 | 64866937 | C | T | *JAK1* | G | 307 | S | Yes |  |  |  |  | 0.266 | 0.412 |
| 1 | 102879703 | T | C | *COL11A1* | T | 1764 | A | Yes |  |  |  |  | 0.2102 | 0.446 |
| 1 | 119921769 | T | A | *NOTCH2* | T | 1752 | S |  |  |  |  |  | 0.342 | 0.372 |
| 1 | 119941728 | T | C | *NOTCH2* | M | 927 | V |  |  |  |  |  | 0.0028 | 0.806 |
| 1 | 147623970 | G | A | *BCL9* | V | 1098 | M |  |  |  |  |  | 0.0446 | 0.634 |
| 1 | 161120209 | A | G | *NIT1* | I | 232 | V |  |  |  |  |  | 0.1804 | 0.466 |
| 1 | 169515495 | C | G | *F5* | E | 2164 | D |  |  |  |  |  | 0.4988 | 0.292 |
| 1 | 169528058 | T | C | *F5* | K | 1824 | R |  |  |  |  |  | 0.427 | 0.326 |
| 1 | 169541533 | A | G | *F5* | I | 1191 | T | Yes |  | Yes |  |  | 0.362 | 0.362 |
| 1 | 169542280 | C | T | *F5* | S | 942 | N |  |  |  |  |  | 0.4354 | 0.322 |
| 1 | 185923512 | C | G | *HMCN1* | H | 382 | D |  | Yes |  |  |  | 0.5332 | 0.278 |
| 1 | 185928575 | C | A | *HMCN1* | A | 487 | E | Yes | Yes |  |  |  | 0.598 | 0.25 |
| 1 | 185997470 | A | C | *HMCN1* | T | 1274 | P | Yes |  |  |  |  | 0.3662 | 0.36 |
| 1 | 185997504 | A | C | *HMCN1* | E | 1285 | A | Yes | Yes |  |  |  | 0.3204 | 0.382 |
| 1 | 186001619 | C | T | *HMCN1* | T | 1409 | I | Yes |  | Yes |  |  | 0.2064 | 0.448 |
| 1 | 186001652 | G | A | *HMCN1* | R | 1420 | K |  |  |  |  |  | 0.336 | 0.374 |
| 1 | 186018277 | A | G | *HMCN1* | N | 1799 | D |  | Yes |  |  |  | 0.1952 | 0.456 |
| 1 | 186045849 | A | G | *HMCN1* | R | 2156 | G | Yes | Yes | Yes |  |  | 0.1498 | 0.488 |
| 1 | 186055522 | A | G | *HMCN1* | H | 2331 | R |  |  |  |  |  | 0.1308 | 0.506 |
| 1 | 186065396 | C | G | *HMCN1* | H | 2558 | D |  | Yes |  |  |  | 0.383 | 0.35 |
| 1 | 186108494 | G | A | *HMCN1* | R | 3629 | Q |  | Yes |  |  |  | 0.3322 | 0.376 |
| 1 | 186144334 | A | C | *HMCN1* | N | 4696 | H |  | Yes |  |  |  | 0.3944 | 0.344 |
| 1 | 186144614 | G | A | *HMCN1* | G | 4726 | D | Yes | Yes |  |  |  | 0.2284 | 0.434 |
| 1 | 186152800 | T | G | *HMCN1* | S | 4983 | A | Yes |  |  |  |  | 0.4518 | 0.314 |
| 1 | 222579797 | T | C | *TAF1A* | N | 123 | D |  | Yes |  |  |  | 0.2762 | 0.406 |
| 1 | 235219888 | G | T | *ARID4B* | D | 496 | E |  |  |  |  |  | 0.2236 | 0.438 |
| 1 | 235220360 | C | T | *ARID4B* | R | 450 | K |  |  |  |  |  | 0.4388 | 0.32 |
| 2 | 25743172 | G | T | *ASXL2* | H | 1055 | Q |  | Yes |  |  |  | 0.4138 | 0.334 |
| 2 | 25743353 | G | C | *ASXL2* | A | 995 | G |  |  |  |  |  | 0.5088 | 0.288 |
| 2 | 25750277 | A | G | *ASXL2* | S | 427 | P | Yes |  |  |  |  | 0.087 | 0.562 |
| 2 | 29694925 | T | C | *ALK* | I | 293 | V |  |  |  |  |  | 0.0962 | 0.548 |
| 2 | 32443528 | C | T | *BIRC6* | P | 1426 | S | Yes |  |  |  |  | 0.051 | 0.622 |
| 2 | 32515367 | G | A | *BIRC6* | S | 3649 | N |  |  |  |  |  | 0.1056 | 0.536 |
| 2 | 32545822 | T | G | *BIRC6* | S | 4258 | A | Yes |  |  |  |  | 0.266 | 0.412 |
| 2 | 39299765 | C | T | *MAP4K3* | G | 386 | S | Yes |  |  |  |  | 0.263 | 0.414 |
| 2 | 39315312 | G | T | *MAP4K3* | T | 332 | N |  |  |  |  |  | 0.3044 | 0.39 |
| 2 | 39356243 | G | A | *MAP4K3* | T | 84 | I | Yes |  | Yes |  |  | 0.9034 | 0.084 |
| 2 | 55296263 | C | T | *CCDC88A* | V | 1696 | I |  |  |  |  |  | 0.2008 | 0.452 |
| 2 | 60921975 | A | G | *REL* | R | 434 | G | Yes | Yes | Yes |  |  | 0.4588 | 0.31 |
| 2 | 88574746 | C | T | *EIF2AK3* | V | 913 | M |  |  |  |  |  | 0.198 | 0.454 |
| 2 | 152676613 | C | T | *PRPF40A* | A | 275 | T | Yes |  |  |  |  | 0.9426 | 0.056 |
| 2 | 177233221 | T | G | *NFE2L2* | E | 128 | A | Yes | Yes |  |  |  | 0.8884 | 0.098 |
| 2 | 208248584 | G | A | *IDH1* | H | 67 | Y |  | Yes |  |  |  | 0.1804 | 0.466 |
| 2 | 213007674 | G | T | *IKZF2* | H | 429 | N |  | Yes |  |  |  | 0.2262 | 0.436 |
| 2 | 215386805 | T | G | *FN1* | H | 1499 | P | Yes | Yes |  |  |  | 0.2358 | 0.43 |
| 2 | 215394669 | T | A | *FN1* | N | 1219 | Y |  |  | Yes |  |  | 0.294 | 0.396 |
| 2 | 229447438 | T | C | *DNER* | N | 455 | S |  |  |  |  |  | 0.1042 | 0.538 |
| 3 | 46266116 | T | G | *CCR3* | L | 341 | V |  |  |  |  |  | 0.5474 | 0.272 |
| 3 | 47120576 | C | T | *SETD2* | D | 1354 | N |  | Yes |  |  |  | 0.4632 | 0.308 |
| 3 | 47122349 | T | A | *SETD2* | M | 763 | L |  |  |  |  |  | 0.3944 | 0.344 |
| 3 | 47123540 | G | A | *SETD2* | P | 366 | S | Yes |  |  |  |  | 0.1834 | 0.464 |
| 3 | 47123848 | A | G | *SETD2* | L | 263 | S | Yes |  | Yes |  |  | 0.402 | 0.34 |
| 3 | 49271377 | T | C | *C3orf62* | M | 203 | V |  |  |  |  |  | 0.374 | 0.356 |
| 3 | 52408019 | C | T | *BAP1* | S | 105 | N |  |  |  |  | Yes | 0.2998 | 0.392 |
| 3 | 52668509 | A | C | *PBRM1* | S | 125 | A | Yes |  |  |  |  | 0.355 | 0.366 |
| 3 | 142829760 | A | T | *PCOLCE2* | I | 266 | K | Yes | Yes |  |  |  | 0.355 | 0.366 |
| 3 | 186804640 | C | G | *RFC4* | S | 25 | T |  |  |  |  |  | 0.6692 | 0.218 |
| 4 | 67740569 | A | C | *GNRHR* | L | 300 | V |  |  |  |  | Yes | 0.4304 | 0.324 |
| 4 | 67753873 | C | G | *GNRHR* | V | 155 | L |  |  |  |  | Yes | 0.634 | 0.234 |
| 4 | 73409419 | A | G | *ALB* | K | 183 | E |  | Yes |  | ALB-ALB |  | 0.6122 | 0.244 |
| 4 | 73413516 | A | C | *ALB* | I | 314 | L |  |  |  |  |  | 0.6028 | 0.248 |
| 4 | 79406640 | C | T | *GK2* | G | 521 | S | Yes |  |  |  |  | 0.4988 | 0.292 |
| 4 | 83073311 | C | T | *COPS4* | S | 398 | F | Yes |  | Yes |  |  | 0.8884 | 0.098 |
| 4 | 105234107 | C | G | *TET2* | H | 76 | Q |  | Yes |  |  |  | 0.263 | 0.414 |
| 4 | 105234820 | G | C | *TET2* | S | 314 | T |  |  |  |  |  | 0.3898 | 0.346 |
| 4 | 105234862 | T | C | *TET2* | L | 328 | P |  |  | Yes |  |  | 0.087 | 0.562 |
| 4 | 105236146 | G | A | *TET2* | S | 756 | N |  |  |  |  |  | 0.0286 | 0.676 |
| 4 | 105236779 | C | T | *TET2* | T | 967 | I | Yes |  | Yes |  |  | 0.0962 | 0.548 |
| 4 | 105237050 | C | G | *TET2* | H | 1057 | Q |  | Yes |  |  |  | 0.3692 | 0.358 |
| 4 | 106925817 | T | C | *DKK2* | S | 119 | G | Yes |  |  |  |  | 0.2428 | 0.426 |
| 4 | 112587395 | G | A | *ZGRF1* | S | 1221 | L | Yes |  | Yes |  |  | 0.5532 | 0.27 |
| 4 | 112587665 | T | G | *ZGRF1* | N | 1131 | T |  |  |  |  |  | 0.3586 | 0.364 |
| 4 | 112617544 | C | G | *ZGRF1* | C | 833 | S | Yes |  |  |  |  | 0.336 | 0.374 |
| 4 | 112618909 | T | C | *ZGRF1* | Y | 378 | C | Yes |  |  |  |  | 0.2542 | 0.418 |
| 4 | 112619578 | C | T | *ZGRF1* | C | 155 | Y | Yes |  | Yes |  |  | 0.1114 | 0.53 |
| 4 | 152382317 | G | C | *FBXW7* | P | 7 | A |  |  |  |  |  | 0.676 | 0.216 |
| 4 | 186588816 | G | A | *FAT1* | P | 4517 | S | Yes |  |  |  |  | 0.2284 | 0.434 |
| 4 | 186611590 | T | C | *FAT1* | I | 3219 | V |  |  |  |  |  | 0.2762 | 0.406 |
| 4 | 186617168 | G | T | *FAT1* | T | 2973 | N |  |  |  |  |  | 0.2912 | 0.398 |
| 4 | 186618523 | A | G | *FAT1* | L | 2690 | P |  |  | Yes |  |  | 0.2262 | 0.436 |
| 4 | 186618740 | C | T | *FAT1* | V | 2618 | I |  |  |  |  |  | 0.4816 | 0.3 |
| 4 | 186620005 | C | T | *FAT1* | S | 2196 | N |  |  |  |  |  | 0.2262 | 0.436 |
| 4 | 186636150 | A | G | *FAT1* | L | 1353 | P |  |  | Yes |  |  | 0.1156 | 0.524 |
| 4 | 186707153 | G | T | *FAT1* | S | 892 | Y |  |  | Yes |  |  | 0.676 | 0.216 |
| 4 | 186707194 | G | T | *FAT1* | N | 878 | K |  | Yes | Yes |  |  | 0.6986 | 0.206 |
| 4 | 186707945 | C | T | *FAT1* | R | 628 | Q |  | Yes |  |  |  | 0.4518 | 0.314 |
| 4 | 186708647 | G | C | *FAT1* | A | 394 | G |  |  |  |  |  | 0.7454 | 0.184 |
| 5 | 36261964 | C | G | *RANBP3L* | D | 212 | H |  | Yes |  |  |  | 0.3864 | 0.348 |
| 5 | 36301379 | G | A | *RANBP3L* | P | 13 | L |  |  | Yes |  |  | 0.3864 | 0.348 |
| 5 | 112209922 | A | G | *EPB41L4A* | I | 383 | T | Yes |  | Yes |  |  | 0.2452 | 0.424 |
| 5 | 112839106 | G | A | *APC* | R | 1171 | H |  |  |  |  |  | 0.087 | 0.562 |
| 5 | 112839321 | C | A | *APC* | P | 1243 | T | Yes |  |  |  |  | 0.216 | 0.442 |
| 5 | 112839582 | C | A | *APC* | P | 1330 | T | Yes |  |  |  |  | 0.2202 | 0.44 |
| 5 | 112841812 | G | C | *APC* | G | 2073 | A |  |  |  |  |  | 0.2262 | 0.436 |
| 5 | 112843452 | T | A | *APC* | F | 2620 | I |  |  |  |  |  | 0.402 | 0.34 |
| 5 | 112843663 | C | G | *APC* | A | 2690 | G |  |  |  |  |  | 0.3692 | 0.358 |
| 5 | 128276121 | T | A | *FBN2* | Y | 2504 | F | Yes |  |  |  |  | 0.1648 | 0.476 |
| 5 | 128311316 | A | T | *FBN2* | D | 1686 | E |  |  |  |  |  | 0.3788 | 0.352 |
| 5 | 177210668 | C | A | *NSD1* | L | 757 | M |  |  |  |  |  | 0.3692 | 0.358 |
| 5 | 177210801 | C | T | *NSD1* | A | 801 | V |  |  |  |  |  | 0.0602 | 0.602 |
| 5 | 177211394 | C | G | *NSD1* | L | 999 | V |  |  |  |  |  | 0.2762 | 0.406 |
| 5 | 177211620 | G | A | *NSD1* | R | 1074 | H |  |  |  |  |  | 0.3586 | 0.364 |
| 5 | 177212088 | C | T | *NSD1* | A | 1230 | V |  |  |  |  |  | 0.0812 | 0.572 |
| 6 | 43337820 | C | T | *ZNF318* | A | 2060 | T | Yes |  |  |  |  | 0.0842 | 0.566 |
| 6 | 43337885 | A | G | *ZNF318* | V | 2038 | A |  |  |  |  |  | 0.1518 | 0.486 |
| 6 | 43338069 | T | C | *ZNF318* | T | 1977 | A | Yes |  |  |  |  | 0.1706 | 0.472 |
| 6 | 43339010 | C | T | *ZNF318* | G | 1663 | D | Yes | Yes |  |  |  | 0.1444 | 0.494 |
| 6 | 43340097 | G | C | *ZNF318* | L | 1301 | V |  |  |  |  |  | 0.2396 | 0.428 |
| 6 | 43340130 | A | C | *ZNF318* | L | 1290 | V |  |  |  |  |  | 0.2596 | 0.416 |
| 6 | 43355683 | C | G | *ZNF318* | V | 551 | L |  |  |  |  |  | 0.5138 | 0.286 |
| 6 | 43365422 | A | G | *ZNF318* | C | 140 | R | Yes | Yes | Yes |  |  | 0.0762 | 0.58 |
| 6 | 75886953 | A | G | *MYO6* | N | 873 | D |  | Yes |  |  |  | 0.2884 | 0.4 |
| 6 | 75886974 | A | G | *MYO6* | T | 880 | A | Yes |  |  |  |  | 0.216 | 0.442 |
| 6 | 75907619 | T | A | *MYO6* | L | 1074 | Q | Yes |  |  |  |  | 0.3044 | 0.39 |
| 6 | 75914102 | G | A | *MYO6* | R | 1170 | Q |  | Yes |  |  |  | 0.1498 | 0.488 |
| 6 | 80027978 | T | G | *TTK* | H | 496 | Q |  | Yes |  |  |  | 0.2802 | 0.404 |
| 6 | 80036582 | G | A | *TTK* | V | 678 | I |  |  |  |  |  | 0.0432 | 0.638 |
| 6 | 80040606 | T | G | *TTK* | V | 798 | G |  |  |  |  |  | 0.4102 | 0.336 |
| 6 | 87233511 | G | A | *ZNF292* | G | 242 | E | Yes | Yes |  |  |  | 0.4304 | 0.324 |
| 6 | 156829396 | C | G | *ARID1B* | A | 571 | G |  |  |  |  |  | 0.3864 | 0.348 |
| 6 | 157174013 | G | A | *ARID1B* | V | 1011 | M |  |  |  |  |  | 0.318 | 0.384 |
| 7 | 6585111 | G | C | *ZDHHC4* | V | 198 | L |  |  |  |  |  | 0.3944 | 0.344 |
| 7 | 23769110 | G | A | *STK31* | S | 511 | N |  |  |  |  |  | 0.3458 | 0.37 |
| 7 | 23832238 | C | A | *STK31* | Q | 978 | K |  | Yes |  |  |  | 0.1308 | 0.506 |
| 7 | 26185212 | C | G | *NFE2L3* | T | 505 | S |  |  |  |  |  | 0.7706 | 0.174 |
| 7 | 26725475 | A | G | *SKAP2* | L | 250 | P |  |  | Yes |  |  | 0.3692 | 0.358 |
| 7 | 105112060 | T | C | *KMT2E* | V | 1435 | A |  |  |  |  |  | 0.0924 | 0.554 |
| 7 | 124825297 | C | T | *POT1* | V | 583 | M |  |  |  |  |  | 0.336 | 0.374 |
| 7 | 124827305 | G | A | *POT1* | A | 532 | V |  |  |  |  |  | 0.1444 | 0.494 |
| 7 | 131470861 | G | A | *MKLN1* | V | 650 | M |  |  |  |  |  | 0.5182 | 0.284 |
| 7 | 151983057 | G | T | *GALNTL5* | C | 147 | F |  |  | Yes |  |  | 0.402 | 0.34 |
| 7 | 151983072 | A | G | *GALNTL5* | Q | 152 | R |  | Yes |  |  |  | 0.0694 | 0.588 |
| 7 | 152002947 | T | C | *GALNTL5* | S | 298 | P | Yes |  |  |  |  | 0.3898 | 0.346 |
| 7 | 152007918 | A | G | *GALNTL5* | R | 334 | G | Yes | Yes | Yes |  |  | 0.4208 | 0.33 |
| 7 | 152138821 | C | T | *KMT2C* | S | 4873 | N |  |  |  |  |  | 0.087 | 0.562 |
| 7 | 152152881 | C | T | *KMT2C* | S | 4117 | N |  |  |  |  |  | 0.1952 | 0.456 |
| 7 | 152176476 | G | A | *KMT2C* | L | 2993 | F |  |  |  |  |  | 0.0952 | 0.55 |
| 7 | 152176551 | C | T | *KMT2C* | A | 2968 | T | Yes |  |  |  |  | 0.0372 | 0.654 |
| 7 | 152176582 | G | T | *KMT2C* | F | 2957 | L |  |  |  |  |  | 0.0992 | 0.544 |
| 7 | 152176649 | G | A | *KMT2C* | S | 2935 | L | Yes |  | Yes |  |  | 0.0824 | 0.57 |
| 7 | 152181547 | C | T | *KMT2C* | V | 2105 | M |  |  |  |  |  | 0.0432 | 0.638 |
| 7 | 152181865 | T | C | *KMT2C* | I | 1999 | V |  |  |  |  |  | 0.1462 | 0.492 |
| 7 | 152187806 | A | G | *KMT2C* | S | 1568 | P | Yes |  |  |  |  | 0.1416 | 0.496 |
| 7 | 152187844 | G | A | *KMT2C* | A | 1555 | V |  |  |  |  |  | 0.163 | 0.478 |
| 7 | 152250903 | G | A | *KMT2C* | A | 562 | V |  |  |  |  |  | 0.114 | 0.526 |
| 8 | 23681609 | G | C | *NKX3-1* | S | 106 | C | Yes |  |  |  |  | 0.2542 | 0.418 |
| 8 | 68625620 | C | T | *C8orf34* | P | 15 | L |  |  | Yes |  |  | 0.7976 | 0.16 |
| 8 | 103766476 | C | G | *RIMS2* | H | 213 | D |  | Yes |  |  |  | 0.9446 | 0.054 |
| 8 | 103886127 | A | G | *RIMS2* | I | 288 | V |  |  |  |  |  | 0.1358 | 0.502 |
| 8 | 112255334 | A | G | *CSMD3* | F | 3319 | S | Yes |  | Yes |  |  | 0.4304 | 0.324 |
| 8 | 112304885 | A | G | *CSMD3* | I | 2701 | T | Yes |  | Yes |  |  | 0.1576 | 0.482 |
| 8 | 112573540 | T | A | *CSMD3* | T | 1335 | S |  |  |  |  |  | 0.3788 | 0.352 |
| 8 | 112682459 | A | G | *CSMD3* | L | 887 | P |  |  | Yes |  |  | 0.1518 | 0.486 |
| 8 | 113436793 | T | G | *CSMD3* | K | 21 | T |  | Yes | Yes |  |  | 0.2064 | 0.448 |
| 8 | 123322961 | G | A | *ATAD2* | H | 1370 | Y |  | Yes |  |  |  | 0.294 | 0.396 |
| 8 | 123325963 | G | C | *ATAD2* | T | 1311 | S |  |  |  |  |  | 0.537 | 0.276 |
| 8 | 123346115 | T | C | *ATAD2* | T | 835 | A | Yes |  |  |  |  | 0.4914 | 0.296 |
| 9 | 122851171 | A | G | *RC3H2* | M | 1097 | T | Yes |  | Yes |  |  | 0.1378 | 0.5 |
| 9 | 122865398 | C | T | *RC3H2* | A | 529 | T | Yes |  |  |  |  | 0.0694 | 0.588 |
| 10 | 825822 | C | T | *LARP4B* | A | 392 | T | Yes |  |  |  |  | 0.093 | 0.552 |
| 10 | 28535632 | G | C | *WAC* | G | 50 | A |  |  |  |  |  | 0.4934 | 0.294 |
| 10 | 28608206 | C | T | *WAC* | P | 314 | S | Yes |  |  |  |  | 0.1042 | 0.538 |
| 10 | 67941041 | C | A | *HERC4* | C | 809 | F |  |  | Yes | HERC4-HERC4 |  | 0.2998 | 0.392 |
| 10 | 67992290 | T | C | *HERC4* | N | 394 | D |  | Yes |  |  |  | 0.0962 | 0.548 |
| 10 | 68789853 | A | C | *CCAR1* | M | 1111 | L |  |  |  |  |  | 0.2912 | 0.398 |
| 11 | 6633538 | C | T | *DCHS1* | D | 777 | N |  | Yes |  |  |  | 0.336 | 0.374 |
| 11 | 60462525 | A | G | *MS4A1* | T | 51 | A | Yes |  |  |  |  | 0.4102 | 0.336 |
| 11 | 102058950 | T | C | *C11orf70* | I | 88 | T | Yes |  | Yes |  |  | 0.1308 | 0.506 |
| 11 | 108267179 | C | A | *ATM* | F | 825 | L |  |  |  |  |  | 0.1086 | 0.532 |
| 11 | 108268582 | A | C | *ATM* | E | 937 | D |  |  |  |  |  | 0.2284 | 0.434 |
| 11 | 108281019 | A | C | *ATM* | T | 1143 | P | Yes |  |  |  |  | 0.3204 | 0.382 |
| 11 | 108284390 | A | G | *ATM* | R | 1304 | G | Yes | Yes | Yes |  |  | 0.2596 | 0.416 |
| 11 | 108293370 | A | C | *ATM* | I | 1557 | L |  |  |  |  |  | 0.3788 | 0.352 |
| 11 | 108321320 | A | T | *ATM* | M | 2158 | L |  |  |  |  |  | 0.2236 | 0.438 |
| 11 | 111968548 | G | A | *DIXDC1* | G | 76 | S | Yes |  |  |  |  | 0.3898 | 0.346 |
| 11 | 111980825 | G | T | *DIXDC1* | A | 249 | S | Yes |  |  |  |  | 0.6444 | 0.23 |
| 11 | 113813946 | G | T | *USP28* | T | 561 | N |  |  |  |  |  | 0.8292 | 0.14 |
| 11 | 118473450 | G | C | *KMT2A* | S | 764 | T |  |  |  |  |  | 0.2102 | 0.446 |
| 11 | 118502791 | T | C | *KMT2A* | V | 2300 | A |  |  |  |  |  | 0.073 | 0.584 |
| 11 | 118504815 | A | G | *KMT2A* | S | 2975 | G | Yes |  |  |  |  | 0.2762 | 0.406 |
| 12 | 6974418 | C | T | *EMG1* | A | 83 | V |  |  |  |  | Yes | 0.2802 | 0.404 |
| 12 | 22457159 | T | C | *C2CD5* | M | 900 | V |  |  |  |  |  | 0.0812 | 0.572 |
| 12 | 32701540 | G | A | *DNM1L* | G | 8 | E | Yes | Yes |  |  | Yes | 0.3982 | 0.342 |
| 12 | 45849627 | G | A | *ARID2* | S | 588 | N |  |  |  |  |  | 0.1678 | 0.474 |
| 12 | 45851673 | C | A | *ARID2* | P | 1184 | T | Yes |  |  |  |  | 0.1648 | 0.476 |
| 12 | 45852184 | G | A | *ARID2* | R | 1354 | K |  |  |  |  |  | 0.01 | 0.748 |
| 12 | 45927351 | T | C | *SCAF11* | T | 784 | A | Yes |  |  |  |  | 0.2064 | 0.448 |
| 12 | 45927936 | G | T | *SCAF11* | L | 589 | I |  |  |  |  |  | 0.35 | 0.368 |
| 12 | 45928289 | C | T | *SCAF11* | G | 471 | E | Yes | Yes |  |  |  | 0.1462 | 0.492 |
| 12 | 49028925 | C | T | *KMT2D* | G | 4762 | D | Yes | Yes |  |  |  | 0.3458 | 0.37 |
| 12 | 98534193 | G | T | *TMPO* | A | 646 | S | Yes |  |  |  |  | 0.634 | 0.234 |
| 12 | 98537591 | C | T | *TMPO* | P | 228 | S | Yes |  |  |  |  | 0.9728 | 0.032 |
| 12 | 101764338 | C | T | *GNPTAB* | S | 860 | N |  |  |  |  |  | 0.2452 | 0.424 |
| 12 | 101764465 | C | A | *GNPTAB* | V | 818 | L |  |  |  |  |  | 0.3044 | 0.39 |
| 12 | 101765206 | C | T | *GNPTAB* | V | 571 | I |  |  |  |  |  | 0.1648 | 0.476 |
| 12 | 112054557 | T | C | *NAA25* | T | 487 | A | Yes |  |  |  |  | 0.4988 | 0.292 |
| 12 | 112074696 | C | G | *NAA25* | S | 282 | T |  |  |  |  |  | 0.4428 | 0.318 |
| 12 | 114676342 | G | A | *TBX3* | A | 357 | V |  |  |  |  |  | 0.1042 | 0.538 |
| 12 | 133202525 | G | A | *ZNF268* | C | 280 | Y | Yes |  | Yes | ZNF268-DNA |  | 0.7454 | 0.184 |
| 12 | 133202560 | C | G | *ZNF268* | L | 292 | V |  |  |  | ZNF268-DNA |  | 0.6218 | 0.24 |
| 13 | 20601844 | A | G | *IFT88* | I | 327 | V |  |  |  |  |  | 0.2696 | 0.41 |
| 13 | 28004086 | A | G | *FLT3* | L | 983 | P |  |  | Yes |  |  | 0.2724 | 0.408 |
| 13 | 28004092 | A | G | *FLT3* | L | 981 | S | Yes |  | Yes |  |  | 0.2762 | 0.406 |
| 13 | 32331021 | G | A | *BRCA2* | A | 262 | T | Yes |  |  |  |  | 0.0406 | 0.644 |
| 13 | 32333187 | A | C | *BRCA2* | N | 570 | T |  |  |  |  |  | 0.0198 | 0.712 |
| 13 | 32337307 | A | C | *BRCA2* | E | 984 | D |  |  |  |  |  | 0.011 | 0.744 |
| 13 | 32338533 | C | T | *BRCA2* | A | 1393 | V |  |  |  |  |  | 0.0032 | 0.798 |
| 13 | 32339114 | G | A | *BRCA2* | A | 1587 | T | Yes |  |  |  |  | 0.0022 | 0.818 |
| 13 | 32340269 | G | A | *BRCA2* | A | 1972 | T | Yes |  |  |  |  | 0.0024 | 0.81 |
| 13 | 32340917 | A | G | *BRCA2* | K | 2188 | E |  | Yes |  |  |  | 0.0222 | 0.7 |
| 13 | 32341085 | A | G | *BRCA2* | K | 2244 | E |  | Yes |  |  |  | 0.0252 | 0.688 |
| 13 | 32363187 | C | T | *BRCA2* | T | 2662 | M | Yes |  | Yes | BRCA2-DSS1 | Yes | 0.0028 | 0.806 |
| 13 | 45967597 | G | C | *ZC3H13* | L | 1410 | V |  |  |  |  |  | 0.4428 | 0.318 |
| 14 | 45148912 | A | G | *FANCM* | T | 279 | A | Yes |  |  |  |  | 0.0924 | 0.554 |
| 14 | 45173055 | G | A | *FANCM* | A | 721 | T | Yes |  |  |  |  | 0.2852 | 0.402 |
| 14 | 45176067 | A | G | *FANCM* | S | 1105 | G | Yes |  |  |  |  | 0.7414 | 0.186 |
| 14 | 45176223 | G | A | *FANCM* | E | 1157 | K |  | Yes |  |  |  | 0.4174 | 0.332 |
| 14 | 45183810 | C | T | *FANCM* | P | 1475 | S | Yes |  |  |  |  | 0.1602 | 0.48 |
| 14 | 80497486 | C | T | *CEP128* | G | 1093 | E | Yes | Yes |  |  |  | 0.4768 | 0.302 |
| 14 | 80504986 | C | T | *CEP128* | R | 1036 | H |  |  |  |  |  | 0.4882 | 0.298 |
| 14 | 80530879 | T | C | *CEP128* | Q | 963 | R |  | Yes |  |  |  | 0.1066 | 0.534 |
| 14 | 80830243 | A | G | *CEP128* | L | 417 | S | Yes |  | Yes |  |  | 0.7976 | 0.16 |
| 14 | 93183881 | C | G | *MOAP1* | S | 121 | T |  |  |  |  |  | 0.5912 | 0.254 |
| 14 | 93242793 | A | G | *BTBD7* | L | 960 | P |  |  | Yes |  |  | 0.097 | 0.546 |
| 14 | 93294043 | G | T | *BTBD7* | T | 326 | K |  | Yes | Yes |  |  | 0.4934 | 0.294 |
| 15 | 41669688 | A | G | *MGA* | N | 265 | S |  |  |  |  |  | 0.5088 | 0.288 |
| 15 | 41711059 | G | A | *MGA* | V | 932 | M |  |  |  |  |  | 0.7746 | 0.172 |
| 15 | 41727374 | C | G | *MGA* | P | 1209 | A |  |  |  |  |  | 0.6398 | 0.232 |
| 15 | 41729309 | C | G | *MGA* | T | 1268 | S |  |  |  |  |  | 0.7078 | 0.2 |
| 15 | 41749626 | G | A | *MGA* | V | 2007 | I |  |  |  |  |  | 0.3944 | 0.344 |
| 15 | 41762238 | A | G | *MGA* | I | 2540 | M |  |  |  |  |  | 0.7746 | 0.172 |
| 15 | 41766286 | A | G | *MGA* | N | 2735 | S |  |  |  |  |  | 0.318 | 0.384 |
| 15 | 43446471 | C | T | *TP53BP1* | V | 986 | M |  |  |  |  |  | 0.3458 | 0.37 |
| 15 | 43456111 | G | C | *TP53BP1* | Q | 833 | E |  | Yes |  |  |  | 0.4518 | 0.314 |
| 15 | 43456472 | C | G | *TP53BP1* | M | 712 | I |  |  |  |  |  | 0.4174 | 0.332 |
| 15 | 43470051 | G | A | *TP53BP1* | T | 399 | M | Yes |  | Yes |  |  | 0.2696 | 0.41 |
| 15 | 54014195 | C | A | *UNC13C* | A | 431 | D | Yes | Yes |  |  |  | 0.4102 | 0.336 |
| 15 | 54014969 | A | G | *UNC13C* | D | 689 | G | Yes | Yes |  |  |  | 0.2396 | 0.428 |
| 15 | 54322051 | A | G | *UNC13C* | I | 1461 | V |  |  |  |  |  | 0.1576 | 0.482 |
| 15 | 57063754 | T | G | *TCF12* | I | 51 | M |  |  |  |  |  | 0.3458 | 0.37 |
| 15 | 59031053 | A | C | *RNF111* | E | 77 | D |  |  |  |  |  | 0.4816 | 0.3 |
| 15 | 63661787 | T | G | *HERC1* | M | 3046 | L |  |  |  |  |  | 0.2762 | 0.406 |
| 15 | 63674984 | C | T | *HERC1* | V | 2402 | I |  |  |  |  |  | 0.127 | 0.51 |
| 15 | 63774774 | C | T | *HERC1* | G | 284 | S | Yes |  |  |  |  | 0.4354 | 0.322 |
| 15 | 72139867 | A | C | *SENP8* | N | 82 | H |  | Yes |  |  |  | 0.686 | 0.212 |
| 16 | 24571895 | G | A | *RBBP6* | S | 1610 | N |  |  |  |  |  | 0.374 | 0.356 |
| 16 | 71283822 | C | T | *CMTR2* | R | 700 | Q |  | Yes |  |  |  | 0.5474 | 0.272 |
| 16 | 71284203 | A | T | *CMTR2* | V | 573 | E | Yes | Yes |  |  |  | 0.705 | 0.202 |
| 16 | 72796906 | C | T | *ZFHX3* | G | 1926 | S | Yes |  |  |  |  | 0.1028 | 0.54 |
| 17 | 31350311 | T | A | *NF1* | S | 2484 | T |  |  |  |  |  | 0.676 | 0.216 |
| 17 | 39501374 | A | T | *CDK12* | E | 848 | D |  |  |  |  |  | 0.0416 | 0.642 |
| 17 | 39530981 | C | G | *CDK12* | L | 1380 | V |  |  |  |  |  | 0.1858 | 0.462 |
| 17 | 43071053 | C | T | *BRCA1* | D | 1642 | N |  | Yes |  |  |  | 0.0222 | 0.702 |
| 17 | 43071148 | C | T | *BRCA1* | R | 1610 | H |  |  |  |  |  | 0.005 | 0.782 |
| 17 | 43082557 | G | C | *BRCA1* | H | 1402 | D |  | Yes |  |  |  | 0.0378 | 0.652 |
| 17 | 43092151 | T | C | *BRCA1* | Y | 1127 | C | Yes |  |  |  |  | 0.0022 | 0.814 |
| 17 | 43092350 | T | C | *BRCA1* | I | 1061 | V |  |  |  |  |  | 0.0072 | 0.772 |
| 17 | 43092595 | C | T | *BRCA1* | R | 979 | H |  |  |  |  |  | 0.0042 | 0.788 |
| 17 | 43092734 | C | A | *BRCA1* | G | 933 | C |  |  |  |  |  | 0.0028 | 0.806 |
| 17 | 43092863 | C | T | *BRCA1* | G | 890 | R | Yes | Yes |  |  |  | 0.0372 | 0.654 |
| 17 | 43092899 | C | G | *BRCA1* | A | 878 | P |  |  |  |  |  | 0.0144 | 0.728 |
| 17 | 43093066 | T | G | *BRCA1* | N | 822 | T |  |  |  |  |  | 0.0106 | 0.746 |
| 17 | 43093398 | C | G | *BRCA1* | K | 711 | N |  | Yes | Yes |  |  | 0.0022 | 0.814 |
| 17 | 43093519 | C | T | *BRCA1* | G | 671 | D | Yes | Yes |  |  |  | 0.0032 | 0.8 |
| 17 | 43094009 | G | C | *BRCA1* | P | 508 | A |  |  |  |  |  | 0.0154 | 0.724 |
| 17 | 43094131 | T | C | *BRCA1* | K | 467 | R |  |  |  |  |  | 0.0088 | 0.762 |
| 17 | 43094326 | T | C | *BRCA1* | E | 402 | G | Yes | Yes |  |  |  | 0.005 | 0.784 |
| 17 | 43094771 | C | T | *BRCA1* | A | 254 | T | Yes |  |  |  |  | 0.0092 | 0.754 |
| 17 | 43099795 | G | T | *BRCA1* | T | 176 | K |  | Yes | Yes |  |  | 0.0132 | 0.736 |
| 17 | 58357351 | T | C | *RNF43* | M | 682 | V |  |  |  |  |  | 0.8696 | 0.114 |
| 17 | 58360151 | G | A | *RNF43* | T | 317 | I | Yes |  | Yes |  |  | 0.2008 | 0.452 |
| 17 | 60663022 | G | C | *PPM1D* | V | 430 | L |  |  |  |  |  | 0.355 | 0.366 |
| 18 | 31598084 | A | T | *TTR* | N | 148 | Y |  |  | Yes |  |  | 0.8576 | 0.122 |
| 18 | 35454846 | C | T | *INO80C* | M | 190 | I |  |  |  |  |  | 0.5816 | 0.258 |
| 18 | 35454869 | T | A | *INO80C* | N | 183 | Y |  |  | Yes |  |  | 0.881 | 0.104 |
| 19 | 14655023 | T | G | *ADGRE3* | D | 179 | A | Yes | Yes |  |  |  | 0.676 | 0.216 |
| 19 | 36878188 | T | C | *ZNF345* | L | 453 | P |  |  | Yes |  |  | 0.376 | 0.354 |
| 20 | 32435446 | A | G | *ASXL1* | K | 912 | E |  | Yes |  |  |  | 0.2038 | 0.45 |
| 20 | 32435467 | A | G | *ASXL1* | I | 919 | V |  |  |  |  |  | 0.1156 | 0.524 |
| 20 | 32435698 | G | A | *ASXL1* | G | 996 | S | Yes |  |  |  |  | 0.0606 | 0.6 |
| 20 | 32435769 | A | T | *ASXL1* | R | 1019 | S |  | Yes | Yes |  |  | 0.2396 | 0.428 |
| 20 | 32436135 | C | G | *ASXL1* | D | 1141 | E |  |  |  |  |  | 0.3692 | 0.358 |
| 20 | 32436200 | A | G | *ASXL1* | D | 1163 | G | Yes | Yes |  |  |  | 0.2396 | 0.428 |
| 20 | 32436910 | G | C | *ASXL1* | E | 1400 | Q |  | Yes |  |  |  | 0.3898 | 0.346 |
| 21 | 33432229 | C | T | *IFNGR2* | S | 224 | F | Yes |  | Yes |  |  | 0.376 | 0.354 |
| 22 | 28781072 | A | G | *CCDC117* | I | 122 | V |  |  |  |  |  | 0.097 | 0.546 |
| 22 | 28786205 | T | C | *CCDC117* | V | 240 | A |  |  |  |  |  | 0.1254 | 0.512 |
| 22 | 28786247 | C | T | *CCDC117* | S | 254 | L | Yes |  | Yes |  |  | 0.3292 | 0.378 |
| 22 | 28786259 | C | A | *CCDC117* | P | 258 | Q | Yes |  |  |  |  | 0.2966 | 0.394 |
| X | 40071015 | T | C | *BCOR* | T | 1066 | A | Yes |  |  |  |  | 0.2102 | 0.446 |
| X | 41357837 | A | G | *DDX3X* | H | 657 | R |  |  |  |  |  | 0.598 | 0.25 |
| X | 77682596 | G | T | *ATRX* | T | 887 | N |  |  |  |  |  | 0.5816 | 0.258 |
| X | 77683803 | T | C | *ATRX* | T | 485 | A | Yes |  |  |  |  | 0.3692 | 0.358 |
| X | 80682030 | G | A | *BRWD3* | H | 1488 | Y |  | Yes |  |  |  | 0.1904 | 0.46 |
| X | 80688114 | A | C | *BRWD3* | I | 1273 | M |  |  |  |  |  | 0.266 | 0.412 |
| X | 132392310 | T | G | *MBNL3* | I | 123 | L |  |  |  |  |  | 0.3322 | 0.376 |

**Supplementary Table 9.**

"Pathogenic" mutations in ClinVar database. BRCA2, DNM1L, BAP1, EMG1, GNRHR were included in this table.

**Supplementary Table 10.**

Identity scoring for BRCA2. Human BRCA2 was used as reference.

**Supplementary Table 11.**

Root mean square deviation (RMSD) of molecular simulation for BRCA2-DSS1 complex.

**Supplementary Table 12.**

Data of bound DSS1 from four independent experiments.

| Experiment 1: | Primary | Dilution | Name | Area | M/T |
| --- | --- | --- | --- | --- | --- |
| 2662T-FLAG + DSS1-myc | Anti-DSS1 | 1:50 | DSS1 | 141624 |  |
| 2662M-FLAG + DSS1-myc | Anti-DSS1 | 1:50 | DSS1 | 172159 | 1.21560611 |
| 2662P-FLAG + DSS1-myc | Anti-DSS1 | 1:50 | DSS1 | 35440 |  |
| Experiment 2: | Primary | Dilution | Name | Area | M/T |
| 2662T-FLAG + DSS1-myc | Anti-DSS1 | 1:250 | DSS1 | 17113 |  |
| 2662M-FLAG + DSS1-myc | Anti-DSS1 | 1:250 | DSS1 | 23414 | 1.36819961 |
| 2662P-FLAG + DSS1-myc | Anti-DSS1 | 1:250 | DSS1 | 4248 |  |
| Experiment 3: | Primary | Dilution | Name | Area | M/T |
| 2662T-FLAG + DSS1-myc | Anti-DSS1 | 1:50 | DSS1 | 88908 |  |
| 2662M-FLAG + DSS1-myc | Anti-DSS1 | 1:50 | DSS1 | 96574 | 1.08622396 |
| 2662P-FLAG + DSS1-myc | Anti-DSS1 | 1:50 | DSS1 | 10240 |  |
| Experiment 4: | Primary | Dilution | Name | Area | M/T |
| 2662T-FLAG + DSS1-myc | Anti-DSS1 | 1:250 | DSS1 | 47591 |  |
| 2662M-FLAG + DSS1-myc | Anti-DSS1 | 1:250 | DSS1 | 58286 | 1.22472736 |
| 2662P-FLAG + DSS1-myc | Anti-DSS1 | 1:250 | DSS1 | 8260 |  |
|  |  |  |  | Mean: | 1.22368926 |
